# A role for the S4-domain containing protein YlmH in ribosome-associated quality control in *Bacillus subtilis*

**DOI:** 10.1101/2024.03.03.583159

**Authors:** Hiraku Takada, Helge Paternoga, Keigo Fujiwara, Jose A. Nakamoto, Esther N. Park, Lyudmila Dimitrova-Paternoga, Bertrand Beckert, Merilin Saarma, Tanel Tenson, Allen R. Buskirk, Gemma C. Atkinson, Shinobu Chiba, Daniel N. Wilson, Vasili Hauryliuk

**Author notes:** Correspondence to: Hiraku Takada, Helge Paternoga (Helge.Paternoga@ uni-hamburg.de). These authors contributed equally (Hiraku Takada, Helge Paternoga).

## Abstract

Ribosomes trapped on mRNAs during protein synthesis need to be rescued for the cell to survive. The most ubiquitous bacterial ribosome rescue pathway is trans-translation mediated by tmRNA and SmpB. Genetic inactivation of trans-translation can be lethal, unless the ribosomes are rescued by ArfA or ArfB alternative rescue factors or the ribosome-associated quality control (RQC) system, which in *B. subtilis* involves MutS2, RqcH, RqcP and Pth. Using transposon sequencing in a trans-translation-incompetent *B. subtilis* strain we identify a poorly characterized S4-domain-containing protein YlmH as a novel potential RQC factor. Cryo-EM structures reveal that YlmH binds peptidyl-tRNA-50S complexes in an position analogous to that of S4-domain-containing RqcP, and that, similarly to RqcP, YlmH can co-habit with RqcH. Consistently, we show that YlmH can assume the role of RqcP in RQC in facilitating the addition of polyalanine tails to the truncated nascent polypeptides. While in *B. subtilis* the function of YlmH is redundant with RqcP, our taxonomic analysis reveals that in multiple bacterial phyla RqcP is absent, while YlmH and RqcH are present, suggesting that in these species the YlmH plays a central role in the RQC.

## Introduction

In all cells, protein synthesis on the ribosome is a multistep process that culminates in release of the finished protein (translation termination) and separation of the ribosome from the mRNA (ribosome recycling)^1, 2^. Ribosomal stalling on damaged or truncated mRNA molecules is harmful due to sequestration of ribosomes and production of potentially cytotoxic truncated protein products. Therefore, all domains of life have evolved diverse ribosome rescue pathways that recycle such stalled ribosomal complexes^3–7^. The first bacterial ribosome rescue system to be discovered was the trans-translation system^8^. The RNA component of the system, transfer-messenger RNA (tmRNA; encoded by the gene *ssrA*, *s*mall *s*table *R*NA *A*), serves as a template encoding the C-terminal SsrA peptide tag (AANDENYALAA in *Escherichia coli*) followed by a stop codon^9^. Acting in concert with canonical bacterial translation termination machinery, the trans-translation system rescues ribosomes stalled on mRNAs lacking an in-frame stop codon, while simultaneously marking the incomplete polypeptides with a C-terminal degron tag for destruction by the ClpXP protease^10, 11^. The key element of the SsrA degron is the C-terminal Ala-Ala-COO^-^ moiety^12, 13^ that is directly recognised by the AAA+ unfoldase ClpX^10^.

Despite the trans-translation system performing an essential function in bacterial cells, in most bacteria, including *Bacillus subtilis* and *E. coli*, Δ*ssrA* strains tend to display only relatively minor phenotypes, such as increased susceptibility to antibiotics causing ribosomal miscoding^14–16^ as well as heat sensitivity^15, 17^. This can be explained by the high degree of functional redundancy of trans-translation with additional ribosome rescue systems present in these organisms. For example, alternative rescue factors (Arfs) catalyse the release of the aberrant polypeptide from the stalled ribosomal complex, but do so without adding a degron tag^18–20^. These systems either recognise the stalled ribosome and recruit canonical class-1 release factors RF1 and RF2 to cleave the stalled polypeptide (as exemplified by *E. coli* ArfA^18^ and *B. subtilis* BrfA^19^) or act directly as a peptidyl-tRNA hydrolase (as exemplified by *E. coli* ArfB^21^). Functionally, both *ssrA* and *arfA* genes are dispensable when disrupted separately in *E. coli*, but simultaneous Δ*ssrA* Δ*arfA* double-deletion results in synthetic lethality^22^, which can be suppressed by overexpression of ArfB^21, 23^. Similarly, simultaneous loss of trans-translation and BrfA-mediated ribosomal rescue in *B. subtilis* results in a severe synthetic growth defect^19^. These strong genetic interactions underscore the functional complementarity, but also redundancy, between trans-translation and Arf-mediated ribosomal rescue.

Trans-translation and Arf-mediated rescue systems do not operate on cytoplasmic eukaryotic ribosomes, making these bacterial systems attractive targets for novel antimicrobial agents^24, 25^. Instead, eukaryotes have evolved so called ribosome-associated quality control (RQC) pathways that perform a similar function^7, 26^. Recently, a homologous RQC pathway has been identified in a diverse set of bacterial species, while notably lacking in *E. coli*^7, 27^. Similarly to Arf-encoding genes, bacterial RQC genes also genetically interact with *ssrA*. Simultaneous genetic disruption of both trans-translation and RQC systems in *B. subtilis* results in a growth defect at 37°C (which is further exacerbated at elevated temperature), as well as increased sensitivity to translation-targeting antibiotics^28–30^. The mechanism of RQC has been recently extensively characterised in *B. subtilis*^28–34^. Like trans-translation, the RQC system also extends stalled polypeptides with C-terminal degron tags^32, 35^.

The RQC process can be divided into initiation (splitting of the stalled ribosome), elongation (extension of the nascent polypeptide with a degron tag) and termination (release of the tagged protein) phases. Both in eukaryotes and bacteria, ribosomal collisions serve as a substrate that is efficiently recognised by the RQC machinery^31, 36, 37^. In *B. subtilis*, rescue of stalled and collided ribosomes (disomes) is initiated by the dimeric MutS2/RqcU ATPase that splits the leading (stalled) ribosome into subunits, thereby generating a free 30S subunit and a 50S-nascent chain complex, 50S-NCC^31^. Recruitment of *B. subtilis* MutS2 to the stalled disomes is mediated by its KOW (Kyprides, Ouzounis, Woese) and Smr (small MutS-related) domains^33^. In *E. coli*, the SmrB nuclease cleaves the mRNA on the collided ribosomes^38^, which is similar to the Cue2 and NONU-1 Smr RNases that perform analogous functions in yeast^39^ and *Caenorhabditis elegans*^40^, respectively. By contrast, the *B. subtilis* MutS2 Smr domain seemingly lacks the RNase activity^33^.

Regardless, the 50S-NCCs generated by MutS2 serve as a substrate for RQC elongation which in bacteria is catalysed by two factors: RqcH and RqcP^28–30, 32^. RqcH is homologous and functionally analogous to the eukaryotic Rqc2/NEMF (Nuclear Export Mediator Factor). Yeast Rqc2 catalyses the addition of C-terminal Ala and Thr tracts, resulting in so-called ‘CAT tails’^41^, whereas mammalian CAT tails were shown to contain Ala as well as Thr, Tyr, Gly, Asp and Asn^42^. A critical step of the eukaryotic CAT tailing mechanism is the stabilization of P-site tRNA by eIF5A which enables the subsequent coordination of an A-site tRNA by Rqc2/NEMF to promote the peptide bond formation^43^. Instead of the eIF5A homologue EF-P, bacterial RQC elongation involves RqcP, a member of the S4 RNA-binding domain protein family^44^. Acting together with RqcH, RqcP catalyses the synthesis of poly-alanine tails^28, 29^. While RqcP is a close homologue of heat shock protein 15 (Hsp15). Hsp15 is an *E. coli* ribosome rescue factor that stabilizes P-tRNAs on stalled 50S subunits^45^. However, it cannot functionally substitute RqcP in *B. subtilis*^28, 30^, despite the S4 domains of the two factors forming similar interactions with the P-site tRNA of the 50S-nascent chain complexes (50S-NCC)^45^.

The final step of bacterial RQC, release of the degron-tagged protein, is the least well-understood stage of the process. In eukaryotes, a dedicated tRNA endonuclease Vms1/ANKZF1 cleaves off the CCA-3′-end of the P-site tRNA, thereby releasing the tagged polypeptide chain from the ribosome^46, 47^. By contrast, no dedicated RQC termination factor has been identified in bacteria, rather, a recent study by Svetlov and colleagues showed that the peptidyl-tRNA hydrolase (Pth) can release the poly-alanine-tagged nascent chains in *B. subtilis*^34^.

By taking advantage of the strong genetic interactions between trans-translation and other ribosome rescue systems, we have employed transposon sequencing (Tn-Seq) to identify potential novel rescue factors in *B. subtilis*. In addition to identification of the well-characterised ArfA-type rescue factor BrfA^19^, as well as the RQC elongation factors RqcP and RqcH^28–30, 32^, our Tn-Seq screen led to the identification of YlmH, a poorly characterized S4-domain-containing protein. Cryo-EM reconstructions of *ex vivo* YlmH-ribosome reveal that YlmH interacts with 50S-NCCs. YlmH has a binding site analogous to RqcP and, similarly to RqcP, can co-habit 50S-NCC with RqcH, which is strongly indicative of YlmH being a RQC factor. Using *in vivo* reporter assays, we demonstrate that overexpression of YlmH in RQC-incompetent Δ*rqcP B. subtilis* stain can, indeed, restore the functionality of the RQC-mediated polyalanine tailing, indicating that YlmH is functionally redundant with RqcP. Finally, our phylogenic analysis indicates that some YlmH-containing bacteria do not encode RqcP, suggesting that in these phyla YlmH is likely to play a central role in RQC-mediated polyalanine tailing.

## Results

### *B. subtilis* S4-domain containing protein YlmH is a potential RQC factor

We have mounted a search for as-yet-unidentified *B. subtilis* factors involved in ribosomal rescue by performing a Tn-Seq experiment with the Δ*ssrA* (VHB257) strain as well as an isogenic parental 168 wild-type *B. subtilis* strain. The relative difference in the transposon insertion frequency between the two libraries reflects the difference in the insertion’s effect on bacterial fitness in the two genetic backgrounds^48^. The screen reliably identified genes encoding known ribosome rescue factors, including the release factor-dependent alternative ribosome rescue factor *brfA*^19^ as well as the RQC factors *rqcH* and *rqcP* (**Fig. 1a**). Furthermore, a decreased frequency of insertion was observed for the *ylmH* ORF, indicative of YlmH also playing a role in ribosomal rescue. The experimentally unexplored *ylmH* gene is part of the *ylm* operon that is located downstream of the *dcw* (*division and cell wall*)^49^ gene cluster^50^. Importantly, just as RqcP and Hsp15, YlmH possesses an S4 RNA-binding domain^44^, raising a possibility of a translation-related function.

**Fig. 1.**
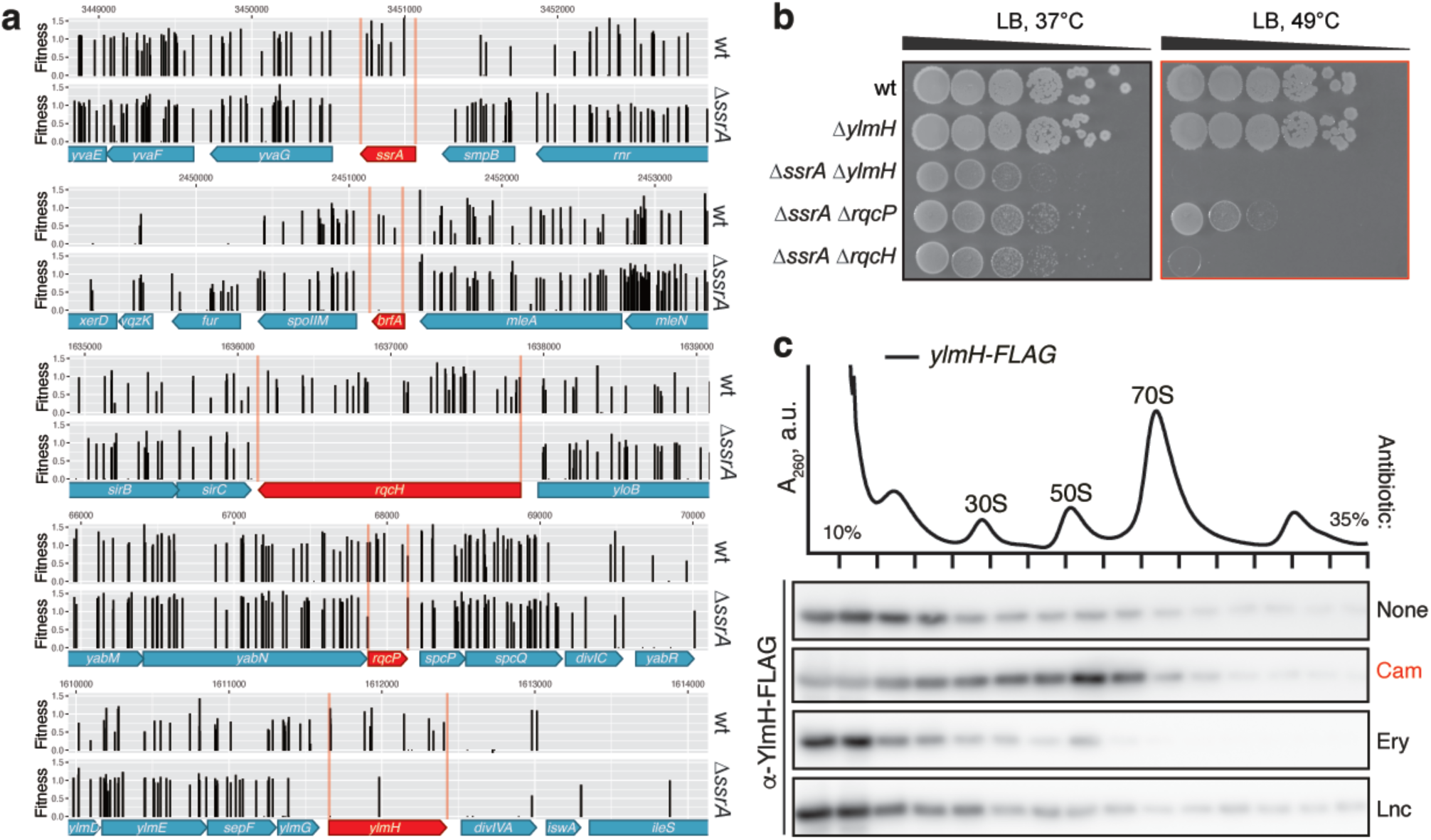
*B. subtilis* YlmH is implicated in Ribosome Quality Control. (**a**) Deletion of *ylmH* in the Δ*ssrA* background is associated with a fitness loss. Transposon libraries were generated in 168 wild-type and Δ*ssrA* (VHB257) *B. subtilis* strains. Sites of transposon insertion were identified by Illumina NextSeq sequencing and mapped onto the *B. subtilis* 168 reference genome. Independent insertions in the genomic region encompassing *ssrA*, *brfA*, *rqcH*, *rqcP* and *ylmH* genes as well as insertion’s fitness effect are shown. (**b**) Deletions of *ylmH*, *rqcH* and *rqcP* in the Δ*ssrA* background cause synthetic growth defect. 10-fold serial dilutions were spotted onto LB agar plates and incubated for 18 hours at either 37°C (left; optimal temperature) or 49°C (right; mild heat shock). (**c**) Chloramphenicol treatment promotes YlmH recruitment to 50S. C-terminally FLAG-tagged YlmH (YlmH-FLAG; C-terminal FLAG tag connected via GS5 linker) was ectopically expressed in Δ*ylmH* background under the control of P*_hy-spank_* promotor (BCHT931). YlmH expression was induced by 30 μM IPTG. Following a 20-min antibiotic treatment either chloramphenicol (Cam, 5 μg/mL), erythromycin (Ery, 1 μg/mL) or lincomycin (Lnc, 80 μg/mL), cellular lysates were resolved on sucrose gradients and fractions probed with anti-FLAG antibodies.

We genetically probed the involvement of YlmH in ribosomal rescue by testing for a synthetic growth defect upon *ylmH* disruption with a spectinomycin resistant gene cassette (*spcR*) in the Δ*ssrA* background. The growth phenotypes associated with Δ*ylmH* and Δ*ylmH* Δ*ssrA* mutations were compared with that associated with Δ*rqcH* and Δ*rqcP* deletions. While the Δ*rqcH* loss is not associated with a strong phenotype in *B. subtilis* under non-stress conditions, the Δ*ssrA* Δ*rqcH* double-deletion strain displays a growth defect at 37°C and at elevated temperature (49°C, mild heat shock conditions) the growth is abrogated completely^32^ (**Fig. 1b**). The Δ*ssrA* Δ*rqcP* double-deletion results in a milder synthetic growth defect, with the effect being more pronounced at 49°C. While the Δ*ylmH B. subtilis* strain displays no growth defect at either 37°C or 49°C, the Δ*ssrA* Δ*ylmH* double-deletion phenocopies the Δ*ssrA* Δ*rqcH* strain, providing further support for the involvement of YlmH in RQC.

*B. subtilis* RQC factors are recruited to the 50S subunit upon challenge with translation-targeting antibiotics^29, 30, 32^. Therefore, we were interested to assess whether YlmH also co-migrates with the 50S subunit under antibiotic stress. To do so, we used sucrose gradient centrifugation and immunoblotting to monitor the potential recruitment of C-terminally FLAG-tagged YlmH to ribosomal particles under different antibiotic stress conditions (**Fig. 1c**). In the absence of antibiotics, YlmH appeared to be predominantly located at the top of the gradient and only minimal association with ribosomal particles was observed. By contrast, in the presence of chloramphenicol YlmH appeared to migrate with the 50S subunit, as observed for previously for RqcP and RqcH^28–30, 32^. Moreover, this association appeared to be specific for chloramphenicol stress, since negligible association with ribosomal particles was observed with other antibiotic stresses, such as the presence of erythromycin or lincomycin (**Fig. 1c**). It is noteworthy that chloramphenicol is a context-dependent elongation inhibitor that forms extensive interactions with the alanine residue at the -1 position of the nascent chain^51–53^. As bacterial RQC is mediated by formation of C-terminal poly-alanine tails^29, 30, 32^, specific stabilisation of YlmH by chloramphenicol could be indicative of the factor’s physical association with poly-alanine-containing ribosomal complexes.

### Cryo-EM structures of *ex vivo* YlmH-50S-NCC

Given the apparent association of YlmH with 50S particles (**Fig. 1c**), we employed affinity chromatography from a *B. subtilis* Δ*ylmH* strain expressing YlmH with a C-terminal FLAG-tag (YlmH-FLAG) to isolate native YlmH-ribosome complexes for structural analysis. We have earlier employed this experimental strategy for other ribosome binding factors^45^ including RqcH and RqcP^28, 30^. SDS-PAGE analysis of the pulldown samples revealed that 50S subunit ribosomal proteins co-purified with YlmH-FLAG (**Fig. 2a**), which was confirmed by mass spectrometry (see **Methods**). Together with the sucrose density gradient analysis (**Fig. 1c**), these findings indicate that YlmH indeed interacts with 50S subunits, analogous to other RQC factors such as RqcP and RqcH^28–30, 32^. The YlmH-50S complexes were then subjected to single particle cryo-EM analysis (see **Methods**). While the initial 3D classification indicated that the majority of ribosomal particles were 50S subunits, little density for additional ligands (tRNAs) or factors (YlmH) was observed (**Supplementary Fig. 1a-c**). Therefore, 3D classification was performed using a mask encompassing the A-, P- and E-sites on the 50S subunit (**Supplementary Fig. 1d**). This yielded two classes with an additional density adjacent to 23S rRNA helix 69 (H69) that we assigned to YlmH. In class 1 (154,237 particles, 6.4%), YlmH was well-resolved and accompanied by additional density for a P-site tRNA, whereas in the major class 2 (1,585,755 particles, 66.3%) YlmH was less resolved and there was no apparent density for any tRNAs (**Supplementary Fig. 1e**). The minor class 3 (61,275 particles, 2.7%) contained 70S particles lacking any additional density for YlmH, so this class was not considered further. Given the well-defined density for YlmH and the presence of P-site tRNA, class 1 was subjected to further rounds of 3D classification, eventually yielding a well-defined YlmH-50S state (“State 1” with 66,113 particles, 2.7% of the total) (**Supplementary Fig. 1f-g**) containing YlmH and P-site tRNA (**Fig. 2b**). State 1 could be refined to generate a structure of the YlmH-50S complex with an average resolution of 2.0 Å (**Table 1** and **Supplementary Fig. 2**).

**Fig. 2.**
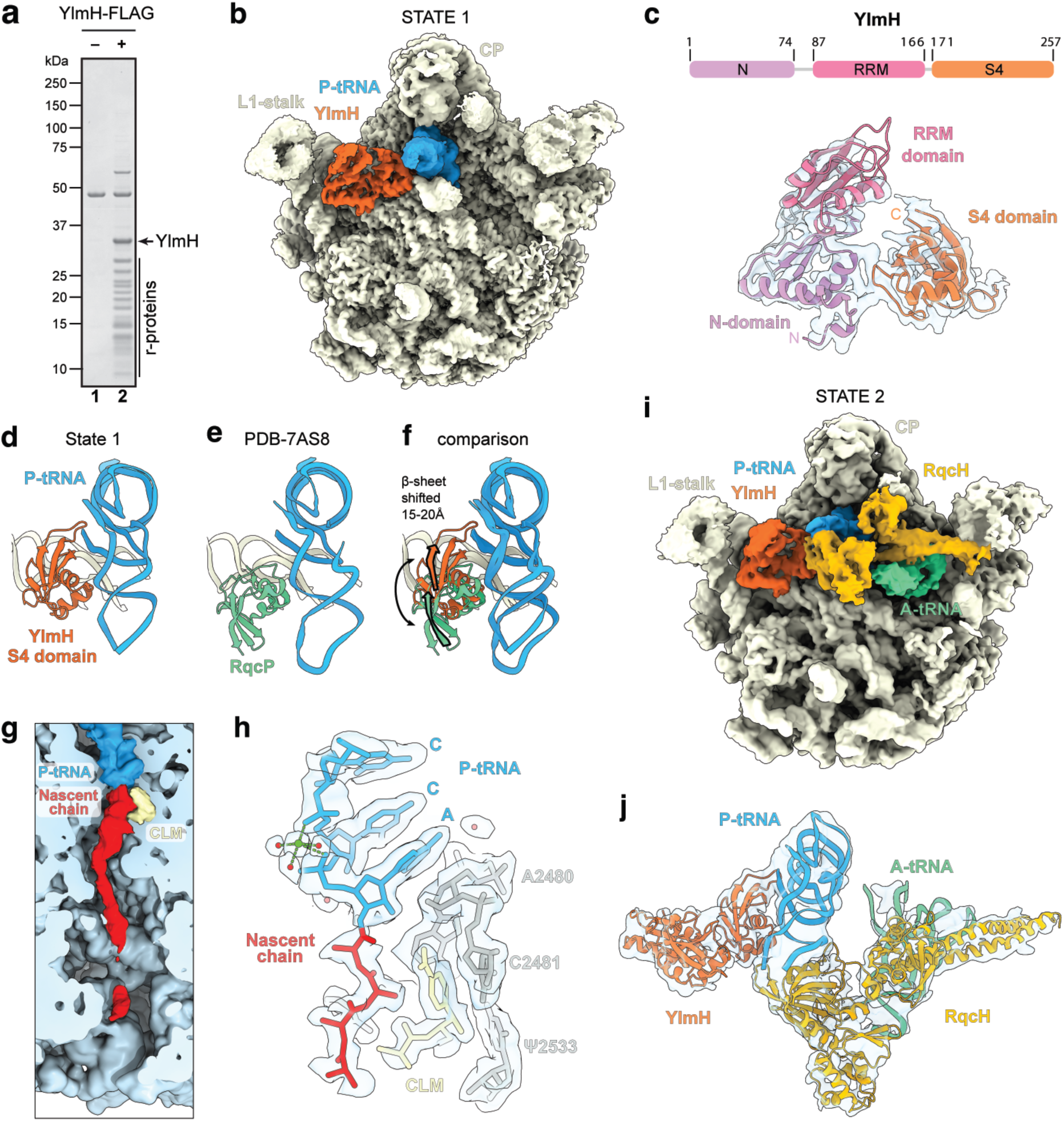
Cryo-EM structure of YlmH-50S complexes. (**a**) SDS-PAGE analysis of control (-) and YlmH-FLAG coimmunoprecipitations revealing the presence of ribosomal proteins of the large subunit. (**b**) Cryo-EM reconstruction of State 1 consisting of YlmH (orange), P-tRNA (blue) bound to the large 50S subunit (yellow) with landmarks L1-stalk and central protuberance (CP) highlighted. (**c**) Schematic of the domain structure of YlmH with N-(purple), central RRM (red) and S4 (orange) domains indicated. Lower panel shows the isolated cryo-EM density (transparent grey) for YlmH with molecular model (colored by domain). (**d-f**) comparison of the relative binding position of (**d**) YlmH (orange), (**e**) RqcP (green, PDB ID 7AS8)^28^ to P-tRNA (blue) and (f) superimposition of (d) and (e) with arrows indicating the shift in position of the β-sheet of YlmH compared to RqcP. Alignments were made on the basis of the 23S rRNA. (**g**) Transverse section and zoom of the polypeptide exit tunnel of State 1 revealing the presence of a polypeptide chain (red) attached to the P-site tRNA (blue) and chloramphenicol (yellow) bound in the A-site of the 50S subunit (light blue). (**h**) Isolated cryo-EM density (transparent grey) of the CCA-end of the P-tRNA (blue), chloramphenicol (yellow) and nascent chain (red, with polyalanine model) with selected 23S rRNA nucleotides (grey). (**i**) Cryo-EM reconstruction of State 2 consisting of YlmH (orange), P-tRNA (blue), A-tRNA (green) and RqcH (mustard) bound to the large 50S subunit (yellow) with landmarks L1-stalk and central protuberance (CP) highlighted. (**j**) Isolated cryo-EM density (transparent grey) and fitted molecular models for YlmH (orange), P-tRNA (blue), A-tRNA (green) and RqcH (mustard) from State 1.

**Table 1.** Cryo-EM data collection, refinement and validation statistics.

| **Model** | **YlmH-50S (P-tRNA)** | **YlmH-RqcH-50S (A- & P-tRNA)** |
| --- | --- | --- |
| **EMDB ID** | EMD-19638 | EMD-19641 |
| **PDB ID** | 8S1P | 8S1U |
| **Data collection and processing** |  |  |
| Magnification (×) | 96,000 | 96,000 |
| Acceleration voltage (kV) | 300 | 300 |
| Electron fluence (e<sup>-</sup>/Å<sup>2</sup>) | 40 | 40 |
| Defocus range (μm) | -0.4 to -0.9 | -0.4 to -0.9 |
| Pixel size (Å) | 0.80 | 0.80 |
| Initial particles | 2,391,525 | 2,391,525 |
| Final particles | 66,113 | 3,770 |
| Average resolution (Å) (FSC threshold 0.143) | 2.0 | 3.4 |
| **Model composition** |  |  |
| Initial model used (PDB code) | 6HA8 | 6HA8 |
| Atoms | 91,097 | 93,269 |
| Protein residues | 3,270 | 3,784 |
| RNA bases | 2,914 | 3,036 |
| **Refinement** |  |  |
| Map CC around atoms | 0.77 | 0.75 |
| Map CC whole unit cell | 0.61 | 0.57 |
| Map sharpening B factor (Å<sup>2</sup>) | -1.04 | -38.21 |
| **R.M.S. deviations** |  |  |
| Bond lengths (Å) | 0.008 | 0.010 |
| Bond angles (°) | 1.528 | 2.061 |
| **Validation** |  |  |
| MolProbity score | 0.85 | 1.63 |
| Clash score | 0.39 | 2.41 |
| Poor rotamers (%) | 0.40 | 1.76 |
| **Ramachandran statistics** |  |  |
| Favoured (%) | 96.66 | 93.65 |
| Allowed (%) | 3.15 | 5.94 |
| Outlier (%) | 0.19 | 0.40 |
| Ramachandran Z-score | -1.21 | -2.69 |

The excellent quality of cryo-EM map density for State 1, particularly within the core of the 50S subunit (**Supplementary Fig. 2**), enabled the identification of nine distinct modifications within the 23S rRNA (**Supplementary Fig. 3**). This encompassed six modifications that have been visualized in the 23S rRNA of the *E. coli* 70S ribosome^54^, namely, the four base methylations 5MU794 (Ec747), 5MU1968 (Ec1939), 2MG2474 (Ec2445), 2MA2532 (Ec2503); one ribose methylation at OMG2280 (Ec2251) and the dihydrouridine DHU2478 (Ec2449). In addition, we observe three modifications that have not been observed previously in *E. coli* or *T. thermophilus* 70S ribosomes^54–56^, including two base modifications, namely, 5MU620 and 7MG2603, as well as one ribose methylation, OMG2582 (**Supplementary Fig. 3**). G2582 is present in the A-loop at the peptidyltransferase center and the 2′-O-methylation was shown recently to be mediated by the methyltransferase RlmP (formerly YsgA)^57^. It is interesting that G2582 (Ec2553) neighbors U2581 (EcU2552), which is 2′-O-methylated in other organisms, such as *E. coli*, that lack the G2582 (Ec2553) modification, suggesting the presence, rather than the exact position, of a modification in the A-loop is important for function.

### YlmH binds analogous to RqcP on the 50S subunit

The additional density in State 1 could be unambiguously assigned to YlmH (**Fig. 2b,c**). Unlike *B. subtilis* RqcP that is 86 aa and comprises only a single S4 domain, *B. subtilis* YlmH is 257 aa and comprises three domains (**Fig. 2c**). In addition to the C-terminal S4 domain (residues 171-257), YlmH contains an N-terminal domain (NTD, residues 1-74) as well as central domain (residues 87-166). The central domain of YlmH contains an RNA recognition motif (RRM) with high structural homology to RRMs found in DEAD-box helicases, such *E. coli* DbpA^58, 59^ (**Supplementary Fig. 4a,b**). A Dali search reveals that the NTD exhibits the highest structural similarity to the R3H domain found in an unpublished structure (PDB ID 3GKU) of a putative RNA-binding protein from *Clostridium symbiosum*, which is homologous to the YidC1-associated Jag/KhpB protein (BSU_41030) in *B. subtilis* (**Supplementary Fig. 4c**). While the NTD and C-terminal S4 domain were generally well-resolved (2.5-3.5 Å; **Supplementary Fig. 3**), enabling sidechains to be modelled with high confidence for the regions interacting with the rRNA, the central RRM domain linking the N- and C-terminal domains was less well-resolved (3.5-5 Å; **Supplementary Fig. 2**), enabling only rigid body fitting of an alpha-fold homology model^60^. Nevertheless, the conformation and arrangement of the NTD and RRM domains observed for YlmH in State 1 appears very similar to that seen in the unpublished X-ray structure (PDB ID 2FPH) of *Streptococcus pneumoniae* YlmH lacking the S4 domain and determined in the absence of the ribosome (**Supplementary Fig. 4d**). As expected, the C-terminal S4 domain of YlmH is located similarly on the 50S subunit to RqcP^28, 29^, where it interacts with both the P-site tRNA and H69 of the 23S rRNA (**Fig. 2d,e**). This is particularly evident for the two α-helices within the S4 domain of YlmH that are oriented to establish multiple interactions with H69, as reported previously for RqcP^28, 29^. By contrast, the five-stranded Δ-sheet in the S4 domain is shifted by 15-20 Å, when comparing the relative positions between YlmH and RqcP (**Fig. 2f**). As a result, YlmH establishes additional interactions with the P-site tRNA from residues located in the loop between Δ-strands Δ4 and Δ5. While this interaction is not possible for RqcP, we note that RqcP contains a short α-helix (residues 68-73), which is absent in the S4 domain of YlmH, that contacts the anticodon stem loop of the P-site tRNA^28, 29^.

In State 1, we observed additional density within the polypeptide exit tunnel, which is better resolved in the upper region of the tunnel and became disordered, probably due to flexibility, towards the tunnel exit (**Fig. 2g**). The density for the nascent chain at the PTC is consistent with four alanine residues extending from the CCA-end of the P-site tRNA (**Fig. 2h**). We also observed density for the antibiotic chloramphenicol bound within the A-site of the PTC (**Fig. 2g,h**). Although initially unexpected, chloramphenicol was used to select for the YlmH-FLAG expression plasmid and was therefore present in the growth medium upon lysis and could also be potentially present at low concentrations within the cell during growth (see **Methods**). Thus, chloramphenicol could bind to the YlmH-50S complexes during purification, if it was not already bound to the complexes within the cell. As mentioned above, chloramphenicol is a context-specific inhibitor that binds stably to the A-site of the PTC when alanine (and to a lesser extent serine or threonine) is present in the -1 position of the nascent chain^51^, providing an explanation for the observed co-migration of YlmH with 50S subunits on sucrose gradients in the presence of this antibiotic (**Fig. 1c**). We also note that the interaction between the -1 alanine and chloramphenicol observed in the State 1 is near-identical to that observed in complexes programmed with alanine in the -1 position^53^ (**Supplementary Fig. 4e,f**). Thus, while we cannot exclude that the nascent chain represents a mixture of sequences (where the sidechains are averaged away and only the backbone density remains), we favor the interpretation that YlmH is bound to 50S subunits bearing polyalanine tails, as expected for a factor involved in RQC^28–30, 32^.

The strong genetic and structural evidence for the involvement of YlmH in poly-alanine tailing during RQC, prompted us to further sub-sort our YlmH:P-tRNA:50S complexes in search of additional functional states. This led to the identification of a small population of 50S subunits (0.2%), which we termed State 2 (**Supplementary Fig. 2h,i**), that contained in addition to YlmH and P-tRNA, also A-tRNA and RqcH (**Fig. 2i**). Despite the limited particle number (3,770 particles), we were nevertheless able to refine State 2 to an average resolution of 3.4 Å (**Table 1** and **Supplementary Fig. 5**). While the local resolution of the core of the 50S subunit reached towards 3.0 Å, the ligands were significantly less well-resolved, presumably due to flexibility (**Supplementary Fig. 5**). Nevertheless, the resolution (5-10 Å) was sufficient to dock homology models for YlmH, RqcH as well as A- and P-tRNAs (**Fig. 2j**), revealing a functional state with remarkable similarity to that reported previously for an RqcP:RqcH:50S-NC complex, which also contained both A- and P-site tRNAs^29^ (**Supplementary Fig. 4g,h**). This further supports the notion that YlmH can participate in polyalanine tailing during RQC in *B. subtilis*.

### Importance of both N- and CTD of YlmH interaction with the 50S subunit

Conserved Arg2 and Arg16 residues within the N-terminal region of the S4 domain of RqcP mediate essential interactions with 23S rRNA, to the extent that alanine substitutions of either residue abrogate the factor’s functionality and phenocopy the Δ*rqcP* strain^30^. These two arginine residues are also conserved in YlmH suggesting an analogous functional importance (**Fig. 3a**). Indeed, YlmH forms extensive contacts with H69 of the 23S rRNA through both the YlmH-specific NTD region (amino acids 1-74) and the S4 domain (amino acids 171-257) (**Fig. 3b**), which it shares with RqcP. The conservation of the functional residues in the S4 domain and extensive contacts observed in cryo-EM reconstructions raised the question as to whether the rRNA contacts provided by the additional YlmH-specific CTD are also essential for functionality. To address this question, we first tested the functional importance of Arg182 (corresponding to Arg2 of *B. subtilis* RqcP), Arg195 (which could be the functional equivalent of Arg16 of *B. subtilis* RqcP) and Lys197 (used as a control; does not form contacts with rRNA) of YlmH in phenotypic assays by assessing the heat sensitivity of *B. subtilis* expressing mutant YlmH variants in the Δ*ssrA* background. As expected, alanine substitutions of R182 and R195 phenocopy the Δ*ylmH* deletion, whereas the K197A substitution had no effect on YlmH functionality, since the *ylmH K197A* Δ*ssrA* strain grew similarly to isogenic Δ*ssrA* at both 37°C and 49°C (**Fig. 3c**). This suggests that the conserved Arg182 and R195 that mediate the interactions of YlmH with H69 of the 23S rRNA (**Fig. 3b**) are likely to be critical for factor’s functionality, whereas the non-conserved Lys197 is unimportant.

**Fig. 3.**
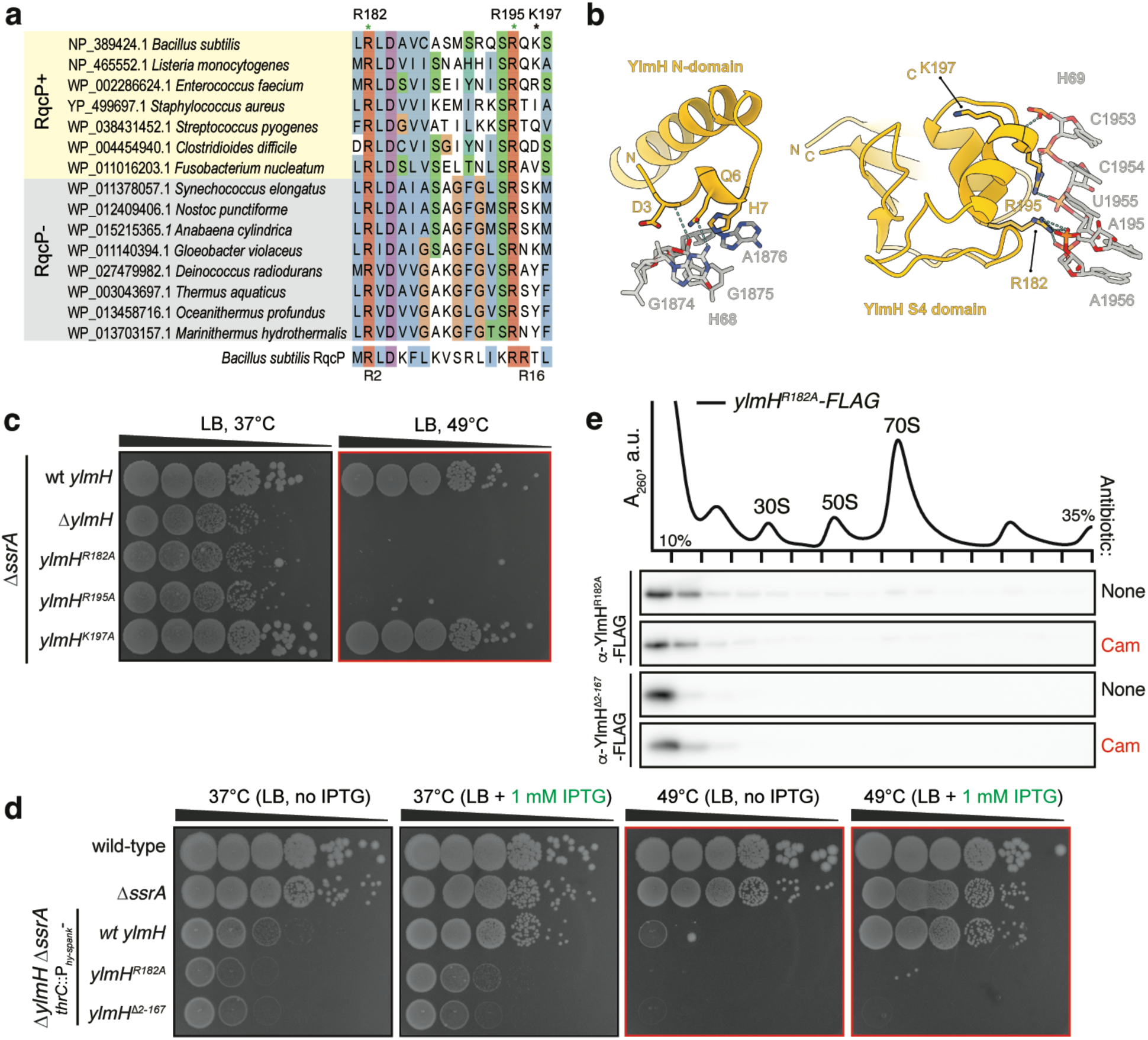
Contacts with H69 23S rRNA and P-site tRNA are essential for YlmH functionality. (**a**) Sequence alignment of YlmH homologs from representative RqcP-encoding and RqcP-lacking bacterial species. (**b**) Interaction network between 23S rRNA H69 and YlmH. R182 and R195 are predicted to interact with H69, whereas K197 is not. (**c**) Disruption of YlmH contacts with H69 abrogates YlmH functionality *in vivo*. Synthetic growth defects *ylmH* mutant variants in the Δ*ssrA* background of were compared with *ylmH* deletion. 10-fold serial dilutions were spotted onto LB agar plates and incubated for 18 hours at 37°C (left) or 49°C (right). (**d**) R182A substitution and N-terminal truncation (Δ2-187) of *ylmH* abrogate its functionality in *in vivo.* Wild-type and mutant *ylmH* variants were expressed in Δ*ylmH* Δ*ssrA* background under the control of P*_hy-spank_* promotor. 10-fold serial dilutions were spotted onto LB agar plates with or without 1 mM IPTG, and incubated for 18 hours at either 37°C or 49°C. (**e**) *R182A* and Δ2-187 YlmH variants compromised in H69 binding are not recruited to the 50S upon chloramphenicol challenge. Sucrose gradient sedimentation and anti-FLAG immunoblotting of YlmH^R182A^-FLAG (BCHT1327) and YlmH^Δ2-187^-FLAG (BCHT932) expressed in Δ*ylmH* background, with or without 5 μg/mL chloramphenicol (Cam) treatment.

Next, we tested the essentiality of the N-terminal domain of YlmH (amino acid residues 1-166) which is absent in RqcP. To do so, we expressed a YlmH variant lacking the N-domain (*ylmH* Δ2-167) in the Δ*ylmH* Δ*ssrA* background under the control of IPTG-inducible P*_hy−spank_* promoter^61^ (**Fig. 3d**). As a positive control, we overexpressed the wild-type full-length YlmH, and as a negative control, we expressed the inactive YlmH R182A variant. As expected, the IPTG-induced expression of wild-type YlmH but not the *ylmH* Δ2-167 or *ylmH R182A* mutant variants suppressed the relatively mild growth defect of Δ*ylmH* Δ*ssrA B. subtilis* at 37°C. Similarly, under the mild heat shock conditions (49°C), expression of neither of the mutant variants could overcome the synthetic lethality, suggesting that both *ylmH R182A* and *ylmH* Δ2-167 are non-functional. Collectively, these results suggest that the N-terminal domain of YlmH is also essential for factors’ functionality.

Finally, we used sucrose gradient centrifugation and immunoblotting experiments to probe the recruitment of the C-terminally FLAG-tagged YlmH Δ2-167 and YlmH R182A variants to 50S subunits upon a chloramphenicol challenge (**Fig. 3e**). In contrast to the wild-type YlmH-FLAG, which co-migrates with 50S subunits in the presence of chloramphenicol (**Fig. 1c**), neither the R182A-substituted YlmH variant nor the N-terminally truncated YlmH Δ2-167 variant were detected in the 50S fraction (**Fig. 3e**). This suggests that the inability of these variants to rescue the growth phenotype of the Δ*ssrA* Δ*rqcH* double deletion strain (**Fig. 3d**) most likely stems from the inability of these variants to stably interact with the 50S subunit. Thus, we conclude that in addition to interaction between the S4 domain of YlmH and H69, the N-terminal domain also contributes to the stable binding of YlmH to the 50S subunit. This interpretation is consistent with the direct interactions observed between N-terminal residues, such as Asp3 and Gln6, and 23S rRNA nucleotides located within H68 (**Fig. 3b**).

### YlmH overexpression can rescue the growth defect of the Δ*ssrA* Δ*rqcH* strain

To address whether YlmH is functionally interchangeable with RqcP, we initially investigated whether overexpression of YlmH can rescue the growth phenotype observed when the Δ*ssrA* Δ*rqcP* double-deletion strain is subjected to heat stress at 49°C (**Fig. 4a**). We also performed the analogous experiment using the Δ*ssrA* Δ*rqcH* double-deletion strain, although we did not expect YlmH to able to functionally substitute for RqcH. Indeed, no restoration of the growth defect at 49°C was observed upon P*_hy-spank_*-driven expression of YlmH in the Δ*ssrA* Δ*rqcH* strain (**Fig. 4a**). Conversely, expression of YlmH restored the growth defect of the Δ*ssrA* Δ*rqcP* double-deletion strain. This result is reminiscent of ArfB overexpression being able to suppress the synthetic lethality of Δ*ssrA* Δ*arfA* double deletion in *E. coli*^21, 23^. Finally, to test the specificity of the effect of YlmH overexpression, we probed whether overexpression of other S4 domain-containing *B. subtilis* proteins can suppress the temperature sensitivity of the Δ*ssrA* Δ*rqcP* strain. The P*_hy-spank_*-driven expression of none of the tested genes (*rluB*, *rpsD*, *tyrS*, *tyrZ*, *yaaA*, *yhcT*, *yjbO*, *ylyB*, *yqxC* and *ytzG*) suppressed the heat-sensitivity phenotype of Δ*ssrA* Δ*rqcP B. subtilis* (**Supplementary Fig. 6**), suggesting that the rescue effect of YlmH overexpression is specific.

**Fig. 4.**
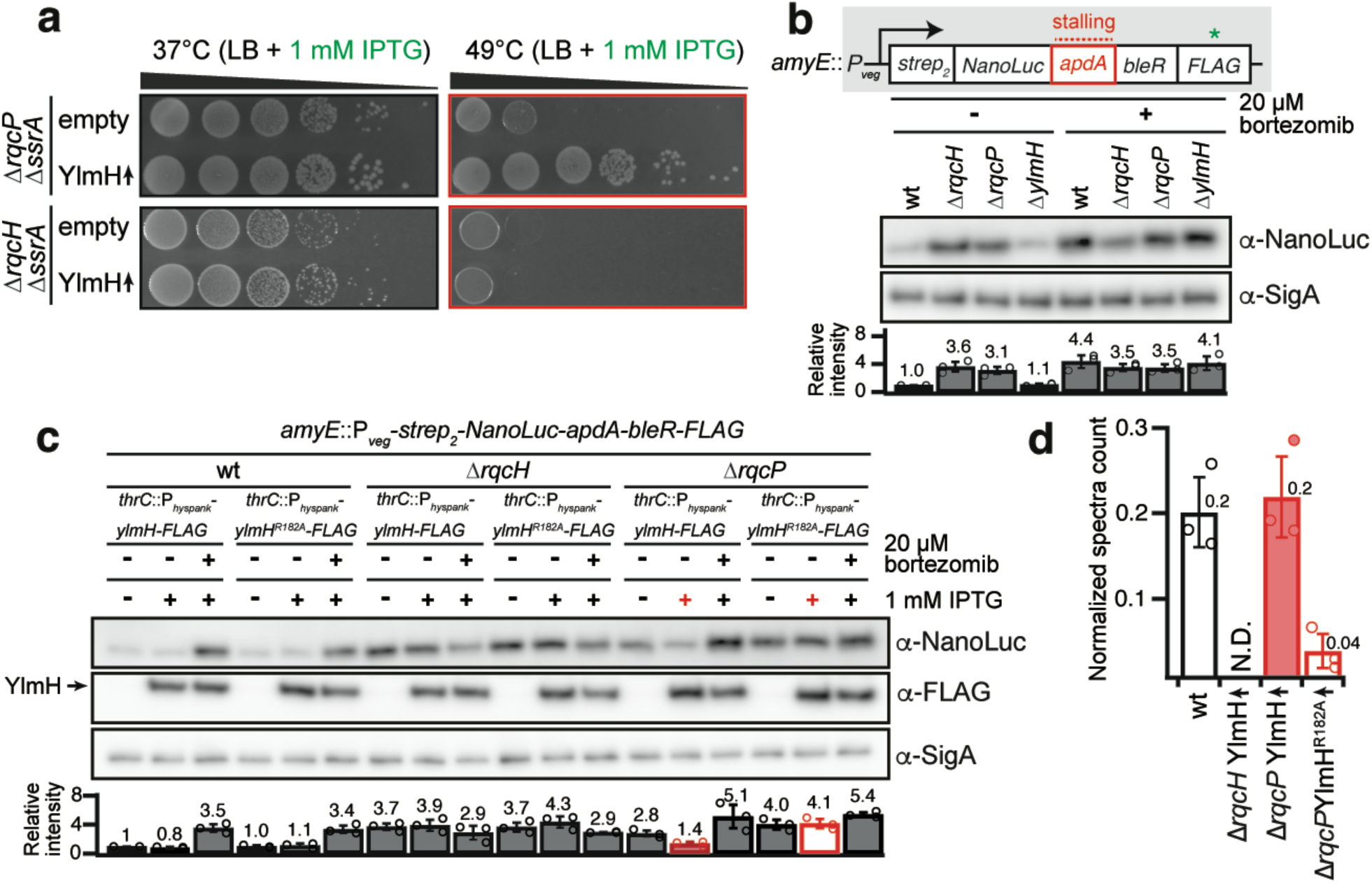
YlmH can substitute RqcP, but not RqcH, during poly-alanine tailing. (**a**) Ectopic expression of *ylmH* suppresses the growth defect of the Δ*rqcP* Δ*ssrA* but not of the Δ*rqcH* Δ*ssrA* strain. Strains with or without IPTG-inducible construct in either Δ*rqcH* Δ*ssrA* (VHB788, VHB787) or Δ*rqcP* Δ*ssrA* (VHB789, VHB600) background were serially diluted, spotted onto LB agar plates without 1 mM IPTG and incubated for 18 hours at either 37°C or 49°C. (**b**) Δ*rqcH* and Δ*rqcP* but not Δ*ylmH* strains are defective in RQC-mediated poly-alanine tagging. A 31-residue ApdA stalling motif from *A. japonica* was inserted between N-terminally twin-strep tagged NanoLuc and C-terminally FLAG tagged bleomycin resistance protein (BleR). The reporter is expressed under the control of constitutive P*_veg_* prompter in wild-type (BCHT1294), Δ*rqcH* (BCHT1295), Δ*rqcP* (BCHT1296) and Δ*ylmH* (BCHT1297) strains after treatment with 20 µM ClpP inhibitor bortezomib and detected with anti-NanoLuc antibodies. The SigA protein was used as a loading control. For quantification the anti-NanoLuc signal was normalized to the internal anti-SigA control, and results of three independent biological replicates are shown as mean ±SD. (**c**) YlmH but not inactive YlmH^R182A^ can substitute RqcP in poly-alanine tailing reaction. Wild-type *ylmH* or *ylmH^R^*^182^*^A^* were ectopically expressed in wild-type (BCHT1133, BCHT1134; *left*), Δ*rqcH* (BCHT1135, BCHT1136; *middle*) and Δ*rqcP* (BCHT1137, BCHT1138; *right*) strains, and accumulation of the NanoLuc reporter was assessed with or without 20 µM bortezomib treatment. Data quantification was done as for panel A. The arrow indicates the position of C-terminally FLAG-tagged wildtype YlmH and YlmH^R182A^. (**d**) Overexpression of wild-type but not *R182A* YlmH rescues poly-alanine tailing in Δ*rqcP* but not Δ*rqcH* strain. The Ala-tail at the stall site of the reporter in wild-type, Δ*rqcH*, and Δ*rqcP* strains expressing either wild-type *ylmH* or *ylmH^R^*^182^*^A^* was detected by LC-MS/MS. The peptide counts are normalized to a different peptide in the reporter protein.

### YlmH overexpression rescues poly-alanine tailing in *ΔrqcP* cells

Taken together, our results indicate that YlmH can functionally substitute for RqcP, and therefore YlmH may also be involved in the elongation phase of RQC. To test this hypothesis, we used the recently developed RQC reporter^33^ expressed under the control of constitutive P*_veg_* promoter^62^. The reporter encodes a N-terminally twin-Strep-tagged NanoLuc luciferase conjoined via a 31-residue ApdA stalling motif from *Amycolatopsis japonica*^63^ with the C-terminally FLAG-tagged bleomycin resistance protein (BleR). A ClpP inhibitor bortezomib^64^ can be used to stabilise the poly-alanine-tailed RQC products^32^. As we have shown earlier, the α-NanoLuc signal increases in the Δ*rqcH* strain due to the absence of efficient tagging and targeting of the stalled product for ClpXP-mediated degradation^33^. The reporter was inserted into the *amyE* locus coding for a nonessential α-amylase (*amyE*::P*_veg_*-*strep*_2_-*NanoLuc*-*apdA*-*bleR*-*FLAG*) of either the wild-type 168 *B. subtilis* strain or *B. subtilis* strains lacking RqcH (Δ*rqcH*), RqcP (Δ*rqcP*) or YlmH (Δ*ylmH*). The cultures were grown in the presence or absence of 20 μM bortezomib and the expression of the reporter protein was visualized on immunoblots with α-NanoLuc antibodies (**Fig. 4b**). To compare the effects across conditions and strains, NanoLuc levels were quantified, and the signal in the untreated wild-type strain was set as 1. In the wild-type strain, the α-NanoLuc signal increased 4.4-fold for the bortezomib-treated culture (**Fig. 4b**), reflecting the failure of ClpP-driven degradation of the poly-alanine-tailed nascent polypeptide. Consistent with the lack of ribosome rescue and poly-alanine tailing in the RQC-deficient Δ*rqcP* and Δ*rqcH B. subtilis*, in these strains the α-NanoLuc signal is high (3.1-3.6-fold increase over the wild-type level) and bortezomib-insensitive (3.5-fold wild-type level), By contrast, the Δ*ylmH* strain behaves similarly to wild-type. In the absence of bortezomib, the α-NanoLuc signal was 1.1 of the wild-type level, and it increased upon the bortezomib treatment to 4.1-fold over the untreated level. These results suggest that in the Δ*ylmH* strain efficient poly-alanine tailing is mediated by canonical RqcH and RqcP factors.

As our genetic experiments suggested that YlmH can compensate for the loss RqcP but not RqcH (**Fig. 4a**), we next employed the NanoLuc reporter to test the effects of YlmH overexpression in wild-type as well as in the Δ*rqcP* and Δ*rqcH* strains, both in the presence and absence of bortezomib (**Fig. 4c**). The inactive R182A-substituted variant of YlmH was used as a negative control, and YlmH was C-terminally FLAG-tagged to monitor its inducible expression. For the wild-type *B. subtilis* strain, the expression of neither the wild-type YlmH, nor the R182A-substituted YlmH-FLAG variant, had any effect on the weak α-NanoLuc signal in the absence of bortezomib (**Fig. 4c**, *left*). Furthermore, expression of neither of the factors had any effect on the increase (3.4-3.5-fold) in the α-NanoLuc signal observed in the presence of bortezomib. This indicates that YlmH expression has no dominant negative effect on the canonical RQC pathway. In the Δ*rqcH* strain, the elevated NanoLuc signal was insensitive to both bortezomib and YlmH expression, indicative of the absence of poly-alanine degron tailing (**Fig. 4c**, *middle*); as expected, YlmH cannot functionally compensate for loss of RqcH. In contrast, expression of YlmH in the Δ*rqcP* strain in the absence of bortezomib led to a decrease (1.4-fold) in the α-NanoLuc signal to near-wild-type levels, and in the presence of bortezomib the signal increased 4.1-fold, (**Fig. 4c**, *right*). Importantly, this effect was specific, since expression of the inactive R182A-substituted YlmH-FLAG in the Δ*rqcP* strain had no effect on the NanoLuc signal, which remained equally elevated (4.1-5.1-fold) regardless the absence or presence of bortezomib. Collectively, these results suggest that, when overexpressed, YlmH can substitute for RqcP in the poly-alanine tailing reaction.

Finally, to directly assess the effect of YlmH on poly-alanine tailing, we used mass spectrometry to quantify the levels of alanine-tailing on the ApdA reporter (**Fig. 4c**). We expressed the reporter in the wild-type strain as well as *ΔrqcH* and *ΔrqcP* knockout strains overexpressing YlmH, either just the wild-type (in *ΔrqcH* background) or both wild-type and R182A-substituted YlmH variants (in *ΔrqcP* background). The strains were grown in the presence of bortezomib to prevent the degradation of truncated or Ala-tagged proteins, the reporter was immunoprecipitated via its N-terminal Strep_2_-tag, the eluted proteins were digested with lysyl endopeptidase (LysC) and the resulting peptides were analyzed with liquid chromatography coupled tandem mass spectrometry (LC-MS/MS). Ribosomes stall on the ApdA sequence at the RAPP motif with the first Pro codon in the P site and the second Pro codon in the A site^63^, and our previous work has shown that poly-alanine tails are added after the first Pro residue, yielding RAP conjugated poly-alanine tails^33^. In the wild-type strain, we observed Ala-tailed peptides at this site, whereas in the *ΔrqcH* strain, there was no detectable poly-alanine tailing despite the overexpression of YlmH; this strict requirement for RqcH for Ala-tailing is consistent with previous studies^7, 30, 32^. However, in cells lacking RqcP but overexpressing YlmH, the number of Ala-tailed peptides was similar to that observed in wild-type cells. This result directly shows that YlmH overexpression can compensate for RqcP’s role in poly-alanine tailing in RQC. Finally, when the YlmH R182A mutant was overexpressed in *ΔrqcP* cells, very few Ala-tailed peptides were detected, further supporting the functional importance of R182 of YlmH.

## Discussion

In this study, we demonstrate that YlmH can participate in the RQC pathway and is involved in poly-alanine tailing, where it can functionally substitute for its homologue RqcP. Until now, little was known about the molecular function of YlmH, with existing reports exclusively focused on the physiological effects of the *ylmH* gene loss. A large-scale *B. subtilis* gene deletion library analysis showed that, with a possible exception of growth on glycerol-ammonium minimal media, the loss of YlmH is not associated with significant fitness loss^65^. In *S. pneumoniae* disruption of the *ylmH* gene results in early cell lysis at the end of the exponential phase^66^, hinting at the factor’s role in stress tolerance. A more prominent phenotype was observed in *Synechocystis* sp. PCC 6803. In this Cyanobacterium, disruption of the *sll1252* gene encoding the YlmH orthologue leads to sensitivity to photoinhibition as well as to Ca^2+^ and Cl^-^ depletion, again pointing towards stress response defects^67^. As mentioned, YlmH is located in the *ylm* operon, located downstream of the division and cell wall (*dcw*) cluster^50^. While in some organisms, the *ylm* operon remains part of the *dcw* cluster, suggesting a potential role in cell division, it is notable that in many others, including *B. subtilis*, the *ylm* operon is separate from the *dcw* cluster^50^ (**Fig. 5b, Supplementary Table 1**).

**Fig. 5.**
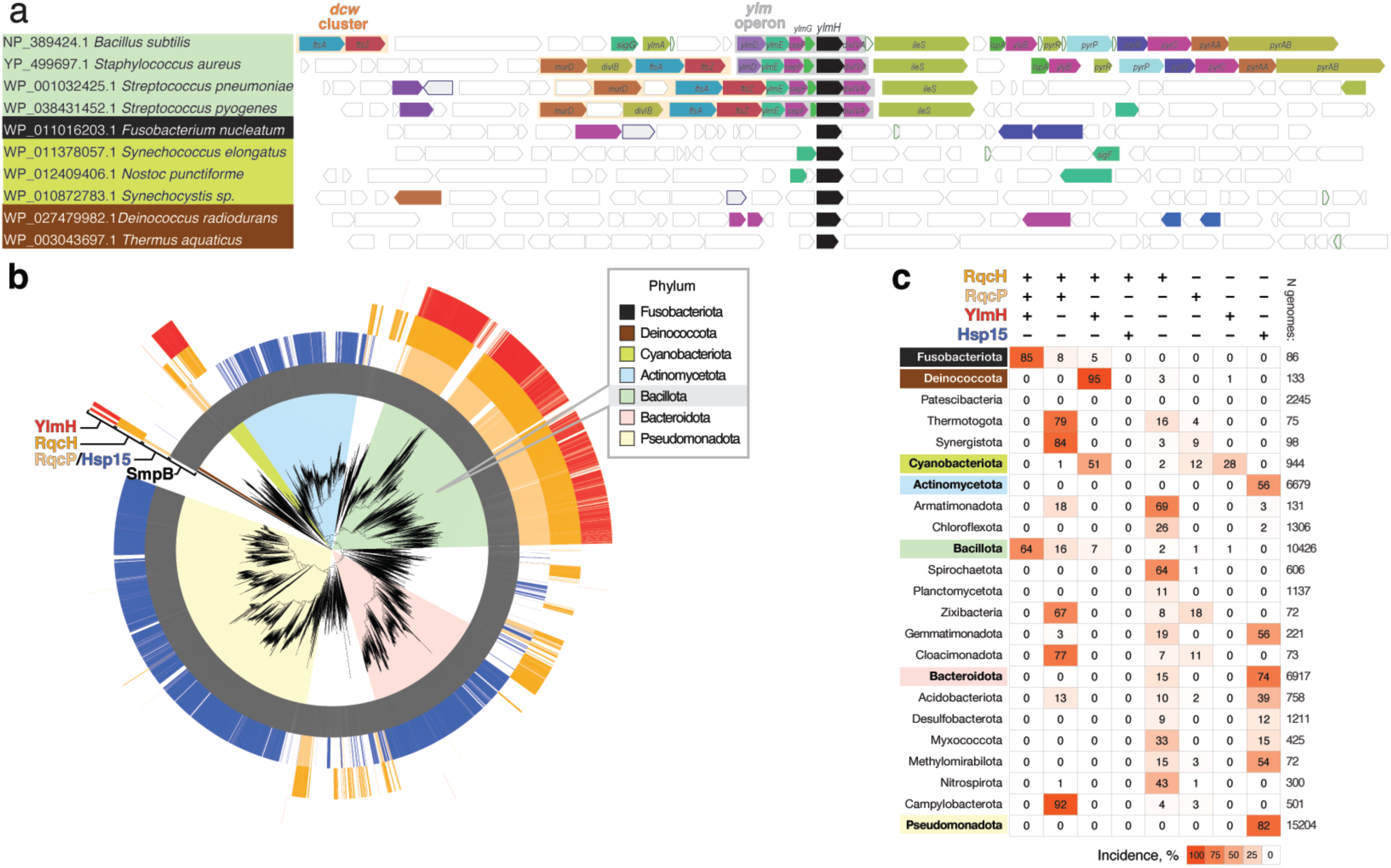
Phylogenetic distribution of the select ribosome rescue factors across bacterial taxonomy. (**a**) Genomic neighbourhood of *ylmH* gene in select species. (**b**) Genomes encoding YlmH (red), RqcH (orange), RqcP (pale orange), Hsp15 (blue) and SmpB (grey) are indicated by colours in the outer circles. The coloured segments behind branches indicate bacterial phyla. (**c**) Summary table of YlmH, RqcH, RqcP, Hsp15 and SmpB incidences in bacterial phyla.

The first indications that YlmH is involved a ribosome rescue pathway comes from our unbiased transposon mutagenesis screen identifying genes that produce a synthetic lethal phenotype when disrupted simultaneously with trans-translation (**Fig. 1a**). In addition to YlmH, we identified the alternative rescue factor BrfA, which plays a back-up function for the trans-translation system^19^, as well as RqcH and RqcP which are directly involved in the RQC pathway^28–30, 32^. Although we did not identify peptidyl-tRNA hydrolase (Pth) that was recently reported to participate in the termination phase of RQC^34^, this is not unexpected because *spoVC*, the gene encoding Pth, is essential in *B. subtilis*^68^, and, indeed, most bacteria^69, 70^. We could recapitulate the genetic interaction between *ylmH* and *ssrA* by demonstrating a slow growth phenotype of the *ΔylmH ΔssrA* double deletion strain at 37°C and genetic lethality at 49°C (**Fig. 1b**). Moreover, we could show that YlmH can bind to peptidyl-tRNA-50S subunits (**Fig. 2b**) with a binding site that is similar to that observed previously for RqcP^28–30^ (**Fig. 2d-f**). Additionally, we observed YlmH bound to peptidyl-tRNA-50S subunits containing RqcH (**Fig. 2i-j**), supporting a potential involvement of YlmH in RQC-mediated polyalanine tailing. Although deletion of YlmH did not lead to a defect in RQC-mediated polyalanine tailing, as we observed for RqcH or RqcP (**Fig. 4b**), we could show that overexpression of YlmH can rescue RQC mediated polyalanine-tailing when RqcP is absent (**Fig. 4c**). The rescue ability of YlmH does not appear to arise simply because it has an S4 domain like RqcP, but rather appears to be specific for YlmH since overexpression of other S4-containing proteins, including RspD, TyrS, TyrZ, YaaA, YhcT, YjbO, YlyB, YqxC, YtzG and RluB, does not suppress the growth defect of the *ΔrqcP ΔssrA* double deletion strain (**Supplementary Fig. 6**).

Interestingly, the growth defect of the *ΔylmH ΔssrA* strain was more dramatic than that observed for the *ΔrqcP ΔssrA and ΔrqcH ΔssrA* double-deletion strains (**Fig. 1b**), raising the question as to whether YlmH has additional key functions in *B. subtilis*. Indeed, YlmH contains additional N-terminal domains that are not present in RqcP (**Fig. 1c**), although we note that our findings indicate that the N-terminal domain of YlmH makes an important contribution to the 50S subunit binding activity (**Fig. 3b,d**). Nevertheless, strong support for a second function of YlmH comes from the cryo-EM analysis of the *ex vivo* YlmH pull-downs where we observe that the majority of the particles (66%) are bound to 50S particles lacking P-site tRNA (**Supplementary Fig. 1**). Closer examination of these particles revealed that H68 is poorly-ordered, hinting that YlmH may have an additional function in a late stage of ribosome biogenesis, however, this will require a further dedicated analysis. If YlmH does play a role in ribosome biogenesis, then this function does not appear to be essential since no growth defect was observed in *ΔylmH* strains, although such a role may account for the stronger growth defect in the *ΔylmH*-*ΔssrA* strain compared to *ΔrqcP ΔssrA and ΔrqcH ΔssrA* deletion strains (**Fig. 1b**).

Although YlmH may only play a critical role in RQC in *B. subtilis* under specific conditions, an analysis of the distribution of YlmH across more bacterial taxonomies indicates that YlmH is likely to be a dedicated component of the RQC machinery in some bacterial lineages (**Fig. 5b,c** and **Supplementary Table 1**). Indeed, a YlmH-RqcH co-distribution to the exclusion of RqcP is characteristic of Deinococcota (95% out of 133 genomes analysed), suggesting that in these bacteria the YlmH-RqcH pair alone is responsible for catalysing poly-alanine tailing. Similarly, half of the analysed cyanobacterial genomes, including the experimentally studied *Synechocystis* sp. PCC 6803, encode YlmH and RqcH, but no RqcP or Hsp15, indicative of RQC being driven by the YlmH RqcH pair in these species. By contrast, Bacillota (formerly Firmicutes) genomes, typically encode YlmH, RqcP and RqcH (64% of 10426 genomes analysed) suggesting that the functional redundancy in RQC observed in *B. subtilis* may be typical for this phylum. Similarly, 85% out of 86 Fusobacteriota genomes encode YlmH, RqcP and RqcH. The fact that both homologues have been maintained throughout evolution of different lineages of Bacillota and Fusobacteriota indicates an important partitioning of functions. S4-domain factors Hsp15 and RqcP are a largely mutually exclusive in distribution, with RqcP strongly co-distributing with RqcH^28^. Thermotogota, Synergistota, Zixibacteria, Cloacimonadota and Campylobacteria genomes predominantly encode the RqcH-RqcP pair but no YlmH (79%, 84%, 67%, 77% and 92%, respectively). While the *sll1252*-disrupted *Synechocystis* sp. PCC 6803 strain that was characterized by Inoue-Kashino and colleagues^67^ is likely to be RQC-deficient, this hypothesis awaits experimental validation. Finally, significant fractions of Armatimonadota, Spirochaetota, Myxococcota and Nitrospirota genomes (69%, 65%, 33% and 43%, respectively) encode RqcH but no RqcP or YlmH, suggesting that if these organisms do have a functional RQC pathway, some yet-to-be-identified factor performs the function of RqcP/YlmH.

## Methods

### Multiple sequence alignment

Representative sequences of RqcP and Hsp15 obtained from our previous phylogenetic study^34^ (the mmc3 dataset) were aligned with MAFFT L-INS-i version 7.453^71^ and Hidden Markov Models (HMMs) profiles were built with HMMER v3.3.2 hmmbuild^72^. The proteome from complete bacterial reference genomes (May 4, 2023) were downloaded from NCBP FTP. RqcP and Hsp15 homologues were identified using the HMMs profiles with HMMER v3.3.2 hmmscan with an E value cut-off of 1e^−10^, while YlmH and RqcH homologues were identified using HMMER v3.3.2 Jackhmmer with a maximum number of 100 iterations and an E value cut-off of 1e^−10^. YlmH sequences were aligned with MAFFT L-INS-I v7.453^71^ and visualized with Jalview^73^.

### Gene neighborhood analysis

The YlmH genomic context of selected taxa was analysed with FlaG2s^74^ (https://github.com/GCA-VH-lab/FlaGs2). YlmH accessions and their respective assemblies were used as queries in FlaGs2 with default parameters and 15 flanking genes.

### Phylogenetic distribution analysis

*Preparation of HMMs profiles*: RqcH sequences were collected by PSI-BLAST using *B. subtilis* RqcH (accession: WP_003232089.1) as the first query against refseq_select_prot database. Sequences with the Query Cover threshold exceeding 50% were collected after three iterations and then divided into each phylum. Sequence alignments were generated for each subgroup using MAFFT FFT-NS-2 v7.490^71^ and the HMMs profiles were built with HMMER v3.4 hmmbuild^72^. In a similar way, HMMs profiles for YlmH were prepared by sequences collected through a two-iteration PSI-BLAST search using *B. subtilis* YlmH (WP_003232140.1) as a query. For Hsp15 and RqcP, representative Hsp15 and RqcP homologs from previous study (see **Table S2** in Ref.^28^) were used as an input for hmmbuild. HMMs profiles for SmpB (PF01668) and other S4-related domains (PF00163, PF00472, PF00579, PF00849, PF01728, PF03462, PF13275) were downloaded from InterPro^75^.

*HMM search and curation*: homologs of YlmH, RqcH, RqcP, Hsp15 and SmpB were identified from predicted proteomes of each GTDB representative genomes, release 214.1^76^, by HMMER v3.4^72^ hmmsearch with the HMMs profiles. The GTDB representative genomes were downloaded from NCBI using NCBI dataset command-line tool v0.3. SmpB homologs were identified by selecting sequences with a bit-score for PF01668 SmpB domain exceeding the gathering cut-off value. To identify YlmH homologs, sequences with at least one of bit-scores for HMMs profiles for YlmH subgroups exceeding 100 were selected and aligned with MAFFT FFT-NS-2. Then, those having the N-terminal domains were selected as YlmH. To identify RqcH homologs, sequences with at least one of bit-scores for the HMM profiles for PqcH subgroups exceeding 80 were selected. After the MAFFT FFT-NS-2 alignment, sequences harbouring conserved Asp-Arg or similar motifs that are crucial for RqcH function^32^ were selected and categorized as RqcH. RqcP and Hsp15 homologs were identified according the following procedure. Sequences with a bit-score for HMMs profiles for RqcP or Hsp15 exceeding 40 were selected. Subsequently, those with a bit-score for HMMs profiles for the following domains equal to or greater than the gathering cuttoff value were excluded: the N-terminal Ribosomal_S4 (PF00163) domain, which is found within uS4 proteins, the N-terminal tRNA-synt_1b (PF00579) domain, which is found in TyrS and TyrZ, the C-terminal PseudoU_synth_2 (PF00849) domain, which is found in RluB, YhcT, YlyB and YtzG, and the C-terminal FtsJ (PF01728) domain, which is found in YqxC. In addition, those with a bit-score for the S4_2 (PF13275) ≥45, and those classified as YlmH homologs were excluded. Sequences harbouring highly conserved Arg2 and Asp4 (*B. subtilis* numbering) were selected and among them, those possessing the K_14_-R_15_-R_16_ motif and having a distance from Arg2 (or the corresponding Arg) to the C-terminus ≤110 residues were classified as RqcP; the remaining were classified as Hsp15 (see **Fig. 3D** in Ref.^28^).

*Phylogenetic tree*: Phylogenetic tree downloaded from GTDB^76^ were pruned to retain only the genomes used in the analysis by APE^77^ and visualized and annotated by iTol v. 6^78^.

### Tn-seq sampling, library preparation and analysis

*Preparation of transposon-inserted cell library*: *in vitro* transposition experiment was performed as described previously^79^. Briefly, the *himar1* transposon harbouring a kanamycin resistant marker gene was transposed from the plasmid pKIG645 by MarC9 transposase^79^ into purified chromosomal DNA derived from either wild-type 168 strain or its *ssrA* deletion mutant derivative (VHB257). The transposon-inserted DNA, in which gaps were filled, was used for the transformation of both the 168 strain and the Δ*ssrA* mutant. Transformants were selected on LB agar plate supplemented with 3 µg/mL kanamycin following an overnight incubation at 37°C. 2x10^5^ and 1.5x10^5^ cells were separately pooled for wild-type and Δ*ssrA* strain, respectively, and stored as transposon-inserted cell libraries with 25% glycerol at – 80°C.

*DNA library preparation and deep DNA sequencing*: DNA libraries were prepared as per previously established protocol with minor modifications^80^. Briefly, cell libraries (3x10^7^ – 2x10^8^ cells) were grown in 50 mL of LB medium supplemented with 3 µg/mL kanamycin at 37°C until OD_600_ of 0.5. The cells were lysed with Nucleobond buffer set iii (MACHEREY-NAGEL) and the chromosomal DNA was purified with Sera-Mag magnetic beads (Cytiva). The resulting DNA sample was then digested by MmeI (NEB) and re-purified on magnetic beads. DNA adapters were ligated to MmeI-digested DNA fragments by T4 DNA ligase (Takara), and the ligation products were purified on magnetic beads. Subsequently, PCR amplification was performed using primers ADPT-Tnseq-PCRPrimer (5′- AATGATACGGCGACCACCGAGATCTACACTCTTTCCCTACACGACGCTCTTCCGATCT -3′) and P1-M6-GAT-MmeI (5′- CAAGCAGAAGACGGCATACGAGATAGACCGGGGACTTATCATCCAACCTGT-3′), and the resulting products were resolved by polyacrylamide gel electrophoresis. DNA fragments with length of ≈148 bp were eluted from the crushed gels and purified by Nucleospin Gel and PCR Cleanup column. The DNA libraries were sequenced on the NextSeq 500 sequencing system (Illumina).

*Tn-seq data analysis:* reads were processed as per the MAGenTA protocol^81^ using an in-house script developed with SeqKit^82^, and processed reads were aligned to the reference *B. subtilis* 168 genomic sequence (Assembly: ASM904v1) using Bowtie version 1.3.1^83^. Fitness calculations were performed in accordance with the MAGenTA protocol using custom scripts implemented in R^84^. The data are provide as **Supplementary Table 2** and the code is provided as **SupplementaryCode.zip**.

### Bacterial strains and plasmids

Strains (and information regarding their sensitivity to lincomycin), plasmids, oligonucleotides and synthetic DNA sequences used in this study are provided in **Supplementary Table S3**.

### Growth assays

*B. subtilis* 168 wild-type and deletion strains were pre-grown on LB plates overnight at 30°C. Fresh individual colonies were used to inoculate liquid LB medium cultures to starting OD_600_ of 0.01) at 37°C. When the cultures reached mid-log phage (OD_600_ of ≈0.4), they were diluted to OD_600_ of 0.1. The resultant cultures were used to prepare 10- to 10^5^-fold serial dilutions which were then spotted onto LB agar plates with or without 1 mM IPTG. The plates were scored after 18 hours incubation at either 37°C or 49°C.

### Sucrose gradient fractionation

The experiments were performed as described previously^85^. Briefly, *B. subtilis* strains were pre-grown on Luria Broth (LB) plates overnight at 30°C. Fresh individual colonies were used to inoculate 200 mL LB cultures. The cultures were grown until OD_600_ of 0.8 at 37°C, the cells were collected by centrifugation and dissolved in 0.5 mL of HEPES:Polymix buffer [5 mM Mg(OAc)_2_]^85^. Cells were lysed using FastPrep homogenizer (MP Biomedicals) and the resultant lysates were clarified by centrifugation. 10 A_260_ units of each extract were loaded onto 10%-35% (w/v) sucrose density gradients prepared in HEPES:Polymix buffer [5 mM Mg(OAc)_2_]. Gradients were resolved at 36,000 rpm for 3 hours at 4°C. Both preparation and fractionation of gradients was done using a Biocomp Gradient Station (BioComp Instruments), and A_280_ was used as a readout during the fractionation.

For immunoblotting 0.5 mL fractions were precipitated with ethanol, pelleted by centrifugation, the supernatants were discarded and the samples were dried. The pellets were resuspended in 2x SDS loading buffer (100 mM Tris-HCl pH 6.8, 4% SDS (w/v) 0.02% Bromophenol blue, 20% glycerol (w/v) 4% β-mercaptoethanol), resolved on the 11% SDS-PAGE and transferred to nitrocellulose membrane. FLAG-tagged proteins were detected using anti-FLAG M2 primary (Sigma-Aldrich, F1804) antibodies combined with anti-mouse-HRP secondary (Rockland; 610-103-040; 1:10,000 dilution) antibodies. Enhanced chemiluminescence (ECL) detection was performed using WesternBright^TM^ Quantum (K-12042-D10, Advansta) Western blotting substrate using ImageQuant LAS 4000 (GE Healthcare) imaging system.

### ApdA reporter detection through Western blotting

Fresh colony of strains expressing the ApdA reporter was inoculated into 1 ml LB medium dispensed into plastic 96 deep well plate (Treff Lab) and grown at 30°C for 18 h with shaking at 1200 rev per min using DWMax MBR-034P (Taitec) shaking incubator. Each 20 µL of overnight culture was transferred to 1 mL fresh LB medium in the presence and absence of 20 µM Bortezomib dispensed into plastic 96 deep-well plate and grown at 37°C with shaking until OD_600_ of 0.15 as monitored by Multiskan GO (Thermo Scientific). A 0.75 mL of the culture was collected, mixed with 83 µL of 50% TCA and kept on ice for 5 min. After a 2-minute centrifugation at 13,500 rpm per min (4°C), the cell pellet was resuspended in 500 µL of 0.1 M Tris-HCl (pH 6.8). After an additional 2-minute centrifugation at 13,500 rpm per min (4°C) the cell pellet was resuspended in lysis buffer (0.5 M sucrose, 20 mM MgCl_2_, 1 mg/mL lysozyme, 20 mM HEPES:NaOH, pH7.5) and incubated at 37°C for 10 min. Next, 2× SDS sample buffer (4% SDS, 30% glycerol, 250 mM Tris pH 6.8, 1 mM DTT, saturated bromophenol blue) was added and the lysate was denatured at 85 °C for 5 min. Proteins were resolved on 11% SDS-PAGE and transferred to PVDF membrane. NanoLuc-tagged proteins were detected using anti-NanoLuc, (Promega, N7000, 1:5,000 dilution) combined with anti-mouse-HRP secondary (Bio-Rad, #1706516, 1:5,000 dilution) antibodies. FLAG-tagged proteins were detected using anti-DYKDDDDK tag primary antibodies (Wako, 1E6, 1:5,000 dilution) combined with anti-mouse-HRP secondary antibodies (Bio-Rad, #1706516, 1:5,000 dilution). SigA proteins were detected using anti-SigA^85^ (1:10,000 dilution) combined with anti-rabbit-HRP secondary (Bio-Rad, #1706515, 1:5,000 dilution) antibodies. Images were obtained using Amersham Imager 600 (GE Healthcare) luminoimager and analyzed using ImageJ^86^.

### MS analysis of tagging sites on the reporter protein

The experiments were performed as described previously^33^. Strains expressing the ApdA reporter were grown in 100 mL of LB with 20 µM of bortezomib until OD_600_ of 0.5. Cells were harvested by centrifugation and frozen at –80°C. Cells were then thawed in 2X Celln Lytic B cell lysis reagent (Sigma) and 0.2 mg/mL lysozyme for 10 min gently rocking. The lysate was clarified by centrifugation at 20,000 x g for 30 min at 4°C. 50 uL of Strep-tactin Sepharose beads (IBA) were added to the supernatant, and then the samples were incubated for 1 h at 4°C. The beads were washed four times 5 min each at 4°C with IP wash buffer (20 mM Tris pH 8, 100 mM NH_4_Cl, 0.4% Triton X-100, 0.1% NP-40). Protein was eluted with elution buffer (20 mM Tris pH 8, 100 mM NH_4_Cl, 5 mM Desthiobiotin) by shaking at 4°C for 1 h. Then, 36 uL of elution was reduced with 100 mM DTT in 100 mM triethylammonium bicarbonate (TEAB) buffer at 58°C for 55 min and then the pH was adjusted to 8. The samples were alkylated with 200 mM iodoacetamide in 100 mM TEAB buffer in the dark at room temperature for 15 min. Proteins were pelleted and resuspended in 100 mM TEAB and proteolyzed with 13.8 ng/µL of LysC (Wyco) at 37°C overnight. Peptides were desalted on Oasis u-HLB plates (Waters), eluted with 60% acetonitrile (ACN)/0.1% trifluoracetic acid (TFA), dried and reconstituted with 2% ACN/0.1% formic acid.

Desalted peptides cleaved by LysC were analyzed by LC-MS/MS. Then peptides were separated by reverse-phase chromatography (2-90% ACN/0.1% formic acid gradient over 85 min at a rate of 300 nL/min) on a 75 um x 150 mm ReproSIL-Pur-120-c18-AQ column (Dr. Albin Maisch, Germany) using the nano-EasyLC 1200 system (Thermo). Eluting peptides were sprayed into an Orbitrap-Lumos_ETD mass spectrometer through a 1-um emitter tip (New Objective) at 2.4 kV. Scans were acquired within 350-1800 Da m/z targeting the truncated reporter. Precursor ions were individually isolated and were fragmented (MS/MS) using an HCD activation collision energy of 30. Precursor (fragment) ions were analyzed with a 200 automatic gain control (AGC), 118 ms maximum injection time (IT) at 60,000 resolution with 6 cycles. The MS/MS spectra were processed with Proteome Discover v2.5 (Thermo) and were analyzed with Mascot v2.8.2 (Matrix Science) using RefSeq2021_204_Bacillus.S and a database with peptides from the NanoLuc-AdpA-BleR reporter protein. Peptide identifications from Mascot searches were processed within the Proteome Discoverer-Percolator to identify peptides with a confidence threshold of a 1% false discovery rate, as determined by an auto-concatenated decoy database search. Proteomics results are provided as **Supplementary Table 4**.

### Cryo-EM sample preparation

Cryo-EM sample preparation was performed as established in previous work^87^ Briefly, *B. subtilis* 168 cells VHB715 (Δ*ylmH trpC2*; Δ*ylmH*::*spcR*) expressing YlmH-FLAG from pHT01 p43 wRBS ylmH-GS5-FLAG wt tGyrA were grown at 37°C in Lysogeny-Broth (LB) medium (Roth) supplemented with 5 μg/mL chloramphenicol and shaking at 145 rpm until OD_600_ of 1.5. Cells were collected in 25 mM HEPES-KOH pH 7.5, 100 mM potassium acetate, 15 mM magnesium acetate, 0.1% (v/v) NP-40 and 0.5 mM Tris carboxy ethyl phosphene (TCEP) buffer supplemented with protease inhibitor cocktail (Roche), flash-frozen in liquid nitrogen and lysed under cryogenic conditions using a Retsch MM400 (Retsch). The lysate was cleared at 16,000 rpm in a JA-25.50 rotor (Beckman Coulter) for 15 min and incubated with anti-Flag M2 affinity beads (Merck) for 1.5 hours at 4°C on a turning wheel. After in-batch wash with 20 ml lysis buffer without protease inhibitors, the beads were transferred to a Mobicol column and washed with 4 ml of 25 mM HEPES-KOH, pH 7.5, 100 mM potassium acetate, 15 mM magnesium acetate, 0.01% DDM (dodecylmaltoside) and 0.5 mM TCEP buffer, after which the ylmH complexes were eluted using 0.2 mg ml^−1^ 3× Flag peptide (Sigma) in wash buffer. The complexes were then applied to grids for electron microscopy analysis or resolved on 4–12% NuPAGE SDS–PAGE gels (Invitrogen) by staining with Instant Blue (Expedeon). Sample volumes of 3.5 µl (8 OD_260_ per ml) were applied to grids (Quantifoil, Cu, 300 mesh, R3/3 with 3 nm carbon) which had been freshly glow-discharged using a GloQube (Quorum Technologies) in negative charge mode at 25 mA for 90 s. Sample vitrification was performed using an ethane-propane mixture (37:63) in a Vitrobot Mark IV (Thermo Fisher Scientific), the chamber was set to 4°C and 100% relative humidity and blotting was done for 3 s with no drain or wait time.

### Cryo-EM data collection

Data were collected in an automated manner using EPU v.2.6 on a cold-FEG fringe-free Titan Krios G4 (Thermo Fisher Scientific) transmission electron microscope operating at 300 kV equipped with a Falcon 4 direct electron detector (Thermo Fisher Scientific). The camera was operated in electron counting mode and data were collected at a magnification of 96,000× with the nominal pixel size of 0.83 Å and a nominal defocus range of −0.4 to −0.9 μm. A total of 25,683 micrographs in EER format were collected with 5.14 e/px/s and 5.31 s of exposure (corresponding to a total dose of 40 e per Å^2^ on the specimen). No statistical methods were used to predetermine the sample size. The sample size was selected on the basis of a three-day data collection, which was chosen to obtain a sufficient number of particles for data processing.

### Cryo-EM data processing

Processing was performed using RELION 4.0.1^88, 89^. The pixel size for processing was adjusted to 0.8 Å from the nominal 0.83 Å during data collection owing to best correlation with published ribosome models at this pixel size. Movie frames were aligned with MotionCor2^90^ using 5×5 patches followed by CTF estimation of the resulting micrographs using CTFFIND4^91^ using power spectra from the MotionCor run. The CTF fits were used to remove outlier micrographs with estimated resolutions greater than 15 Å, which retained 25,457 micrographs. crYOLO 1.8.3^92, 93^ with its general model (gmodel_phosnet_202005_N63_c17.h5) was used for particle picking, which resulted in 2,550,984 particles. These were extracted in a box size of 64 px at a pixel size of 4.8 Å and subjected to 2D classification.

After 2D classification, 2,391,525 particles resembling 50S subunits were selected and used for a first 3D auto-refinement to centre all particles for further refinement steps. An empty mature 50S subunit was used as reference, with the initial volume being generated from PDB ID 6HA8, Ref.^94^. Afterwards, particles were extracted with re-centring from the previous Refine3D-job at a box size of 128 px and a pixel size of 2.4 Å. The particles were aligned into a 3D volume using the output of the initial Refine3D-job as a reference (re-scaled to the new box and pixel sizes). From these aligned particles, 3D classification was performed without further angular sampling. Particle sorting was performed according to **Supplementary Fig. 1**. Particles for final classes of the main YlmH-state and the YlmH-RqcH-state were re-extracted at a box size of 384 px with a pixel size of 0.8 Å and subjected to 3D auto-refinement. Particles for the main YlmH-state were further CTF-refined to correct for anisotropic magnification, trefoil and higher-order aberrations, defocus and astigmatism. Furthermore, particles for the main YlmH-state were subsequently subjected to Bayesian polishing followed by another round of CTF refinements. After these procedures, the final volumes were generated by 3D auto-refinement and postprocessing in RELION.

### Molecular model building

The initial model for the 50S subunit of the main YlmH-bound state was generated based on a published *B. subtilis* 70S structure (PDB ID: 6HA8; Ref.^94^). For YlmH, an AlphaFold^60, 95^ model was used and rigid-body fitted into the density using ChimeraX^96, 97^. Afterwards, the model was manually adjusted in Coot^98, 99^. Model refinement was performed using REFMAC5 as implemented in Servalcat^100^ using protein restraints generated by ProSmart from AlphaFold models for YlmH, uL5, uL6 and bL31^60, 95, 101, 102^. Subsequently, the model for the YlmH-RqcH-state was derived by iterative adjustment from the main model, to this end an AlphaFold model of RqcH was added and settled into density. For refinement of the RqcH-state, ProSmart restraints for YlmH, uL5, uL6, uL13, uL18, bL21, uL23, bL31, as well as LibG^103^ restraints for RNA were applied. Cryo-EM data collection, refinement and validation statistics for all models are listed in **Table 1**.

### Figure preparation

UCSF ChimeraX 1.6.1^104^ was used to isolate density and visualize density images and structural superpositions. Figures were assembled with Adobe Illustrator (latest development release, regularly updated) and Inkscape v1.3.

### Data availability

Cryo-EM maps have been deposited in the Electron Microscopy Data Bank (EMDB) with accession codes EMD-19638 [https://www.ebi.ac.uk/pdbe/entry/emdb/EMD-19638] (*YlmH- 50S-P-tRNA complex*), EMD-19641 [https://www.ebi.ac.uk/pdbe/entry/emdb/EMD-19641] (*YlmH-50S-RqcH-A-tRNA and P-tRNA complex*). Molecular models have been deposited in the Protein Data Bank with accession code 8S1P [https://doi.org/10.2210/pdb8S1P/pdb] (*YlmH-50S-P-tRNA complex*) and 8S1U [https://doi.org/10.2210/pdb8S1U/pdb] (*YlmH-50S- RqcH-A-tRNA and P-tRNA complex*). Sequencing data generated in this study is available at the DDBJ Sequence Read Archive (BioProject accession number PRJDB17234). Tn-Seq fitness data used across the manuscript can be found in **Supplementary Table 2** and the code is provided as **SupplementaryCode.zip**. Results of the MS analysis of tagging sites on the reporter protein are provided as a **Supplementary Table 4**. MS proteomics data for the YlmH pulldown samples analysis have been deposited to the ProteomeXchange Consortium via the PRIDE^105^ partner repository (project accession: PXD049370, project DOI: 10.6019/PXD049370).

## Acknowledgments

This work was supported by the Deutsche Forschungsgemeinschaft (DFG) (grant WI3285/11-1 to D.N.W), the Estonian Research Council (PRG335 to T.T.), the Knut and Alice Wallenberg Foundation (project grant 2020-0037 to G.C.A. and V.H.), the Swedish Research Council (Vetenskapsrådet) grants (2019-01085 and 2022-01603 to G.C.A., 2021-01146 to V.H.), Crafoord foundation (project grant Nr 20220562 to V.H.), the Estonian Research Council (PRG335 to V.H.), Cancerfonden (20 0872 Pj to V.H.), NIH grant GM136960 (to A.R.B.), postdoctoral grant from the Umeå Centre for Microbial Research, UCMR (to H.T.), JST, ACT X, Japan (JP1159335 to H.T.), MEXT, JSPS Grant-in-Aid for Scientific Research (grants 20H05926 and 21K06053 to S.C., 23K05017 to H.T., 21K15020 to K.F.), Institute for Fermentation, Osaka (grant G-2021-2-063 to S.C.). Part of this work was performed at the Multi-User CryoEM Facility at the Centre for Structural Systems Biology, Hamburg, supported by the Universität Hamburg and DFG grant numbers (INST 152/772-1|152/774-1|152/775-1|152/776-1|152/777-1 FUGG).

## Author contributions

K.F. performed Tn-Seq and analyzed the data. H.T. performed genetics and molecular biology experiments. L.D.P. generated the native YlmH-50S samples. H.P. prepared and screened the cryo-EM grids. B.B. collected high resolution cryo-EM data. H.P. processed the cryo-EM data, as well as generated and refined the molecular models. G.C.A, K.F. and J.A.N. performed bioinformatic analyses. A.R.B., S.C., T.T. provided materials and supervised the project. M.S. performed mass spectrometry on the native YlmH:50S samples and E.N.P. performed mass spectrometry on the ApdA reporter samples. V.H., D.N.W., H.P. and H.T. wrote the manuscript with input from all authors. V.H., D.N.W., H.P. and H.T. conceived and supervised the project.

## Additional information

**Supplementary information** The online version contains supplementary material available at https://…

**Correspondence** and requests for materials should be addressed to H.T. and H.P.

---

**Supplementary Fig. 1.**
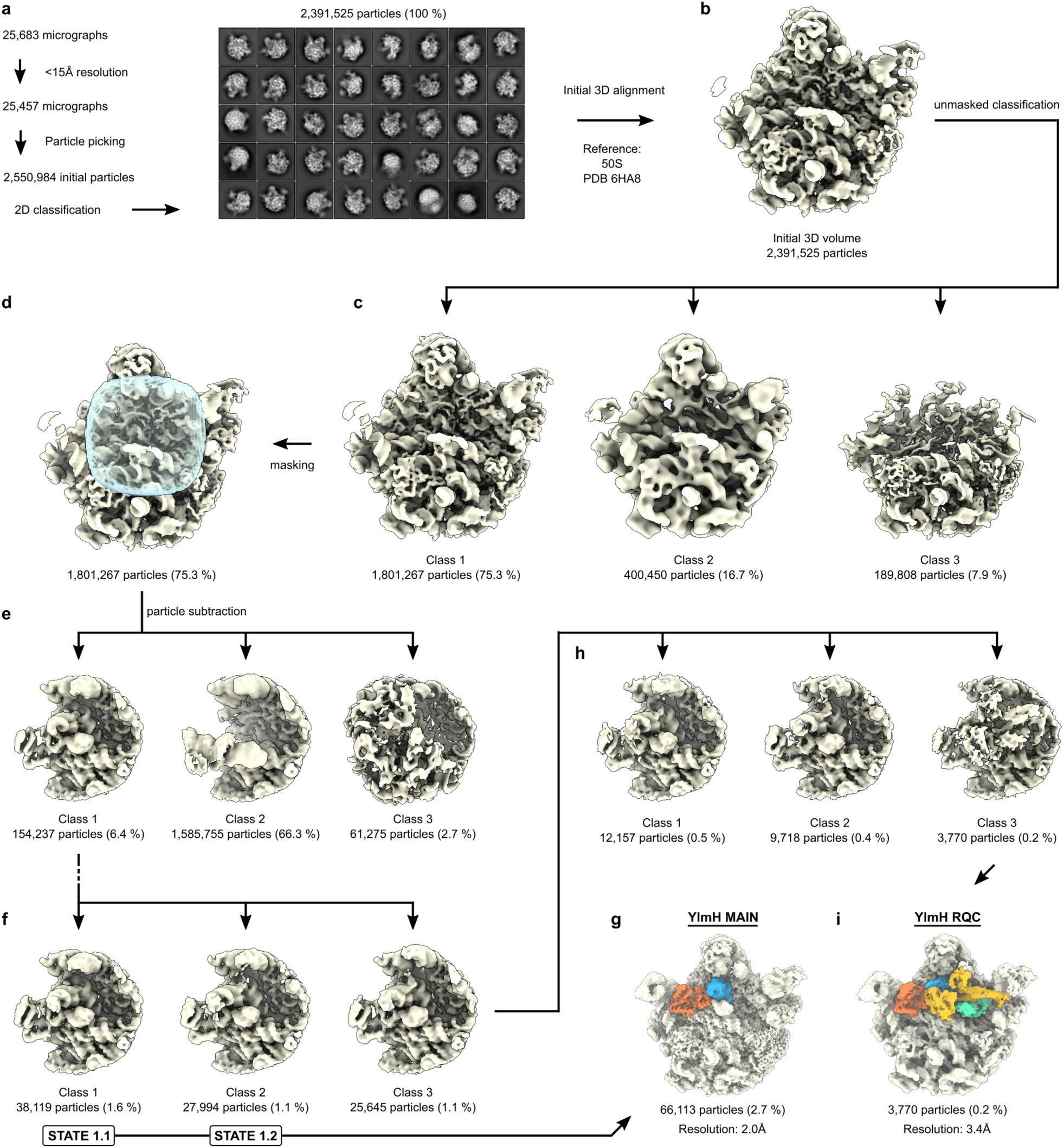
Data processing of the YlmH-50S complexes. (**a**) 26,683 micrographs were collected and 25,457 remained after applying a resolution cut-off. 2,550,984 particles were selected from the micrographs and subjected to 2D classification. Of these 2,391,525 (termed 100%) were (**b**) 3D aligned and subjected to unmasked 3D classification, resulting in (**c**) three classes containing 50S or 50S-like particles. The major class 1 was (**d**) masked and subjected to 3D classification with particle subtraction, resulting in (**e**) three classes, only one of which (class 1) contained density for YlmH and P-tRNA. (**f**) Class 1 was further subjected to further 3D classification resulting in two well-defined states 1.1 and 1.2 (class 1 and 2) that were combined (66,113 particles, 2.7%), yielding (**g**) a final reconstruction of YlmH-P-tRNA-50S complex at 2.0 Å, which was termed State 1. (**h**) Class 3 was further subsorted into three classes, one of which (class 3 containing 3,770 particles, 0.2%) contained additional density that was further refined to yield (**i**) a cryo-EM structure of the YlmH-RqcH-50S complex with A- and P-tRNAs at 3.4 Å, which was referred to as State 2.

**Supplementary Fig. 2.**
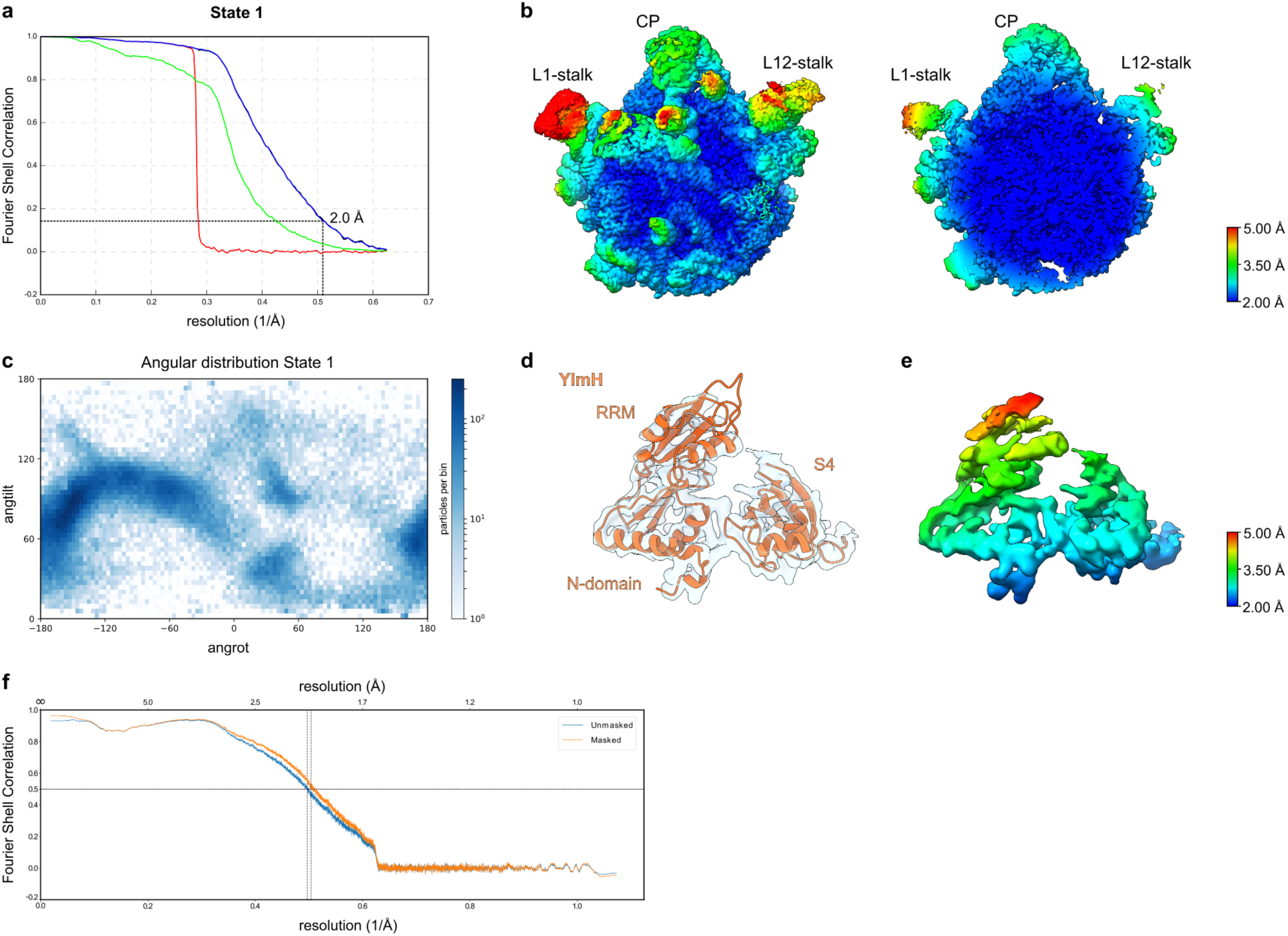
Cryo-EM structure of the YlmH-50S complex (State 1). (**a**) Fourier Shell Correlation (FSC) curves for State 1 with the dashed line at 0.143 indicating an average resolution of 2.0 Å. The different curves include the masked map (green), unmasked map (blue), the phase-randomized masked map (red). (**b**) Cryo-EM map of State 1 colored according to local resolution with (left panel) interface overview and (right panel) transverse section revealing core of the 50S subunit. Landmarks for L1-stalk, L12-stalk and central protuberance (CP) are indicated. (**c**) Angular distribution of the particles that comprise State 1. (**d**) Isolated cryo-EM density (grey transparency) and molecular model (orange) for YlmH. (**e**) Isolated cryo-EM density as in (**d**) but coloured according to local resolution. (**f**) FSC map versus model with curves shown for unmasked (blue) and masked (orange) volumes.

**Supplementary Fig. 3.**
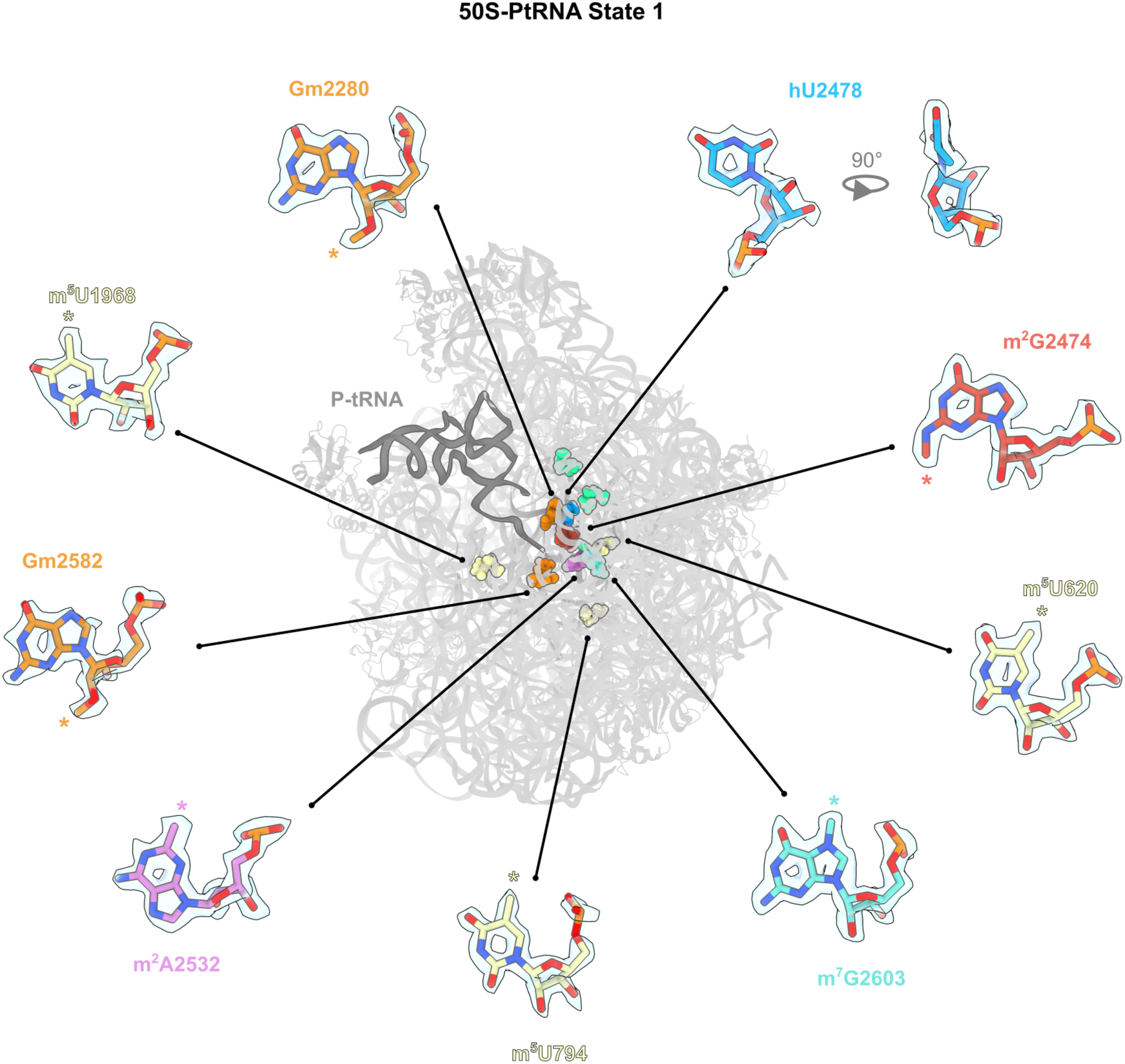
Identification of rRNA modifications in the YlmH-50S complex (State 1). Sideview (from L1 side) of the 50S subunit (grey) with P-tRNA (dark grey) for reference, showing the position of 23S rRNA modifications identified in the cryo-EM of State 1. Enlargements show isolated cryo-EM density (grey transparency) with molecular models for each modification. Asterisks indicate the position of the modification.

**Supplementary Fig. 4.**
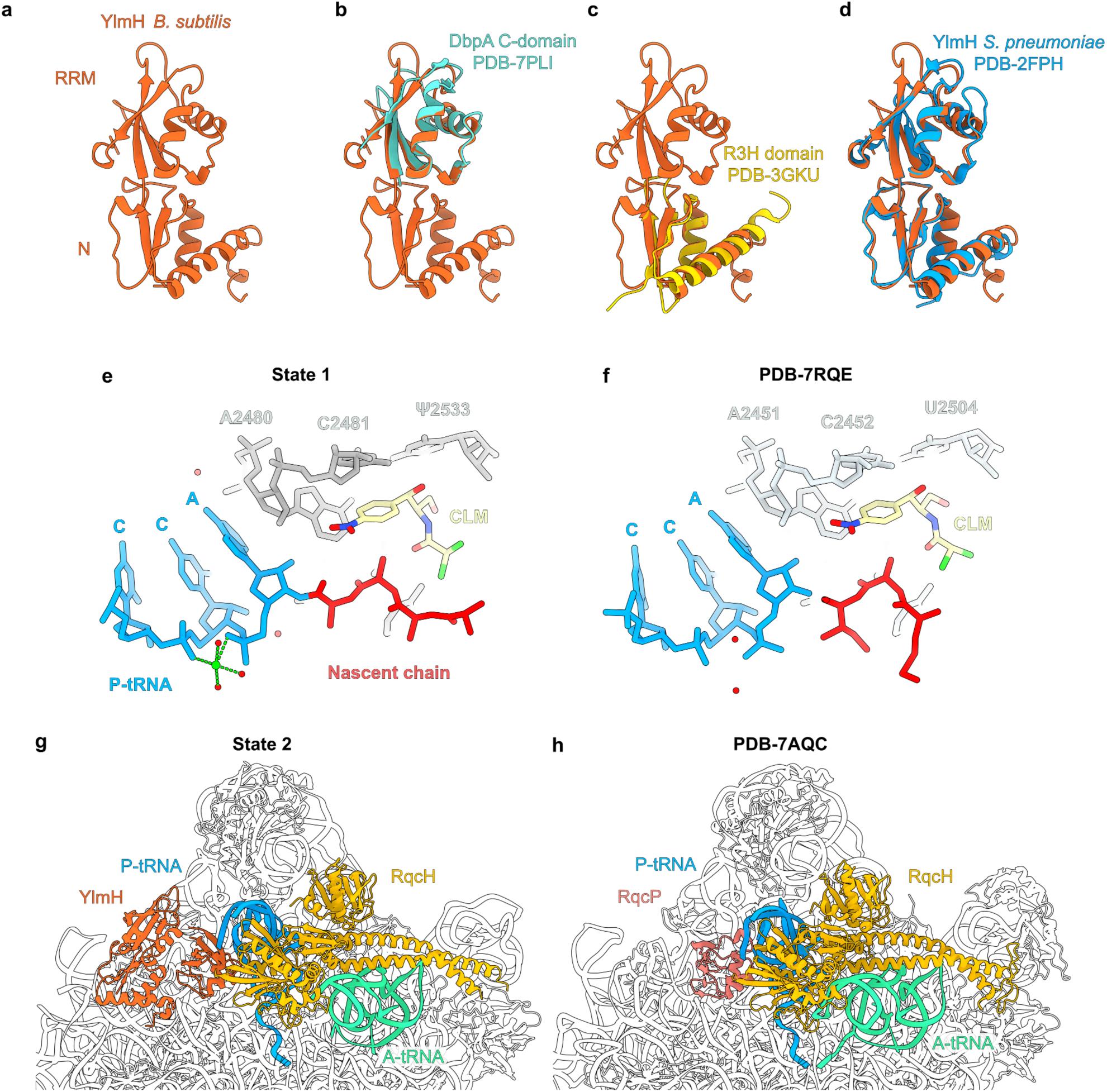
Comparison of YlmH State 1 and 2 with previous structures. (**a-d**) Comparison of the RRM and N-domain of YlmH (orange) from State 1 with (**b**) the C-domain of DbpA (cyan, PDB ID 7PLI)^59^, (**c**) the R3H domain of a probable RNA-binding protein from Clostridium symbiosum ATCC 14940 (PDB ID 3GKU), and (**d**) the crystal structure of the RRM and N-domain of YlmH from *Streptococcus pneumoniae* (PDB ID 2FPH). (**e-f**) Comparison of the CCA-end of the P-tRNA (blue), nascent chain (red), chloramphenicol (yellow) and selected 23S rRNA nucleotides (grey) from (**e**) YlmH-50S complex State 1 and (**f**) the crystal structure of the *Thermus thermophilus* 70S ribosome in complex with protein Y, A-site deacylated tRNA analog CACCA, P-site MAI-tripeptidyl-tRNA analog ACCA-IAM, and chloramphenicol (PDB ID 7RQE)^52^. (**g-h**) Comparison of 50S subunit (white ribbons) in complex with YlmH (orange), RqcH (mustard), P-tRNA (blue) and A-tRNA (green) from (**g**) the YlmH-RqcH-50S complex with A- and P-tRNA (State 2), with (**h**) the structure of the bacterial RQC complex containing RqcP (orange), RqcH (mustard), P-tRNA (blue) and A-tRNA (green) (PDB ID 7AQC)^29^.

**Supplementary Fig. 5.**
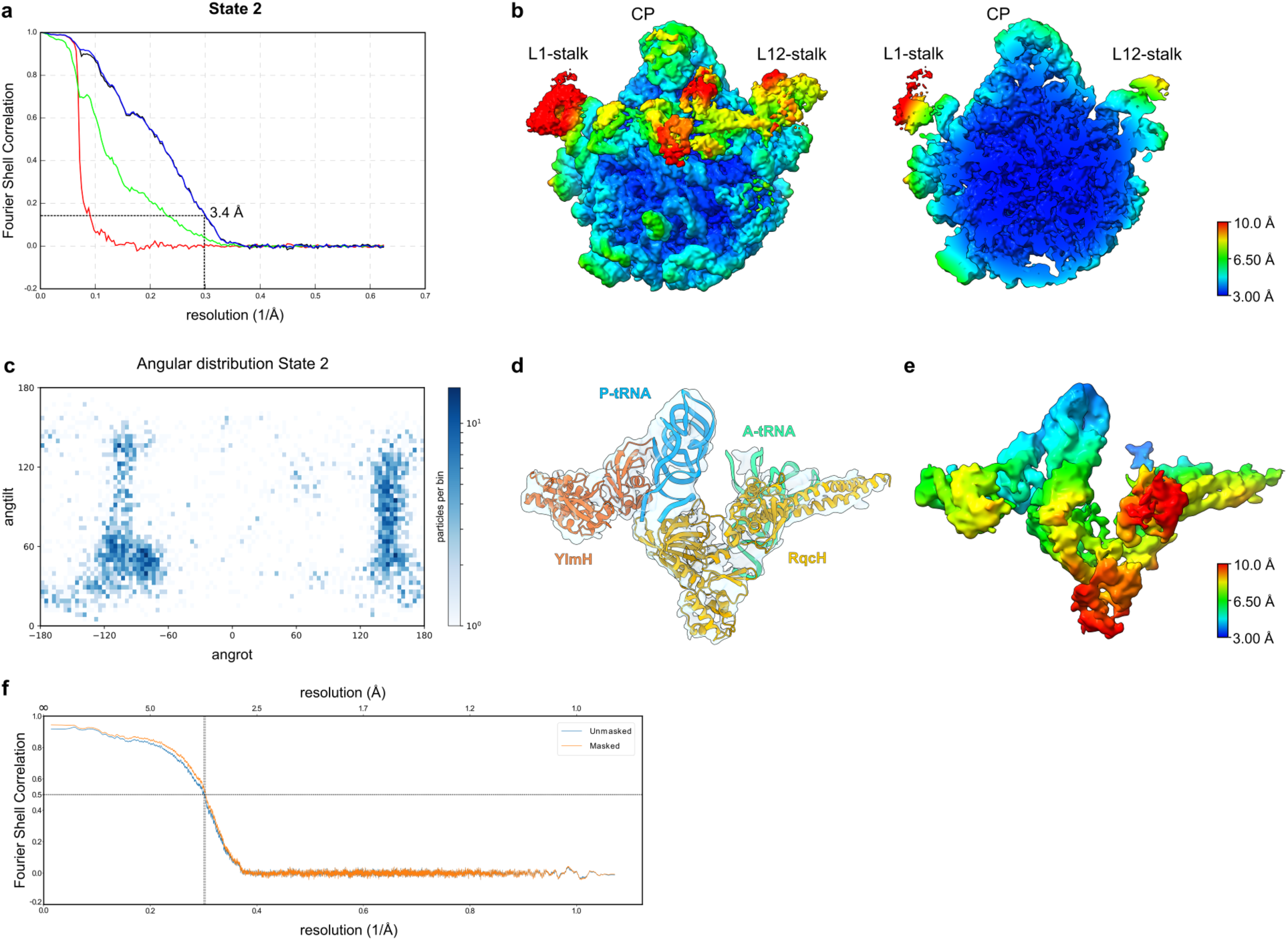
Cryo-EM structure of the YlmH-RqH-50S complex (State 2). (**a**) Fourier Shell Correlation (FSC) curves for State 2 with the dashed line at 0.143 indicating an average resolution of 3.4 Å. The different curves include the masked map (green), unmasked map (blue), the phase-randomized masked map (red). (**b**) Cryo-EM map of State 2 colored according to local resolution with (left panel) interface overview and (right panel) transverse section revealing core of the 50S subunit. Landmarks for L1-stalk, L12-stalk and central protuberance (CP) are indicated. (**c**) Angular distribution of the particles that comprise State 2. (**d**) Isolated cryo-EM density (grey transparency) and molecular model for YlmH (orange), RqcH (mustard), P-tRNA (blue) and A-tRNA (green). (**e**) Isolated cryo-EM density as in (**d**) but coloured according to local resolution. (**f**) FSC map versus model with curves shown for unmasked (blue) and masked (orange) volumes.

**Supplementary Fig. 6.**
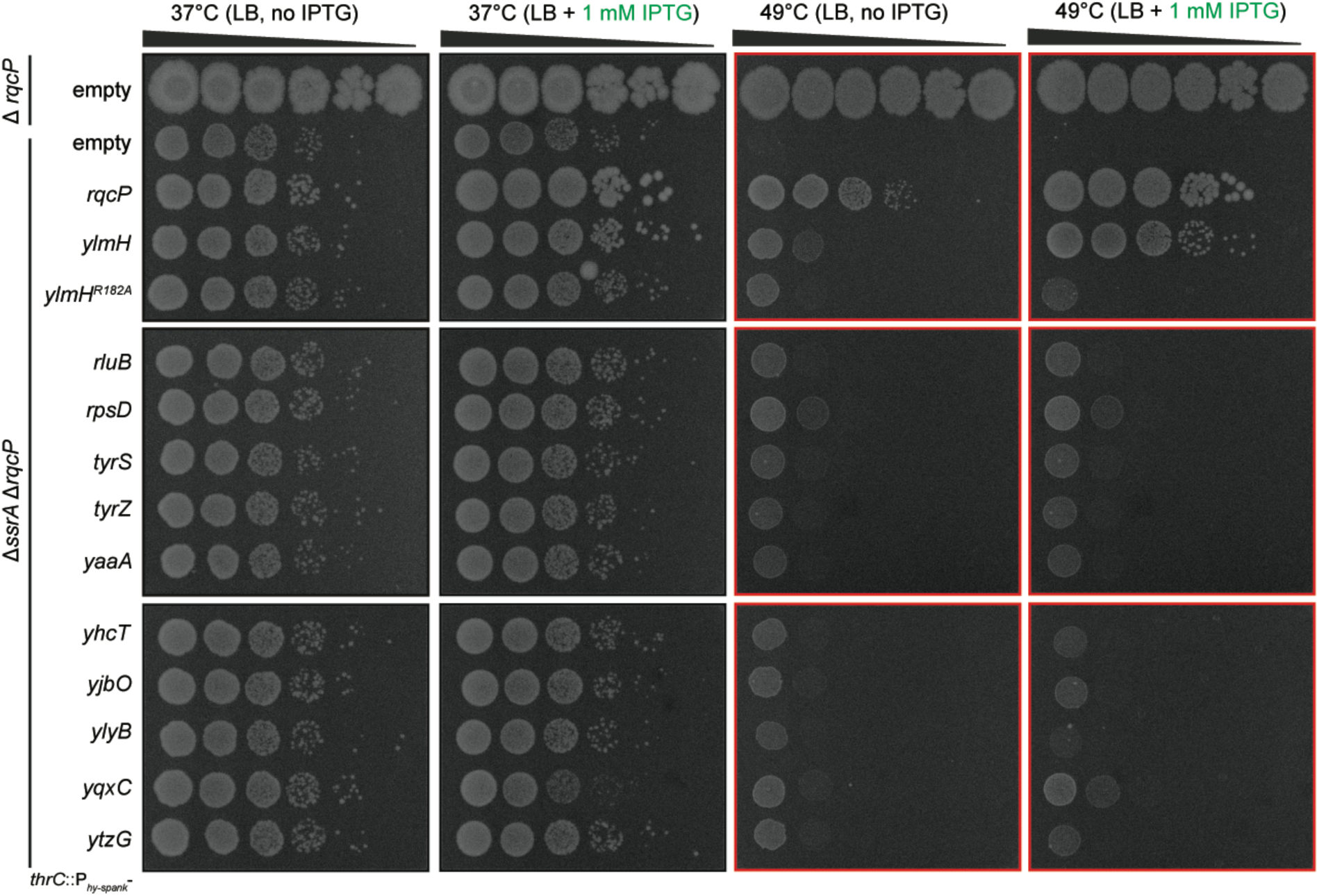
Ectopic expression of YlmH but of not other tested *B. subtilis* S4 domain-encoding proteins suppresses the temperature sensitivity of the Δ*ssrA* Δ*rqcP* strain. Expression of S4-domain-encoding *B. subtilis* proteins [negative control strain (BCHT1328), *rqcP* (BCHT1338), *ylmH* (BCHT1329), *ylmH^R^*^182^*^A^* (BCHT1330), *rluB* (BCHT1332), *rpsD* (BCHT1339), *tyrS* (BCHT1340), *tyrZ* (BCHT1341), *yaaA* (BCHT1331), *yhcT* (BCHT1333), *yjbO* (BCHT1334), *ylyB* (BCHT1335), *yqxC* (BCHT1336) or *ytzG* (BCHT1337)] cloned under the control of P*_hy-spank_* promotor the in Δ*ssrA* Δ*rqcP* deletion background was induced by 1 mM IPTG. 10-fold serial dilutions were spotted onto LB agar plates with or without 1 mM IPTG and plates were scored after 18-hours incubation at either 37°C or 49°C.

