## Supplementary Code for "A role for the S4-domain containing protein YlmH in ribosome-associated quality control in *Bacillus subtilis*": curation_RqcH.R

```
# RqcH analysis

# input -------------------------------------------------------------------

input_d1 <- 'hmm_search/output/PF05833/PF05833.csv'
input_d2 <- 'hmm_search/output/PF05670/PF05670.csv'
input_d3 <- 'hmm_search/output/PF18297/PF18297.csv'

# path to a list of genome id
input_genome_list <- 'genome_list/genome_id_for_analysis.csv'
input_path2genome_files='hmm_search/target/hmm_target_list.txt'

output <- 'output/RqcH.csv'

# set iTol dataset values
dataset_label = 'RqcH'
output_tree_dataset= 'tree/4_dataset_protein/iTol_RqcH.txt'


# package -----------------------------------------------------------------

library(tidyverse)
library(Biostrings)
library(tidysq)
library(ips)
library(patchwork)

# function ----------------------------------------------------------------


source('fn_SelectiveReadAAStringSet.R')
source('fn_write_Fasta.R')


# load --------------------------------------------------------------------

# RqcH domains
# PF05833
# PF05670
# PF18297

df_dom <- 
  tibble(
    domain_name = c('NFACT_N', 'NFACT-R_1', 'NFACT-R_2'),
    Pfam = c('PF05833', 'PF05670', 'PF18297'),
    dom_order = c(1:3),
    GA = c(31, 29.8, 26.9)
  ) %>% print()


# load genome id list
df_genome_list <- read_csv(input_genome_list) %>% print()


# wrangling ---------------------------------------------------------------

input_RqcH <- fs::dir_ls('hmm_search/output/') %>% str_subset('RqcH') %>% purrr::map(fs::dir_ls, type = 'file') %>% 
  purrr::map(str_subset, 'csv$') %>% 
  purrr::list_simplify()
df_results <- c(input_d1, input_d2, input_d3, input_RqcH) %>% 
  purrr::map(readr::read_csv) %>% print()


df_dom_all <- df_results %>%
  purrr::map(dplyr::select, protein_id = target_name, genome_id = genome, query_name, score_fullseq) %>% 
  purrr::map2(str_c('d', 1:26), ~ dplyr::mutate(.x, 'dom' = .y)) %>% 
  purrr::list_rbind() %>% 
  nest(data=c(query_name, score_fullseq)) %>% 
  tidyr::pivot_wider(id_cols = c(protein_id, genome_id), names_from = dom, values_from = data) %>% 
  unnest(col = c(d1:d26), names_sep = '.')


v_name_score_fullseq <- str_c('d', 4:26, '.score_fullseq')

df_RqcH <- df_dom_all %>% 
  filter(
    if_any(
      .cols = all_of(v_name_score_fullseq),
      .fns = \(x) x >=80)
  ) %>%
  left_join(df_genome_list %>% select(genome_id, genome_database, gtdb_taxonomy, phylum, class), by = 'genome_id') %>% 
  print()

  
# get protein sequence

# get path to protein.faa
path_faa <- read_csv(input_path2genome_files, col_names = c('path')) %>% 
  mutate(genome_id = str_extract(path, 'GC[AF]_\\d+.\\d+')) %>% 
  filter(genome_id %in% df_RqcH$genome_id) %>% 
  distinct() %>% 
  pull(path)


TargetSeqSet <- path_faa %>% 
  purrr::map(fn_SelectiveReadAAStringSet, df_RqcH$protein_id) %>% 
  purrr::list_rbind() %>% 
  distinct() %>% 
  print()

df_RqcH_s <- df_RqcH %>% left_join(TargetSeqSet) %>% print()

# Analyse seq -------------------------------------------------------------

df_RqcH_s %>% #count(phylum, sort = T)
  add_count(phylum) %>% 
  mutate(
    phylum_c = case_when(
      phylum %in% c('p__Bacillota', 'p__Bacillota_A') ~ str_remove(phylum, 'p__'),
      str_detect(phylum, 'Bacillota') ~ 'Bacillota_Other',
      n >= 10 ~ str_remove(phylum, 'p__'),
      n < 10 ~ 'others' 
    ),
    phylum_c = if_else(is.na(phylum_c), 'Others', phylum_c)
  ) %>% 
  select(phylum_c, protein_id, seq) %>% 
  distinct() %>% 
  group_by(phylum_c) %>% 
  group_walk(
    ~write_fasta(
      x = as.sq(.x$seq),
      name = paste0(.x$protein_id),
      file = file.path(
        str_c(
          'output/fasta_raw/RqcH_hmm_',
          .y$phylum_c,
          ".fasta"))
    )
  )

# prep msa by mafft (FFTNS)
# script: mafft_fftns_RqcH.sh

list_files_mafft <- fs::dir_ls('output/mafft_raw/fftns/', type = 'file') %>% str_subset('fftns/RqcH')


mafft_res <- tibble(
  files = list_files_mafft,
  X1 = list_files_mafft %>% 
    purrr::map(Biostrings::readAAStringSet)
) %>% 
  transmute(
    group = str_remove_all(files, paste0(c('mafft/fftns/', '.txt'), collapse = '|')),
    seq = map(X1, ~as.character(.))
    ) %>% 
  unnest_longer(col = seq) %>% 
  print()


list_DR_fftns <- read_csv('input/tmp_RqcH_residues.csv')

mafft_res %>% left_join(list_DR_fftns) %>% 
  mutate(DR = str_sub(seq, Dx, Rx)) %>% 
  count(group, DR, sort = T) %>% 
  arrange(group, desc(n)) %>% 
  write_csv('c.csv')
  
v_RqcH <- mafft_res %>% left_join(list_DR_fftns) %>% 
  mutate(DR = str_sub(seq, Dx, Rx)) %>% 
  filter(DR %in% c('DR','ER','DK','EK','NR','NK','QR')) %>% 
  select(seq_id) %>% distinct() %>% pull()

df_RqcH_s_sel <- df_RqcH_s %>% 
  filter(protein_id %in% v_RqcH) %>% 
  mutate(protein = 'RqcH')

# output ------------------------------------------------------------------


df_RqcH_s_sel %>% 
  write_csv(output)


# iTol --------------------------------------------------------------------

# prep iTol dataset keywords
df_dataset_keywords <- tibble(
  lines = c(
    'DATASET_COLORSTRIP',
    'SEPARATOR',
    'DATASET_LABEL',
    'COLOR,#005AFF',
    'STRIP_WIDTH',
    'DATA'
  )) %>% 
  mutate(
    lines = case_when(
      lines == 'SEPARATOR' ~ str_c(lines, ' ', 'COMMA'),
      lines == 'DATASET_LABEL' ~ str_c(lines, ',', dataset_label),
      lines == 'STRIP_WIDTH' ~ str_c(lines, ',', 200),
      !is.na(lines) ~ lines,
    )
  )

# prep iTol dataset
df_dataset_data <- df_RqcH_s_sel %>%
  select(genome_id) %>% 
  distinct() %>% 
  mutate(
    genome_id_num = str_extract(genome_id, '\\d+'),
    color = '#FF4B00'
  ) %>% 
  transmute(
    dataset_row = str_c(
      'CONTAINS==',
      genome_id_num,
      ',',
      color
    )
  )

write_lines(
  df_dataset_keywords$lines,
  output_tree_dataset
)

df_dataset_data %>% 
  write_delim(output_tree_dataset, append = TRUE, col_names = FALSE)
```
