## Supplementary Code for "A role for the S4-domain containing protein YlmH in ribosome-associated quality control in *Bacillus subtilis*": curation_RqcP_HslR.R

```
# RqcP/Hsp15 analysis

# input -------------------------------------------------------------------

input_d1 <- 'hmm_search/output/rqcP_Hsp15/rqcP.csv' # result of RqcP
input_d2 <- 'hmm_search/output/rqcP_Hsp15/Hsp15.csv' # result of RqcP
input_d3 <- 'hmm_search/output/PF01479/PF01479.csv' # S4 (RqcP/Hsp15)
input_d4 <- 'hmm_search/output/PF13275/PF13275.csv' # S4_2 (YaaA)
input_d5 <- 'hmm_search/output/PF00163/PF00163.csv' # Ribosomal_S4 (RpsD)
input_d6 <- 'hmm_search/output/PF00849/PF00849.csv' # PseudoU_synth_2 (RluA)
input_d7 <- 'hmm_search/output/PF00579/PF00579.csv' # tRNA-synt_1b (TyrS/TyrZ)
input_d8 <- 'hmm_search/output/PF01728/PF01728.csv' # FtsJ (YqxC)
input_ylmH <- 'output/YlmH.csv'

# path to a list of genome id
input_genome_list <- 'genome_list/genome_id_for_analysis.csv'
input_path2genome_files='hmm_search/target/hmm_target_list.txt'


output <- 'output/RqcP_Hsp15_4216.csv'

# set iTol dataset values
dataset_label = 'RqcP_Hsp15'
output_tree_dataset= 'tree/4_dataset_protein//iTol_RqcP_Hsp15_4216.txt'

# package -----------------------------------------------------------------


library(tidyverse)
library(Biostrings)
library(tidysq)


# function ----------------------------------------------------------------


source('fn_SelectiveReadAAStringSet.R')


# load --------------------------------------------------------------------

# S4 related domains
df_dom <- 
  tibble(
    domain_name = c('RqcP', 'Hsp15', 'S4', 'S4_2', 'Ribosomal_S4', 'PseudoU_synth_2', 'tRNA-synt_1b', 'FtsJ'),
    Pfam = c('RqcP','Hsp15','PF01479','PF13275','PF00163','PF00849','PF00579','PF01728'),
    dom_order = c(1:8),
    GA = c(NA, NA, 22.8, 24.6, 23, 26.8, 21.1, 22.2)
  ) %>% print()

df_YlmH <- read_csv(input_ylmH)

# load genome id list
df_genome_list <- read_csv(input_genome_list) %>% print()


# wrangling ---------------------------------------------------------------

df_results <- c(
  input_d1, input_d2, input_d3,
  input_d4, input_d5, input_d6, 
  input_d7, input_d8 ) %>% 
  purrr::map(readr::read_csv)

# get all accession number of RqcP/Hsp15-like proteins (d1~d3)
v_id_RqcP_Hsp15 <- df_results[1:2] %>% 
  purrr::map(select, protein_id = target_name) %>% 
  purrr::list_rbind() %>% distinct() %>% pull()

df_dom_all <- df_results %>%
  purrr::map(dplyr::select, protein_id = target_name, genome_id = genome, query_name, score_fullseq, description_of_target) %>% 
  purrr::map2(str_c('d', 1:8), ~ dplyr::mutate(.x, 'dom' = .y)) %>% 
  purrr::list_rbind() %>% 
  filter(protein_id %in% v_id_RqcP_Hsp15) %>% 
  nest(data=c(query_name, score_fullseq, description_of_target)) %>% 
  tidyr::pivot_wider(id_cols = c(protein_id, genome_id), names_from = dom, values_from = data) %>% 
  unnest(col = c(d1:d8), names_sep = '.') %>% 
  select(-ends_with('description_of_target'), d1.description_of_target, d2.description_of_target)


df_RqcP_Hsp15_raw <- df_dom_all %>% 
  # remove YlmH
  left_join(df_YlmH %>% select(protein_id, protein) %>% distinct(), by = 'protein_id') %>%
  filter(is.na(protein)) %>% 
  
  filter(
    # remove proteins which has Ribosomal_S4 domain
    # some RqcP/Hsp15-like proteins (whose dom1.score_fullseq or dom2.score_fullseq is relatively very high) shows also over the GA value of dom4.score_fullseq, thus we set a cut off value of 45 that is not strict
    d4.score_fullseq < 45 | is.na(d4.score_fullseq)
  ) %>%
  filter(
    # remove proteins which has Ribosomal_S4 domain
    d5.score_fullseq < filter(df_dom, dom_order == 5) %>% pull(GA) | is.na(d5.score_fullseq)
  ) %>%
  filter(
    # remove proteins which has PseudoU_synth_2 domain
    d6.score_fullseq < filter(df_dom, dom_order == 6) %>% pull(GA) | is.na(d6.score_fullseq)
  ) %>%
  filter(
    # remove proteins which has tRNA-synt_1b domain
    d7.score_fullseq < filter(df_dom, dom_order == 7) %>% pull(GA) | is.na(d7.score_fullseq)
  ) %>%
  filter(
    # remove proteins which has FtsJ domain
    d8.score_fullseq < filter(df_dom, dom_order == 8) %>% pull(GA) | is.na(d8.score_fullseq)
  ) %>% 
  filter(
    # select score S4 > GA
    d3.score_fullseq >= filter(df_dom, dom_order == 3) %>% pull(GA)
  ) %>% 
  filter(d1.score_fullseq > 40 | d2.score_fullseq > 40) %>% 
  mutate(
    d1.score_fullseq = if_else(is.na(d1.score_fullseq), 0, d1.score_fullseq),
    d2.score_fullseq = if_else(is.na(d2.score_fullseq), 0, d2.score_fullseq),
  ) %>% 
  left_join(df_genome_list %>% select(genome_id, genome_database, gtdb_taxonomy, phylum, class), by = 'genome_id') %>% 
  filter(is.na(protein)) %>% 
  print()

# get protein sequence

# get path to protein.faa
path_faa <- read_csv(input_path2genome_files, col_names = c('path')) %>% 
  mutate(genome_id = str_extract(path, 'GC[AF]_\\d+.\\d+')) %>% 
  distinct() %>% 
  pull(path)


TargetSeqSet <- path_faa %>% 
  purrr::map(fn_SelectiveReadAAStringSet, df_RqcP_Hsp15_raw$protein_id) %>% 
  purrr::list_rbind() %>% 
  distinct() %>% 
  print()

# TargetSeqSet %>%
#   write_csv('output/tmp_RqcP_seq_32119.csv')
# TargetSeqSet <- read_csv('output/tmp_RqcP_seq_32119.csv')

df_RqcP_Hsp15_raw_s <- df_RqcP_Hsp15_raw %>% 
  left_join(TargetSeqSet) %>%
  mutate(
    seq_len = str_length(seq),
    IsPartial = str_detect(d1.description_of_target, '[Pp]artial')
  ) %>% 
  filter(IsPartial == FALSE) %>%
  filter(seq_len < 310) %>% # remove sequences that is too long
  print()


df_RqcP_Hsp15_sel <- df_RqcP_Hsp15_raw_s %>% 
  mutate(
    protein = case_when(
      d1.score_fullseq > d2.score_fullseq ~ 'RqcP',
      d1.score_fullseq <= d2.score_fullseq ~ 'Hsp15'
    )
  )

df_RqcP_Hsp15_sel %>% glimpse()


# output ------------------------------------------------------------------


df_RqcP_Hsp15_sel %>% 
  write_csv(output)


# iTol --------------------------------------------------------------------

# prep iTol dataset keywords
df_dataset_keywords <- tibble(
  lines = c(
    'DATASET_COLORSTRIP',
    'SEPARATOR',
    'DATASET_LABEL',
    'COLOR,#005AFF',
    'STRIP_WIDTH',
    'DATA'
  )) %>% 
  mutate(
    lines = case_when(
      lines == 'SEPARATOR' ~ str_c(lines, ' ', 'COMMA'),
      lines == 'DATASET_LABEL' ~ str_c(lines, ',', dataset_label),
      lines == 'STRIP_WIDTH' ~ str_c(lines, ',', 200),
      !is.na(lines) ~ lines,
    )
  )

# prep iTol dataset
df_dataset_data <- df_RqcP_Hsp15_sel %>% 
  select(genome_id, protein) %>% 
  distinct() %>% 
  mutate(
    genome_id_num = str_extract(genome_id, '\\d+'),
    color = case_when(
      protein == 'RqcP' ~ '#ff4b00',
      protein == 'Hsp15' ~ '#005AFF'
    )
  ) %>% 
  transmute(
    dataset_row = str_c(
      'CONTAINS==',
      genome_id_num,
      ',',
      color
    )
  )

write_lines(
  df_dataset_keywords$lines,
  output_tree_dataset
)

df_dataset_data %>% 
  write_delim(output_tree_dataset, append = TRUE, col_names = FALSE)
```
