## Supplementary Code for "A role for the S4-domain containing protein YlmH in ribosome-associated quality control in *Bacillus subtilis*": curation_SmpB.R

```
# SmpB analysis

# input -------------------------------------------------------------------

input_d1 <- 'hmm_search/output/PF01668/PF01668.csv'

# path to a list of genome id
input_genome_list <- 'genome_list/genome_id_for_analysis.csv'

output <- 'output/SmpB.csv'

# set iTol dataset values
dataset_label = 'SmpB'
output_tree_dataset = 'tree/4_dataset_protein/iTol_SmpB.txt'

# package -----------------------------------------------------------------

library(tidyverse)

# load --------------------------------------------------------------------

# SmpB domains

df_dom <- tibble(
  domain_name = c('SmpB'),
  Pfam = c('PF01668'),
  dom_order = c(1),
  GA = c(27)
) %>% print()


# load hmmsearch result table
df_hmmsearch_d1 <- read_csv(input_d1) %>% dplyr::rename(genome_id = genome) %>% distinct() %>% print()


# load genome id list
df_genome_list <- read_csv(input_genome_list) %>% print()

# wrangling ---------------------------------------------------------------


# get all accession number
df_prot_all <- df_hmmsearch_d1 %>% 
  dplyr::select(
    protein_id = target_name, 
    genome_id
  ) %>% 
  distinct() %>% arrange(genome_id) %>% print()

df_dom_all <- df_prot_all %>% 
  dplyr::left_join(df_hmmsearch_d1, by = c('protein_id' = 'target_name', 'genome_id')) %>%
  dplyr::rename(
    d1.name = query_name,
    d1.accession = query_accession,
    d1.E_fullseq = E_fullseq,
    d1.score_fullseq = score_fullseq,
    d1.description_of_target = description_of_target
  ) %>% 
  distinct() %>% 
  print()

df_SmpB <- df_dom_all %>% 
  filter(
    # select only proteins whose score_fullseq > gathering cutoff
    d1.score_fullseq >= filter(df_dom, dom_order == 1) %>% pull(GA)
  ) %>% 
  mutate(
    protein = 'SmpB'
  ) %>%
  left_join(df_genome_list %>% select(genome_id, genome_database, gtdb_taxonomy, phylum), by = 'genome_id') %>% 
  print()

# output ------------------------------------------------------------------


df_SmpB %>% 
  filter(protein == 'SmpB') %>% 
  write_csv(output)


# iTol --------------------------------------------------------------------

# prep iTol dataset keywords
df_dataset_keywords <- tibble(
  lines = c(
    'DATASET_COLORSTRIP',
    'SEPARATOR',
    'DATASET_LABEL',
    'COLOR,#005AFF',
    'STRIP_WIDTH',
    'DATA'
  )) %>% 
  mutate(
    lines = case_when(
      lines == 'SEPARATOR' ~ str_c(lines, ' ', 'COMMA'),
      lines == 'DATASET_LABEL' ~ str_c(lines, ',', dataset_label),
      lines == 'STRIP_WIDTH' ~ str_c(lines, ',', 200),
      !is.na(lines) ~ lines,
    )
  )

# prep iTol dataset
df_dataset_data <- df_SmpB %>% 
  select(genome_id) %>% 
  distinct() %>% 
  mutate(
    genome_id_num = str_extract(genome_id, '\\d+'),
    color = '#666666'
  ) %>% 
  transmute(
    dataset_row = str_c(
      'CONTAINS==',
      genome_id_num,
      ',',
      color
    )
  ) %>% print()

write_lines(
  df_dataset_keywords$lines,
  output_tree_dataset
)

df_dataset_data %>% 
  write_delim(output_tree_dataset, append = TRUE, col_names = FALSE)
```
