## Supplementary Code for "A role for the S4-domain containing protein YlmH in ribosome-associated quality control in *Bacillus subtilis*": curation_YlmH.R

```
# YlmH analysis

# input -------------------------------------------------------------------

input_d1 <- 'hmm_search/output/PF21278/PF21278.csv'
input_d2 <- 'hmm_search/output/PF17774/PF17774.csv'
input_d3 <- 'hmm_search/output/PF01479/PF01479.csv'

# path to a list of genome id
input_genome_list <- 'genome_list/genome_id_for_analysis.csv'
input_path2genome_files='hmm_search/target/hmm_target_list.txt'

output <- 'output/YlmH.csv'

# set iTol dataset values
dataset_label = 'YlmH'
output_tree_dataset = 'tree/4_dataset_protein/iTol_YlmH.txt'


# package -----------------------------------------------------------------

library(tidyverse)
library(Biostrings)
library(tidysq)
library(ips)
library(patchwork)

# function ----------------------------------------------------------------

source('fn_SelectiveReadAAStringSet.R')
source('fn_write_Fasta.R')

# load --------------------------------------------------------------------

# YlmH domains
# PF21278
# PF17774
# PF01479

df_dom <- 
tibble(
  domain_name = c('YlmH_1st', 'YlmH_RBD', 'S4'),
  Pfam = c('PF21278', 'PF17774', 'PF01479'),
  dom_order = c(1:3),
  GA = c(27,23,22.8)
) %>% print()


# load genome id list
df_genome_list <- read_csv(input_genome_list) %>% print()

# wrangling ---------------------------------------------------------------

input_YlmH <- fs::dir_ls('hmm_search/output/') %>% str_subset('YlmH') %>% purrr::map(fs::dir_ls, type = 'file') %>% 
  purrr::map(str_subset, 'csv$') %>% 
  purrr::list_simplify()
df_results <- c(input_d1, input_d2, input_d3, input_YlmH) %>% 
  purrr::map(readr::read_csv) %>% print()

df_dom_all <- df_results %>%
  purrr::map(dplyr::select, protein_id = target_name, genome_id = genome, query_name, score_fullseq) %>% 
  purrr::map2(str_c('d', 1:12), ~ dplyr::mutate(.x, 'dom' = .y)) %>% 
  purrr::list_rbind() %>% 
  nest(data=c(query_name, score_fullseq)) %>% 
  tidyr::pivot_wider(id_cols = c(protein_id, genome_id), names_from = dom, values_from = data) %>% 
  unnest(col = c(d1:d12), names_sep = '.')


v_name_score_fullseq <- str_c('d', 4:12, '.score_fullseq')

df_YlmH <- df_dom_all %>% 
  filter(
    if_any(
      .cols = all_of(v_name_score_fullseq),
      .fns = \(x) x >=100)
  ) %>% 
  left_join(df_genome_list %>% select(genome_id, genome_database, gtdb_taxonomy, phylum, class), by = 'genome_id') %>% 
  print()


# get protein sequence
# get path to protein.faa
path_faa <- read_csv(input_path2genome_files, col_names = c('path')) %>% 
  mutate(genome_id = str_extract(path, 'GC[AF]_\\d+.\\d+')) %>% 
  filter(genome_id %in% df_YlmH$genome_id) %>% 
  distinct() %>% 
  pull(path)


TargetSeqSet <- path_faa %>% 
  purrr::map(fn_SelectiveReadAAStringSet, df_YlmH$protein_id) %>% 
  purrr::list_rbind() %>% 
  distinct() %>% 
  print()

df_YlmH_s <- df_YlmH %>% left_join(TargetSeqSet) %>% print()


# Analyse seq -------------------------------------------------------------

df_YlmH_s %>%
  mutate(
    phylum_c = case_when(
      phylum == 'p__Bacillota' ~ str_remove(phylum, 'p__'),
      phylum == 'p__Bacillota_A' ~ str_remove(phylum, 'p__'),
      str_detect(phylum, 'Bacillota') ~ 'Bacillota_Other',
      phylum == 'p__Cyanobacteriota' ~ str_remove(phylum, 'p__'),
      phylum == 'p__Deinococcota' ~ str_remove(phylum, 'p__'),
      phylum == 'p__Fusobacteriota' ~ str_remove(phylum, 'p__'),
    ),
    phylum_c = if_else(is.na(phylum_c), 'Others', phylum_c)
  ) %>% #count(phylum, phylum_c, sort = T) %>%
  select(phylum_c, protein_id, seq) %>% 
  distinct() %>% 
  group_by(phylum_c) %>% 
  group_walk(
    ~write_fasta(
      x = as.sq(.x$seq),
      name = paste0(.x$protein_id),
      file = file.path(
        str_c(
          'output/fasta_raw/YlmH_hmm_',
          .y$phylum_c,
          ".fasta"))
    )
  )


# prep msa by mafft
# script: 5.1_mafft_fftns_YlmH.sh


# import mafft result (fasta file)
fn_read_mafft = function(x, Ns, Ne, Rs, Re, Ss, Se){
  tmp_mafft <- readAAStringSet(x)
  output <- tibble(
    protein_id = names(tmp_mafft), 
    alignment = as.character(tmp_mafft),
    subseq_Nt = str_sub(alignment, start = Ns, end = Ne), # Extract YlmH-characteristic N-terminal sequence
    subseq_RBD= str_sub(alignment, start = Rs, end = Re), 
    subseq_S4 = str_sub(alignment, start = Ss, end = Se) # Extract R181-R197 region
    )
}


mafft_y_B <- fn_read_mafft('output/mafft_raw/fftns/YlmH_hmm_Bacillota_fftns.txt', 110-109, 183, 183+1, 382-1, 382, -1)
mafft_y_BA <- fn_read_mafft('output/mafft_raw/fftns/YlmH_hmm_Bacillota_A_fftns.txt', 160-159, 740, 740+1, 1231-1, 1231, -1)
mafft_y_BX <- fn_read_mafft('output/mafft_raw/fftns/YlmH_hmm_Bacillota_Other_fftns.txt', 90-89, 170, 170+1, 312-1, 312, -1)
mafft_y_CYA <- fn_read_mafft('output/mafft_raw/fftns/YlmH_hmm_Cyanobacteriota_fftns.txt', 90-89, 170, 170+1, 280-1, 280, -1)
mafft_y_DIN <- fn_read_mafft('output/mafft_raw/fftns/YlmH_hmm_Deinococcota_fftns.txt', 36-35, 77, 77+1, 185-1, 185, -1)
mafft_y_FSO <- fn_read_mafft('output/mafft_raw/fftns/YlmH_hmm_Fusobacteriota_fftns.txt', 44-43, 87, 87+1, 246-1, 246, -1)
mafft_y_X <- fn_read_mafft('output/mafft_raw/fftns/YlmH_hmm_Others_fftns.txt', 41-40, 120, 120+1, 220-1, 220, -1)

# merge tibble
mafft_y_merge <- mafft_y_B %>% 
  bind_rows(mafft_y_BA) %>% 
  bind_rows(mafft_y_BX) %>% 
  bind_rows(mafft_y_CYA) %>% 
  bind_rows(mafft_y_DIN) %>% 
  bind_rows(mafft_y_FSO) %>% 
  bind_rows(mafft_y_X) %>% 
  distinct() %>% 
  print()
  
df_YlmH_s2 <- df_YlmH_s %>% 
  left_join(mafft_y_merge) %>% 
  filter(str_detect(subseq_Nt, '\\w')) %>% # remove proteins which don't have the N-terminal region
  filter(str_detect(subseq_S4, '\\w')) %>% # remove proteins which don't have the N-terminal region
  mutate(protein = 'YlmH') %>% 
  print()


# output ------------------------------------------------------------------


df_YlmH_s2 %>% 
  write_csv(output)


# iTol --------------------------------------------------------------------

# prep iTol dataset keywords
df_dataset_keywords <- tibble(
  lines = c(
    'DATASET_COLORSTRIP',
    'SEPARATOR',
    'DATASET_LABEL',
    'COLOR,#005AFF',
    'STRIP_WIDTH',
    'DATA'
  )) %>% 
  mutate(
    lines = case_when(
      lines == 'SEPARATOR' ~ str_c(lines, ' ', 'COMMA'),
      lines == 'DATASET_LABEL' ~ str_c(lines, ',', dataset_label),
      lines == 'STRIP_WIDTH' ~ str_c(lines, ',', 200),
      !is.na(lines) ~ lines,
    )
  )

# prep iTol dataset
df_dataset_data <- df_YlmH_s2 %>% 
  distinct() %>% 
  mutate(
    genome_id_num = str_extract(genome_id, '\\d+'),
    color = '#ff4b00'
  ) %>% 
  transmute(
    dataset_row = str_c(
      'CONTAINS==',
      genome_id_num,
      ',',
      color
    )
  )

write_lines(
  df_dataset_keywords$lines,
  output_tree_dataset
)

df_dataset_data %>% 
  write_delim(output_tree_dataset, append = TRUE, col_names = FALSE)
```
