## Supplementary Code for "A role for the S4-domain containing protein YlmH in ribosome-associated quality control in *Bacillus subtilis*": fn_SelectiveReadAAStringSet.R

```
# function for extract protein sequence from faa file by protein_id as a query 
# x = a vector of path for .faa files, like c(path/to/protein_seq_1.faa, path/to/protein_seq_2.faa)
# y = a vector of protein_id, like c(WP_xxxxxxxxx.1, WP_yyyyyyyyy.2)


# output tibble: 1 × 2
# protein_id seq                                
# <chr>      <chr>                              
# 1 WP_xxxxxxxxx.1 MAAAAAAAAAAAAAAAAAAAAAAAAA

# function ----------------------------------------------------------------

fn_SelectiveReadAAStringSet <- function(x, y){
  # load faa file as AAStringSet 
  tmp_AAset <- readAAStringSet(x)
  
  # extract protein names used in the faa file
  tmp_AAname <- tibble(name = names(tmp_AAset)) %>%
    
    # extract protein_id from the protein name and select proteins by protein_id
    mutate(protein_id = str_extract(name, '\\w+.\\w+')) %>% 
    filter(protein_id %in% y)
  
  # prep output fie which constitute protein_id and sequence
  output <- tibble(
    name=names(tmp_AAset[tmp_AAname$name]),
    seq=as.character(tmp_AAset[tmp_AAname$name])
  ) %>%
    transmute(
      protein_id = str_extract(name, '\\w+.\\w+'),
      seq
    ) %>%
    distinct()
}
```
