## Supplementary Code for "A role for the S4-domain containing protein YlmH in ribosome-associated quality control in *Bacillus subtilis*": fn_write_Fasta.R

```
fn_write_Fasta <- function(data, output_fasta_name, seq_name, sequence){
  fastaLines = c()
  for (rowNum in 1:nrow(data)){
    fastaLines = c(fastaLines, as.character(paste(">", data[rowNum,seq_name], sep = "")))
    fastaLines = c(fastaLines,as.character(data[rowNum,sequence]))
  }
  fileConn<-file(output_fasta_name)
  writeLines(fastaLines, fileConn)
  close(fileConn)
}
```
