## Supplementary Code for "A role for the S4-domain containing protein YlmH in ribosome-associated quality control in *Bacillus subtilis*": hmmsearch.sh

```
#!/bin/zsh

cd path/to/directory

# path
input_proteome_list="target/hmm_target_list.txt"
proteome_list=($(cat "$input_proteome_list"))

for file_hmm in input/*.hmm; do
	# extract file name
	echo start ${filename_hmm}
	date "+%Y.%m.%d.%W %H:%M:%S"
	TIME_A=`date +%s`   #A

	filename_hmm=${file_hmm##*/}
	# make output folders
	mkdir -p output/${filename_hmm%.*}/tblout/
	mkdir -p output/${filename_hmm%.*}/mainout/
	outdir_tbl=output/${filename_hmm%.*}/tblout/
	outdir_main=output/${filename_hmm%.*}/mainout/
# hmm search
for file_prt in "${proteome_list[@]}"; do
    
	# extract genome_id
    filename_prt=${file_prt##*/}
	dirname_prt=${file_prt%/*}
	genome_id=${dirname_prt##*/}


	output_tbl=${outdir_tbl}/${genome_id}_t.txt
	output_main=${outdir_main}/${genome_id}_m.txt

    echo target_genome: ${genome_id}
	
	#hmmsearch
    hmmsearch --noali --tblout ${output_tbl} -o ${output_main} ${file_hmm} ${file_prt}

done
TIME_B=`date +%s`   #B
PT=`expr ${TIME_B} - ${TIME_A}`
H=`expr ${PT} / 3600`
PT=`expr ${PT} % 3600`
M=`expr ${PT} / 60`
S=`expr ${PT} % 60`

echo "search_completed"
date "+%Y.%m.%d.%W %H:%M:%S"
echo "${H}:${M}:${S}"

echo "merging"
for file in ${outdir_tbl}/*.txt; do
    echo "Contents of $file:" >> output/${filename_hmm%.*}/merged_result.txt
    cat "$file" >> output/${filename_hmm%.*}/merged_result.txt
done
Rscript ../sc_HMM_merge.R ${filename_hmm%.*}
echo "merging_completed"

done
```
