## Supplementary Code for "A role for the S4-domain containing protein YlmH in ribosome-associated quality control in *Bacillus subtilis*": mafft_fftns_RqcH.sh

```
#!/bin/zsh

# alignment by mafft
# fftns input_file > output_file


cd path/to/directory
mkdir -p ./mafft_raw/fftns

for f in ./fasta_raw/RqcH*.fasta; do
    echo ${f}
    filename=${f##*/}
    echo ./mafft_raw/fftns/${filename%%.*}_fftns.txt
    fftns --thread -1 ${f} > ./mafft_raw/fftns/${filename%%.*}_fftns.txt
done
```
