## Supplementary Code for "A role for the S4-domain containing protein YlmH in ribosome-associated quality control in *Bacillus subtilis*": sc_HMM_merge.R

```
# input -------------------------------------------------------------------

args <- commandArgs(trailingOnly = T)
input <- args[1]

# package -----------------------------------------------------------------


library(tidyverse)


# merge -------------------------------------------------------------------

input_files <- str_c('output/',input,'/merged_result.txt') %>% print()
output <- str_c('output/', input,'/',input,'.csv') %>% print()

# Read the entire file
all_text <- readLines(input_files)


tibble(raw = str_subset(all_text, '^#', negate = T)) %>% 
  transmute(
    file = if_else(str_detect(raw, '^Contents of '), str_extract(raw, 'GC[AF]_\\d+.\\d'), NA),
    data = if_else(str_detect(raw, '^Contents of '), NA, raw),
  ) %>% 
  fill(file, .direction = 'down') %>% 
  filter(!is.na(data)) %>% 
  mutate(
    data = str_split(data, '[[:blank:]]{1,}')
  ) %>% 
  unnest_wider(data, names_sep = '_') %>% 
  dplyr::rename(
    target_name = data_1,
    target_accession = data_2,
    query_name = data_3,
    query_accession = data_4,
    E_fullseq = data_5,
    score_fullseq = data_6,
    bias_fullseq = data_7,
    E_best1 = data_8,
    score_best1 = data_9,
    bias_best1 = data_10,
    exp = data_11,
    reg = data_12,
    clu = data_13,
    ov = data_14,
    env = data_15,
    dom = data_16,
    rep = data_17,
    inc = data_18
  ) %>%  
  mutate(
    description_of_target=do.call(paste, c(select(., starts_with('data_')), sep = " ")) %>% str_remove_all('( NA)+$'),
  ) %>% 
  select(-file, -starts_with('data_'), genome = file) %>%
  write_csv(output)
```
