## Supplementary Code for "A role for the S4-domain containing protein YlmH in ribosome-associated quality control in *Bacillus subtilis*": summarize_merging.R

```
# YlmH analysis

# input -------------------------------------------------------------------

input_YlmH <- 'output/YlmH.csv'
input_RqcH <- 'output/RqcH.csv'
input_RqcP_Hsp15 <- 'output/RqcP_Hsp15_4216.csv'
input_SmpB <- 'output/SmpB.csv'

# path to a list of genome id
input_genome_list <- 'genome_list/genome_id_for_analysis.csv'

output <- 'output/list_fin_4216.csv'

# package -----------------------------------------------------------------

library(tidyverse)

# function ----------------------------------------------------------------


# load --------------------------------------------------------------------

# load genome id list
df_genome_list <- read_csv(input_genome_list) %>% print()

# load each csv
df_proteins <- c(input_YlmH, input_RqcH, input_RqcP_Hsp15, input_SmpB) %>% 
  purrr::map(readr::read_csv)


# formatting --------------------------------------------------------------


# format proteins
df_all <- df_proteins %>% 
  purrr::map(
    dplyr::select,
    protein_id,
    genome_id,
    protein
    ) %>% 
  purrr::map(
    pivot_wider,
    id_cols = genome_id,
    names_from = protein,
    values_from = protein_id
  )
  
df_YlmH <- df_all[[1]] %>% 
  unnest_wider(YlmH, names_sep = '_') %>% 
  rowwise() %>% 
  mutate(
    YlmH_2=str_replace_na(YlmH_2, 'NA'),
    YlmH_3=str_replace_na(YlmH_3, 'NA')
  ) %>% 
  transmute(
    genome_id, 
    YlmH=str_c(YlmH_1, ";", YlmH_2, ";", YlmH_3) %>%
      str_remove_all(';$|;NA$|;NA;NA$') 
    ) %>% 
  print()
  
df_RqcH <- df_all[[2]] %>% 
  unnest_wider(RqcH, names_sep = '_') %>% 
  rowwise() %>% 
  mutate(
    RqcH_2=str_replace_na(RqcH_2, 'NA'),
    RqcH_3=str_replace_na(RqcH_3, 'NA')
  ) %>% 
  transmute(
    genome_id, 
    RqcH=str_c(RqcH_1, ";", RqcH_2, ";", RqcH_3) %>%
      str_remove_all(';$|;NA$|;NA;NA$') 
    ) %>% 
  print()

df_RqcP_Hsp15 <- df_all[[3]] %>% 
  unnest_wider(c(RqcP, Hsp15), names_sep = '_') %>% 
  mutate(
    RqcP_1=str_replace_na(RqcP_1, 'NA'),
    RqcP_2=str_replace_na(RqcP_2, 'NA'),
    RqcP_3=str_replace_na(RqcP_3, 'NA'),
    Hsp15_1=str_replace_na(Hsp15_1, 'NA'),
    Hsp15_2=str_replace_na(Hsp15_2, 'NA'),
    # Hsp15_short_1=str_replace_na(Hsp15_short_1, 'NA'),
    # Hsp15_short_2=str_replace_na(Hsp15_short_2, 'NA')
  ) %>% 
  rowwise() %>% 
  transmute(
    genome_id,
    RqcP=str_c(RqcP_1, ";", RqcP_2, ";", RqcP_3) %>% 
      str_remove_all('NA$|;NA$|;NA;NA$|;NA;NA;NA$'),
    Hsp15=str_c(Hsp15_1, ";", Hsp15_2) %>% 
      str_remove_all('NA$|;NA$|;NA;NA$|;NA;NA;NA$'),
    # Hsp15_short=str_c(Hsp15_short_1, ";", Hsp15_short_2) %>% 
    #   str_remove_all('NA$|;NA$|;NA;NA$|;NA;NA;NA$') 
    ) %>% 
  transmute(
    genome_id,
    `Hsp15/RqcP` = str_c(RqcP, ";", Hsp15) %>% 
      str_remove_all(';$|;NA$|;NA;NA$|NA;'),
    RqcP = RqcP
  ) %>% 
  print()

df_SmpB <- df_all[[4]] %>% 
  unnest_wider(SmpB, names_sep = '_') %>% 
  rowwise() %>% 
  mutate(
    SmpB_2=str_replace_na(SmpB_2, 'NA'),
    SmpB_3=str_replace_na(SmpB_3, 'NA')
  ) %>% 
  transmute(
    genome_id, 
    SmpB=str_c(SmpB_1, ";", SmpB_2, ";", SmpB_3) %>%
      str_remove_all(';$|;NA$|;NA;NA$')
  ) %>% 
  print()


# output ------------------------------------------------------------------


df_merged <- df_genome_list %>% 
  left_join(df_SmpB, by = 'genome_id') %>% 
  left_join(df_YlmH, by = 'genome_id') %>% 
  left_join(df_RqcH, by = 'genome_id') %>% 
  left_join(df_RqcP_Hsp15, by = 'genome_id') %>% 
  arrange(gtdb_taxonomy) %>% 
  select(-c(genome_id_database, genome_database, phylum:species)) %>% 
  write_csv(output)
```
