## Supplementary Code for "A role for the S4-domain containing protein YlmH in ribosome-associated quality control in *Bacillus subtilis*": summarize_stats_combination.R

```
# input -------------------------------------------------------------------

input_list <- '/output/list_fin_4216.csv'
input_phylum_num <- '/phylum_numbering.csv'

output1 <- 'StatsTable1_CountGenomesRepPhyla_full_4216.csv'
output2 <- 'StatsTable2_HaveProteins_full_4216.csv'
output3 <- 'StatsTable3_RqcH_vs_RqcP_full_4216.csv'
output4 <- 'StatsTable4_RqcH_vs_YlmH_full_4216.csv'
output5 <- 'StatsTable5_RqcHPY_full_4216.csv'
output_heatmap <- 'Heatmap_full_4216.pdf'

# package -----------------------------------------------------------------


library(tidyverse)
library(stringi)


# load --------------------------------------------------------------------


df <- read_csv(input_list)
df_phylum_numbering <- read_csv(input_phylum_num)

df2 <- df %>% 
  rowwise() %>% 
  mutate(
    tmp = list(str_split(gtdb_taxonomy, ';')),
    phylum = unlist(tmp)[2],
  ) %>% 
  mutate(
    Hsp15 = if_else(!is.na(`Hsp15/RqcP`) & is.na(RqcP), `Hsp15/RqcP`, NA_character_),
    phylum=if_else(str_detect(phylum, 'p__Bacillota'), 'p__Bacillota', phylum),
    phylum=if_else(str_detect(phylum, 'p__Desulfobacterota'), 'p__Desulfobacterota', phylum)
  ) %>% 
  ungroup() %>% 
  select(-tmp, -`Hsp15/RqcP`)

# select representative phyla
v_sel_phyla <- df2 %>% 
  dplyr::count(phylum, sort = T) %>% 
  dplyr::left_join(df_phylum_numbering) %>% 
  dplyr::arrange(phylum_numbering) %>% 
  dplyr::select(-n) %>% print()


# stats_Genome_count ------------------------------------------------------

df2 %>% 
  distinct(genome_id) %>% 
  count()

# stats_Genomes/phyla -----------------------------------------------------


df2 %>% 
  count(phylum, sort = T) %>% 
  dplyr::mutate(phylum_numbering=row_number()) %>% 
  write_csv(output1)


# stats_HaveProtein -------------------------------------------------------

# SmpB
HaveSmpB <- df2 %>% 
  right_join(v_sel_phyla, by = 'phylum') %>% 
  count(
    phylum_numbering,
    phylum,
    HaveProtein=!is.na(SmpB),
    sort = T
  ) %>%
  tidyr::complete(nesting(phylum, phylum_numbering), HaveProtein) %>%
  dplyr::mutate(n=if_else(is.na(n), 0, n)) %>% # NA->0, 
  dplyr::rename(n_HaveSmpB=n) %>% 
  dplyr::arrange(phylum_numbering, -HaveProtein) %>% print()

# RqcH
HaveRqcH <- df2 %>% 
  right_join(v_sel_phyla, by = 'phylum') %>% 
  count(
    phylum_numbering,
    phylum,
    HaveProtein=!is.na(RqcH),
    sort = T
  ) %>% 
  tidyr::complete(nesting(phylum, phylum_numbering), HaveProtein) %>%
  dplyr::mutate(n=if_else(is.na(n), 0, n)) %>% # NA->0, 
  dplyr::rename(n_HaveRqcH=n) %>% 
  dplyr::arrange(phylum_numbering, -HaveProtein) %>% print()

# YlmH
HaveYlmH <- df2 %>% 
  right_join(v_sel_phyla, by = 'phylum') %>% 
  count(
    phylum_numbering,
    phylum,
    HaveProtein=!is.na(YlmH),
    sort = T
  ) %>% 
  tidyr::complete(nesting(phylum, phylum_numbering), HaveProtein) %>%
  dplyr::mutate(n=if_else(is.na(n), 0, n)) %>% # NA->0, 
  dplyr::rename(n_HaveYlmH=n) %>% 
  dplyr::arrange(phylum_numbering, -HaveProtein) %>% print()

# RqcP
HaveRqcP <- df2 %>% 
  right_join(v_sel_phyla, by = 'phylum') %>% 
  count(
    phylum_numbering,
    phylum,
    HaveProtein=!is.na(RqcP),
    sort = T
  ) %>% 
  tidyr::complete(nesting(phylum, phylum_numbering), HaveProtein) %>%
  dplyr::mutate(n=if_else(is.na(n), 0, n)) %>% # NA->0, 
  dplyr::rename(n_HaveRqcP=n) %>% 
  dplyr::arrange(phylum_numbering, -HaveProtein) %>% print()

# Hsp15
HaveHsp15 <- df2 %>% 
  right_join(v_sel_phyla, by = 'phylum') %>% 
  count(
    phylum_numbering,
    phylum,
    HaveProtein=!is.na(Hsp15),
    sort = T
  ) %>% 
  tidyr::complete(nesting(phylum, phylum_numbering), HaveProtein) %>%
  dplyr::mutate(n=if_else(is.na(n), 0, n)) %>% # NA->0, 
  dplyr::rename(n_HaveHsp15=n) %>% 
  dplyr::arrange(phylum_numbering, -HaveProtein) %>% print()

HaveProtein <- HaveSmpB %>% 
  left_join(HaveRqcH, by = c('phylum', 'HaveProtein', 'phylum_numbering')) %>% 
  left_join(HaveRqcP, by = c('phylum', 'HaveProtein', 'phylum_numbering')) %>% 
  left_join(HaveHsp15, by = c('phylum', 'HaveProtein', 'phylum_numbering')) %>% 
  left_join(HaveYlmH, by = c('phylum', 'HaveProtein', 'phylum_numbering')) %>% 
  select(phylum_numbering, everything()) %>% 
  group_by(phylum) %>% 
  mutate(
    `%HaveSmpB` = n_HaveSmpB/sum(n_HaveSmpB),
    `%HaveRqcH` = n_HaveRqcH/sum(n_HaveRqcH),
    `%HaveYlmH` = n_HaveYlmH/sum(n_HaveYlmH),
    `%HaveRqcP` = n_HaveRqcP/sum(n_HaveRqcP),
    `%HaveHsp15` = n_HaveHsp15/sum(n_HaveHsp15)
  )
HaveProtein %>% 
  write_csv(output2)


# Stats_ProteinCombination ------------------------------------------------

HaveSmpB <- df2 %>% 
  right_join(v_sel_phyla, by = 'phylum') %>% 
  count(
    phylum_numbering,
    phylum,
    HaveProtein=!is.na(SmpB),
    sort = T
  ) %>%
  tidyr::complete(nesting(phylum, phylum_numbering), HaveProtein) %>%
  dplyr::mutate(n=if_else(is.na(n), 0, n)) %>% # NA->0, 
  dplyr::rename(n_HaveSmpB=n) %>% 
  dplyr::arrange(phylum_numbering, -HaveProtein) %>% print()


# Combination
RqcH_RqcP_B <- df2 %>% 
  right_join(v_sel_phyla, by = 'phylum') %>% 
  count(
    phylum_numbering,
    phylum,
    HaveProteins=!is.na(`RqcP`) & !is.na(`RqcH`),
    sort = T
  ) %>% 
  tidyr::complete(nesting(phylum, phylum_numbering), HaveProteins) %>%
  dplyr::mutate(n=if_else(is.na(n), 0, n)) %>% # NA->0, 
  dplyr::rename(`n_Both RqcH & RqcP`=n) %>% 
  dplyr::arrange(phylum_numbering, -HaveProteins) %>% print()

RqcH_RqcP_E <- df2 %>% 
  right_join(v_sel_phyla, by = 'phylum') %>% 
  count(
    phylum_numbering,
    phylum,
    HaveProteins=(!is.na(`RqcP`) & is.na(`RqcH`)) | (is.na(`RqcP`) & !is.na(`RqcH`)),
    sort = T
  ) %>% 
  tidyr::complete(nesting(phylum, phylum_numbering), HaveProteins) %>%
  dplyr::mutate(n=if_else(is.na(n), 0, n)) %>% # NA->0, 
  dplyr::rename(`n_either RqcH or RqcP`=n) %>% 
  dplyr::arrange(phylum_numbering, -HaveProteins) %>% print()

RqcH_RqcP_N <- df2 %>% 
  right_join(v_sel_phyla, by = 'phylum') %>% 
  count(
    phylum_numbering,
    phylum,
    HaveProteins=is.na(`RqcP`) & is.na(`RqcH`),
    sort = T
  ) %>% 
  tidyr::complete(nesting(phylum, phylum_numbering), HaveProteins) %>%
  dplyr::mutate(n=if_else(is.na(n), 0, n)) %>% # NA->0, 
  dplyr::rename(`n_None`=n) %>% 
  dplyr::arrange(phylum_numbering, -HaveProteins) %>% print()

RqcH_RqcP <- RqcH_RqcP_B %>% 
  left_join(RqcH_RqcP_E, by = c('phylum', 'HaveProteins', 'phylum_numbering')) %>% 
  left_join(RqcH_RqcP_N, by = c('phylum', 'HaveProteins', 'phylum_numbering')) %>% 
  select(phylum_numbering, everything()) %>% 
  group_by(phylum) %>% 
  mutate(
    `%BOTH` = `n_Both RqcH & RqcP`/sum(`n_Both RqcH & RqcP`),
    `%EITHER` = `n_either RqcH or RqcP`/sum(`n_either RqcH or RqcP`),
    `%NON` = n_None/sum(n_None)
  )


RqcH_RqcP %>% 
  write_csv(output3)


RqcH_YlmH_B <- df2 %>% 
  right_join(v_sel_phyla, by = 'phylum') %>% 
  count(
    phylum_numbering,
    phylum,
    HaveProteins=!is.na(`YlmH`) & !is.na(`RqcH`),
    sort = T
  ) %>% 
  tidyr::complete(nesting(phylum, phylum_numbering), HaveProteins) %>%
  dplyr::mutate(n=if_else(is.na(n), 0, n)) %>% # NA->0, 
  dplyr::rename(`n_Both RqcH & YlmH`=n) %>% 
  dplyr::arrange(phylum_numbering, -HaveProteins) %>% print()

RqcH_YlmH_E <- df2 %>% 
  right_join(v_sel_phyla, by = 'phylum') %>% 
  count(
    phylum_numbering,
    phylum,
    HaveProteins=(!is.na(`YlmH`) & is.na(`RqcH`)) | (is.na(`YlmH`) & !is.na(`RqcH`)),
    sort = T
  ) %>% 
  tidyr::complete(nesting(phylum, phylum_numbering), HaveProteins) %>%
  dplyr::mutate(n=if_else(is.na(n), 0, n)) %>% # NA->0, 
  dplyr::rename(`n_either RqcH or YlmH`=n) %>% 
  dplyr::arrange(phylum_numbering, -HaveProteins) %>% print()

RqcH_YlmH_N <- df2 %>% 
  right_join(v_sel_phyla, by = 'phylum') %>% 
  count(
    phylum_numbering,
    phylum,
    HaveProteins=is.na(`YlmH`) & is.na(`RqcH`),
    sort = T
  ) %>% 
  tidyr::complete(nesting(phylum, phylum_numbering), HaveProteins) %>%
  dplyr::mutate(n=if_else(is.na(n), 0, n)) %>% # NA->0, 
  dplyr::rename(`n_None`=n) %>% 
  dplyr::arrange(phylum_numbering, -HaveProteins) %>% print()

RqcH_YlmH <- RqcH_YlmH_B %>% 
  left_join(RqcH_YlmH_E, by = c('phylum', 'HaveProteins', 'phylum_numbering')) %>% 
  left_join(RqcH_YlmH_N, by = c('phylum', 'HaveProteins', 'phylum_numbering')) %>% 
  select(phylum_numbering, everything()) %>% 
  group_by(phylum) %>% 
  mutate(
    `%BOTH` = `n_Both RqcH & YlmH`/sum(`n_Both RqcH & YlmH`),
    `%EITHER` = `n_either RqcH or YlmH`/sum(`n_either RqcH or YlmH`),
    `%NON` = n_None/sum(n_None)
  )


RqcH_YlmH %>% 
  write_csv(output4)


RqcHPY_ALL <- df2 %>% 
  right_join(v_sel_phyla, by = 'phylum') %>% 
  count(
    phylum_numbering,
    phylum,
    HaveProteins=!is.na(`RqcH`) & !is.na(`RqcP`) & !is.na(`YlmH`) & is.na(`Hsp15`),
    sort = T
  ) %>% 
  tidyr::complete(nesting(phylum, phylum_numbering), HaveProteins) %>%
  dplyr::mutate(n=if_else(is.na(n), 0, n)) %>% # NA->0, 
  dplyr::rename(`n_RqcH_RqcP_YlmH`=n) %>% 
  dplyr::arrange(phylum_numbering, -HaveProteins) %>% print()

RqcHP <- df2 %>% 
  right_join(v_sel_phyla, by = 'phylum') %>% 
  count(
    phylum_numbering,
    phylum,
    HaveProteins=!is.na(`RqcH`) & !is.na(`RqcP`) & is.na(`YlmH`) & is.na(`Hsp15`),
    sort = T
  ) %>% 
  tidyr::complete(nesting(phylum, phylum_numbering), HaveProteins) %>%
  dplyr::mutate(n=if_else(is.na(n), 0, n)) %>% # NA->0, 
  dplyr::rename(`n_RqcH_RqcP`=n) %>% 
  dplyr::arrange(phylum_numbering, -HaveProteins) %>% print()

RqcHY <- df2 %>% 
  right_join(v_sel_phyla, by = 'phylum') %>% 
  count(
    phylum_numbering,
    phylum,
    HaveProteins=!is.na(`RqcH`) & is.na(`RqcP`) & !is.na(`YlmH`) & is.na(`Hsp15`),
    sort = T
  ) %>% 
  tidyr::complete(nesting(phylum, phylum_numbering), HaveProteins) %>%
  dplyr::mutate(n=if_else(is.na(n), 0, n)) %>% # NA->0, 
  dplyr::rename(`n_RqcH_YlmH`=n) %>% 
  dplyr::arrange(phylum_numbering, -HaveProteins) %>% print()

RqcH15 <- df2 %>% 
  right_join(v_sel_phyla, by = 'phylum') %>% 
  count(
    phylum_numbering,
    phylum,
    HaveProteins=!is.na(`RqcH`) & is.na(`RqcP`) & is.na(`YlmH`) & !is.na(`Hsp15`),
    sort = T
  ) %>% 
  tidyr::complete(nesting(phylum, phylum_numbering), HaveProteins) %>%
  dplyr::mutate(n=if_else(is.na(n), 0, n)) %>% # NA->0, 
  dplyr::rename(`n_RqcH_Hsp15`=n) %>% 
  dplyr::arrange(phylum_numbering, -HaveProteins) %>% print()

# only
RqcH_only <- df2 %>% 
  right_join(v_sel_phyla, by = 'phylum') %>% 
  count(
    phylum_numbering,
    phylum,
    HaveProteins=!is.na(`RqcH`) & is.na(`RqcP`) & is.na(`YlmH`) & is.na(`Hsp15`),
    sort = T
  ) %>% 
  tidyr::complete(nesting(phylum, phylum_numbering), HaveProteins) %>%
  dplyr::mutate(n=if_else(is.na(n), 0, n)) %>% # NA->0, 
  dplyr::rename(`n_RqcH_only`=n) %>% 
  dplyr::arrange(phylum_numbering, -HaveProteins) %>% print()

RqcP_only <- df2 %>% 
  right_join(v_sel_phyla, by = 'phylum') %>% 
  count(
    phylum_numbering,
    phylum,
    HaveProteins=is.na(`RqcH`) & !is.na(`RqcP`) & is.na(`YlmH`) & is.na(`Hsp15`),
    sort = T
  ) %>% 
  tidyr::complete(nesting(phylum, phylum_numbering), HaveProteins) %>%
  dplyr::mutate(n=if_else(is.na(n), 0, n)) %>% # NA->0, 
  dplyr::rename(`n_RqcP_only`=n) %>% 
  dplyr::arrange(phylum_numbering, -HaveProteins) %>% print()

YlmH_only <- df2 %>% 
  right_join(v_sel_phyla, by = 'phylum') %>% 
  count(
    phylum_numbering,
    phylum,
    HaveProteins=is.na(`RqcH`) & is.na(`RqcP`) & !is.na(`YlmH`) & is.na(`Hsp15`),
    sort = T
  ) %>% 
  tidyr::complete(nesting(phylum, phylum_numbering), HaveProteins) %>%
  dplyr::mutate(n=if_else(is.na(n), 0, n)) %>% # NA->0, 
  dplyr::rename(`n_YlmH_only`=n) %>% 
  dplyr::arrange(phylum_numbering, -HaveProteins) %>% print()

Hsp15_only <- df2 %>% 
  right_join(v_sel_phyla, by = 'phylum') %>% 
  count(
    phylum_numbering,
    phylum,
    HaveProteins=is.na(`RqcH`) & is.na(`RqcP`) & is.na(`YlmH`) & !is.na(`Hsp15`),
    sort = T
  ) %>% 
  tidyr::complete(nesting(phylum, phylum_numbering), HaveProteins) %>%
  dplyr::mutate(n=if_else(is.na(n), 0, n)) %>% # NA->0, 
  dplyr::rename(`n_Hsp15_only`=n) %>% 
  dplyr::arrange(phylum_numbering, -HaveProteins) %>% print()

SmpB_only <- df2 %>% 
  right_join(v_sel_phyla, by = 'phylum') %>% 
  count(
    phylum_numbering,
    phylum,
    HaveProteins=is.na(`RqcH`) & is.na(`RqcP`) & is.na(`YlmH`) & is.na(`Hsp15`) & !is.na(`SmpB`),
    sort = T
  ) %>% 
  tidyr::complete(nesting(phylum, phylum_numbering), HaveProteins) %>%
  dplyr::mutate(n=if_else(is.na(n), 0, n)) %>% # NA->0, 
  dplyr::rename(`n_SmpB_only`=n) %>% 
  dplyr::arrange(phylum_numbering, -HaveProteins) %>% print()

ND <- df2 %>% 
  right_join(v_sel_phyla, by = 'phylum') %>% 
  count(
    phylum_numbering,
    phylum,
    HaveProteins=is.na(`RqcH`) & is.na(`RqcP`) & is.na(`YlmH`) & is.na(`Hsp15`) & is.na(`SmpB`),
    sort = T
  ) %>% 
  tidyr::complete(nesting(phylum, phylum_numbering), HaveProteins) %>%
  dplyr::mutate(n=if_else(is.na(n), 0, n)) %>% # NA->0, 
  dplyr::rename(`n_NotDetected`=n) %>% 
  dplyr::arrange(phylum_numbering, -HaveProteins) %>% print()

# merge
RqcHPY <- RqcHPY_ALL %>% 
  left_join(
    df2 %>% 
      count(phylum, sort = T)
  ) %>% 
  left_join(RqcHP, by = c('phylum', 'HaveProteins', 'phylum_numbering')) %>% 
  left_join(RqcHY, by = c('phylum', 'HaveProteins', 'phylum_numbering')) %>%
  left_join(RqcH15, by = c('phylum', 'HaveProteins', 'phylum_numbering')) %>% 
  left_join(RqcH_only, by = c('phylum', 'HaveProteins', 'phylum_numbering')) %>% 
  left_join(RqcP_only, by = c('phylum', 'HaveProteins', 'phylum_numbering')) %>% 
  left_join(YlmH_only, by = c('phylum', 'HaveProteins', 'phylum_numbering')) %>%
  left_join(Hsp15_only, by = c('phylum', 'HaveProteins', 'phylum_numbering')) %>%
  left_join(SmpB_only, by = c('phylum', 'HaveProteins', 'phylum_numbering')) %>%
  left_join(ND, by = c('phylum', 'HaveProteins', 'phylum_numbering')) %>% 
  select(phylum_numbering, phylum, total_genomes = n, everything()) %>% 
  group_by(phylum) %>% 
  mutate(
    `%RqcH_RqcP_YlmH` = `n_RqcH_RqcP_YlmH`/sum(`n_RqcH_RqcP_YlmH`),
    `%RqcH_RqcP` = `n_RqcH_RqcP`/sum(`n_RqcH_RqcP`),
    `%RqcH_YlmH` = `n_RqcH_YlmH`/sum(`n_RqcH_YlmH`),
    `%RqcH_Hsp15` = `n_RqcH_Hsp15`/sum(`n_RqcH_Hsp15`),
    `%onlyRqcH` = `n_RqcH_only`/sum(`n_RqcH_only`),
    `%onlyRqcP` = `n_RqcP_only`/sum(`n_RqcP_only`),
    `%onlyYlmH` = `n_YlmH_only`/sum(`n_YlmH_only`),
    `%onlyHsp15` = `n_Hsp15_only`/sum(`n_Hsp15_only`),
    `%onlySmpB` = `n_SmpB_only`/sum(`n_SmpB_only`),
    `%NotDetected` = `n_NotDetected`/sum(`n_NotDetected`)
  ) %>% 
  ungroup() %>% 
  print()

RqcHPY %>% 
  write_csv(output5)

v_name_comb <- tibble(
  v_order = c(1:9),
  v_name = c("%RqcH_RqcP_YlmH", "%RqcH_RqcP", "%RqcH_YlmH", "%RqcH_Hsp15", "%onlyRqcH", "%onlyRqcP", "%onlyYlmH", "%onlyHsp15", "%onlySmpB")
)

df_plot <- RqcHPY %>%
  filter(HaveProteins == T) %>% 
  filter(
    if_any(
      .cols = all_of(v_name_comb$v_name[1:8]),
      .fns = \(x) x >=0.33
    ) | (`%onlySmpB`	> 0.33 & total_genomes >= 1000) # as representatives
  ) %>%
  mutate(phylum = str_remove(phylum, "p__")) %>%
  dplyr::select(
    phylum_numbering, phylum, starts_with('%'), #total_genomes
  ) %>% 
  pivot_longer(cols = starts_with("%"), names_to = 'v_name', values_to = 'ratio') %>% 
  left_join(v_name_comb, by = 'v_name') 
p1 <- df_plot %>% 
  ggplot(aes(reorder(v_name, v_order), reorder(phylum, -phylum_numbering), fill=ratio)) +
  geom_tile(color = 'grey60') +
  scale_fill_gradient(low = 'white', high = '#ff4b00') +
  geom_text(aes(label = round(ratio, 2) * 100), size = 3) +
  guides(fill = guide_legend(reverse = T)) +
  #theme_classic() +
  theme(
    axis.title.x = element_blank(),
    axis.title.y = element_blank(),
    axis.text.x = element_text(size = 8),
    axis.text.y = element_text(size = 8)
  )
p1
ggsave(file = output_heatmap, plot = p1, width = 6, height = 6)


RqcHPY %>% 
  mutate(phylum = str_remove(phylum, "p__")) %>%  
  filter(HaveProteins == T) %>% 
  filter(phylum %in% df_plot$phylum) %>% 
  select(total_genomes) %>% 
  print(n=Inf)
```
