## Supplementary Code for "A role for the S4-domain containing protein YlmH in ribosome-associated quality control in *Bacillus subtilis*": tree_1_prune.R

```
# to prepare genome list

# input -------------------------------------------------------------------

input_genome_list <- 'genome_list/genome_id_for_analysis.csv'

# tree infor
input_tree <- 'input/gtdb/bac120.tree'
input_tree_info <- 'input/gtdb/bac120_taxonomy.tsv'

output_tree_pruned <- 'tree/1_bac/b120_r214_52895_pruned'

# package -----------------------------------------------------------------

library(tidyverse)
library(phytools)

# function ----------------------------------------------------------------


# load --------------------------------------------------------------------

df_genome_list <- read_csv(input_genome_list)

tree_raw <- phytools::read.newick(input_tree)

# tree formatting ---------------------------------------------------------

tree_pruned <- ape::drop.tip(
  tree_raw,
  setdiff(
    tree_raw$tip.label,
    df_genome_list$genoem_id_database
  )
)


# output_tree -------------------------------------------------------------

# pruned tree
tree_pruned %>% 
  writeNexus(output_tree_pruned)
```
