## Supplementary Code for "A role for the S4-domain containing protein YlmH in ribosome-associated quality control in *Bacillus subtilis*": tree_2_split.R

```
# to prepare genome list

# input -------------------------------------------------------------------

input_genome_list <- 'genome_list/genome_id_for_analysis.csv'

# tree infor
input_tree_pruned <- 'tree/1_bac/b120_r214_52895_pruned.tree'

output_tree_phylum <- 'tree/2_tree_split/1_phylum/'
output_tree_class <- 'tree/2_tree_split/2_class/'
output_tree_order <- 'tree/2_tree_split/3_order//'


# package -----------------------------------------------------------------

library(tidyverse)
library(phytools)

# function ----------------------------------------------------------------


# load --------------------------------------------------------------------

df_genome_list <- read_csv(input_genome_list) %>% print()
tree_pruned <- phytools::readNexus(input_tree_pruned)


# formatting --------------------------------------------------------------

df_tree_info_f1 <- df_genome_list

# output, split by phyla --------------------------------------------------

# split the pruned tree by phylum
tree_pruned
v_phylum <- df_tree_info_f1 %>% count(phylum, sort = T) %>% pull(phylum)

set <- vector('list', 1)
for(i in seq_along(v_phylum)){
  set[[i]] <- df_tree_info_f1 %>% 
    filter(phylum == v_phylum[[i]])
  tree_pruned %>% 
    ape::drop.tip(setdiff(tree_pruned$tip.label, set[[i]]$genome_id_database)) %>% 
    writeNexus(
      str_c(
        output_tree_phylum,
        'b120_r214_52895_',
        v_phylum[[i]],
        '.tree'
      )
    )
}


# output, split by class --------------------------------------------------


# split the pruned tree by phylum
tree_pruned
v_class_1k <- df_tree_info_f1 %>% count(class, sort = T) %>% filter(n>1000) %>% pull(class) %>% print()

set <- vector('list', 1)
for(i in seq_along(v_class_1k)){
  set[[i]] <- df_tree_info_f1 %>% 
    filter(class == v_class_1k[[i]])
  tree_pruned %>% 
    ape::drop.tip(setdiff(tree_pruned$tip.label, set[[i]]$genome_id_database)) %>% 
    writeNexus(
      str_c(
        output_tree_class,
        'b120_r214_52895_',
        v_class_1k[[i]],
        '.tree'
      )
    )
}


# output, split by order --------------------------------------------------


# split the pruned tree by order
v_order_1k <- df_tree_info_f1 %>% count(order, sort = T) %>% filter(n>1000) %>% pull(order) %>% print()

set <- vector('list', 1)
for(i in seq_along(v_order_1k)){
  set[[i]] <- df_tree_info_f1 %>% 
    filter(order == v_order_1k[[i]])
  tree_pruned %>% 
    ape::drop.tip(setdiff(tree_pruned$tip.label, set[[i]]$genome_id_database)) %>% 
    writeNexus(
      str_c(
        output_tree_order,
        'b120_r214_52895_',
        v_order_1k[[i]],
        '.tree'
      )
    )
}


# output, split by family -------------------------------------------------
```
