## Supplementary Code for "A role for the S4-domain containing protein YlmH in ribosome-associated quality control in *Bacillus subtilis*": tree_3_iTol_lineage.R

```
# to prepare genome list

# input -------------------------------------------------------------------

# genome information
input_genome_list <- 'genome_list/genome_id_for_analysis.csv'

# color set
input_color_set <- 'tree/3_dataset_lineage/lineage_color_23Z28.xlsx'

# output
output_tree_ds_p <- 'tree/3_dataset_lineage/tree_ds_phylum_23Z28.txt'

# package -----------------------------------------------------------------

library(tidyverse)
library(readxl)


# function ----------------------------------------------------------------


# load --------------------------------------------------------------------

# load genome information
df_genome_list <- read_csv(input_genome_list) %>% print()

# load color dataframe
df_color_p <- read_excel(input_color_set, sheet = 'phylum') %>% print()
df_color_c <- read_excel(input_color_set, sheet = 'class') %>% print()
df_color_o <- read_excel(input_color_set, sheet = 'order') %>% print()
df_color_f <- read_excel(input_color_set, sheet = 'family') %>% print()
df_color_g <- read_excel(input_color_set, sheet = 'genus') %>% print()


# prep dataset, phylum ----------------------------------------------------

df_tree_color_p <- df_genome_list %>% 
  left_join(select(df_color_p, -n), by = 'phylum') %>% print()


df_dataset_keywords_p <- tibble(
  lines = c('DATASET_COLORSTRIP','SEPARATOR','DATASET_LABEL','COLOR,#005AFF','STRIP_WIDTH','MARGIN','DATA')) %>% 
  mutate(
    lines = case_when(
      lines == 'SEPARATOR' ~ str_c(lines, ' ', 'COMMA'),
      lines == 'DATASET_LABEL' ~ str_c(lines, ',', 'phylum'),
      lines == 'STRIP_WIDTH' ~ str_c(lines, ',', '600'),
      lines == 'MARGIN' ~ str_c(lines, ',', '-600'),
      !is.na(lines) ~ lines,
    )
  )

# prep iTol dataset
df_dataset_data_p <- df_tree_color_p %>% 
  select(genome_id_database, color) %>% 
  mutate(
    database_id_num = str_extract(genome_id_database, '\\d{9,9}')
  ) %>% 
  transmute(dataset_row = str_c('CONTAINS==',database_id_num,',',color)) %>% 
  filter(!is.na(dataset_row))

write_lines(
  df_dataset_keywords_p$lines,
  output_tree_ds_p
)

df_dataset_data_p %>% 
  write_delim(output_tree_ds_p, append = TRUE, col_names = FALSE)


# prep dataset, class -----------------------------------------------------


df_tree_color_c <- df_genome_list %>% 
  left_join(select(df_color_c, -n), by = c('phylum', 'class') ) %>% print()


df_dataset_keywords_c <- tibble(
  lines = c('DATASET_COLORSTRIP','SEPARATOR','DATASET_LABEL','COLOR,#005AFF','STRIP_WIDTH','MARGIN','DATA')) %>% 
  mutate(
    lines = case_when(
      lines == 'SEPARATOR' ~ str_c(lines, ' ', 'COMMA'),
      lines == 'DATASET_LABEL' ~ str_c(lines, ',', 'class'),
      lines == 'STRIP_WIDTH' ~ str_c(lines, ',', '600'),
      lines == 'MARGIN' ~ str_c(lines, ',', '-600'),
      !is.na(lines) ~ lines,
    )
  )

# prep iTol dataset
df_dataset_data_p <- df_tree_color_c %>% 
  select(genome_id_database, color) %>% 
  mutate(
    database_id_num = str_extract(genome_id_database, '\\d{9,9}')
  ) %>% 
  transmute(dataset_row = str_c('CONTAINS==',database_id_num,',',color)) %>% 
  filter(!is.na(dataset_row))

write_lines(
  df_dataset_keywords_c$lines,
  output_tree_ds_c
)

df_dataset_data_c %>% 
  write_delim(output_tree_ds_c, append = TRUE, col_names = FALSE)


# prep dataset, order -----------------------------------------------------


df_tree_color_o <- df_genome_list %>% 
  left_join(select(df_color_o, -n), by = c('phylum', 'class', 'order') ) %>% print()


df_dataset_keywords_o <- tibble(
  lines = c('DATASET_COLORSTRIP','SEPARATOR','DATASET_LABEL','COLOR,#005AFF','STRIP_WIDTH','MARGIN','DATA')) %>% 
  mutate(
    lines = case_when(
      lines == 'SEPARATOR' ~ str_c(lines, ' ', 'COMMA'),
      lines == 'DATASET_LABEL' ~ str_c(lines, ',', 'order'),
      lines == 'STRIP_WIDTH' ~ str_c(lines, ',', '600'),
      lines == 'MARGIN' ~ str_c(lines, ',', '-600'),
      !is.na(lines) ~ lines,
    )
  )

# prep iTol dataset
df_dataset_data_o <- df_tree_color_o %>% 
  select(genome_id_database, color) %>% 
  mutate(
    database_id_num = str_extract(genome_id_database, '\\d{9,9}')
  ) %>% 
  transmute(dataset_row = str_c('CONTAINS==',database_id_num,',',color)) %>% 
  filter(!is.na(dataset_row))

write_lines(
  df_dataset_keywords_o$lines,
  output_tree_ds_o
)

df_dataset_data_o %>% 
  write_delim(output_tree_ds_o, append = TRUE, col_names = FALSE)
```
