## Supplementary Code for "A role for the S4-domain containing protein YlmH in ribosome-associated quality control in *Bacillus subtilis*": A1_persing_genome.R

```
# Input -------------------------------------------------------------------

# genomic information
input_fna <- "reference/GCF_000009045.1_ASM904v1_genomic.fna"
input_gff <- 'reference/GCF_000009045.1_ASM904v1_genomic.gff'

# motif to find
input_motif <- "TA"

seqid <- "NC_000964.3"

# output name
output_1 <- "reference/TA_position_168.csv"
output_2 <- "reference/TA_annotation.csv"

# Package -----------------------------------------------------------------

library(Biostrings)
library(tidyverse)
library(fuzzyjoin)
library(rtracklayer)

packageVersion("Biostrings")
packageVersion("tidyverse")
packageVersion("fuzzyjoin")
packageVersion("rtracklayer")

# Script ------------------------------------------------------------------

# import genome sequence
seq_genome <- readDNAStringSet(input_fna, format="fasta") %>% print()


# search
hoge1 <- vmatchPattern(
  pattern = input_motif, 
  subject = seq_genome, 
  fixed = TRUE
)


# prep T-A list
df_TA <- tibble(
  TA_start = start(unlist(hoge1)),
  TA_end = end(unlist(hoge1)),
  seqid = seqid,
  strand_pos = 1,
  strand_neg = -1
) %>% 
  tidyr::pivot_longer(
    cols = c(strand_pos, strand_neg),
    values_to = "TA_str"
  )   %>% 
  dplyr::mutate(
    TA_pos = case_when(
      TA_str == 1 ~ TA_end,
      TA_str == -1 ~ TA_start
    ),
    TA_id = str_c("TA", TA_pos,"_", seqid)
  ) %>% 
  dplyr::select(TA_id, seqid, TA_pos, TA_start, TA_end, TA_str) %>% 
  dplyr::arrange(TA_pos) %>% 
  distinct() %>% 
  print()

df_TA %>% 
  write_csv(output_1)

# gff ---------------------------------------------------------------------

gff <- readGFF(input_gff) %>% as_tibble()

df_TA_annotation <- gff  %>% 
  filter(type != 'region') %>% 
  dplyr::mutate(
    strand = case_when(
      strand == "+" ~ 1,
      strand == "-" ~ -1
    ),
    gene_length = end - start + 1
  ) %>% 
  dplyr::rename(
    gene_start = start,
    gene_end = end,
    gene_strand = strand,
  ) %>% 
  fuzzyjoin::interval_left_join(
    df_TA %>% select(-seqid), 
    by = c(
      c("gene_start" = "TA_start"),
      c("gene_end" = "TA_end")
    ),
    minoverlap = 2
  ) %>% 
  distinct() %>% 
  print()

df_TA_annotation %>% 
  write_csv(output_2)
```
