## Supplementary Code for "A role for the S4-domain containing protein YlmH in ribosome-associated quality control in *Bacillus subtilis*": A2_calc_RPM.R

```
# perse_sam & calc RPM

# usage -------------------------------------------------------------------


# Put the following files in the same directory 
# sample_sheet.csv
# sample_compare_sheet.csv
# Place a SAM folder and put a sam file inside with the same name as the one entered in the sample_sheet.csv file. 


# Put the following files in one upper directory 
# p34_a3_TA_position_168.csv

# If there is no outputs folder, create it.

# Package -----------------------------------------------------------------

library(tidyverse)

# parse sample sheet and TA position list------------------------------------------------------

sample_sheet <- read_csv("sample_sheet.csv")
TA_pos_list <- read_csv('reference/TA_position_168.csv')

output_directory <- '4_analysis/q20/'

# function ----------------------------------------------------------------


# Function to read SAM files and make them tibble
fx_parseSAMfile <- function(x){
  # Create a vector with SAM column names
  SamRownames <- c("QNAME","FLAG","RNAME","POS","MAPQ","CIGAR","RNEXT","PNEXT","TLEN","SEQ","QUAL","Opt1","Opt2","Opt3","Opt4")
  
  # Read SAM files as tsv (tab separated values)
  # The three header lines starting with @ are not READ information, so they are not read.
  # Specify the column name vector you just created for the column name.
  Output <- read_tsv(x, skip = 3, col_names = SamRownames)
  # problems(), but the reason is that there are rows with insufficient number of columns. Since those rows are the case without Opt2~Opt4, NA can be put in them.
}


# function for changing SAM contents to TnSeq-specific dataframe
fx_SAM4Tnseq <- function(x){
  # Sequences derived from plasmid DNA are often intermingled.The sequences are later filtered out.
  pDNAseq <- "GCTTGATATCATTCTA"
  
  output <- x %>% 
    
    # Since QNAME has the structure ""read_id"-"read_count"", divide it into each of the following
    # After splitting, we want each variable type to be converted from string to numeric, so convert = TRUE
    tidyr::separate(QNAME, into = c("read_id", "read_count"), sep = "-", convert = TRUE) %>% 
    
    # Sequences derived from plasmid DNA are filtered out here.
    filter(SEQ != pDNAseq) %>% 
    
    # Reads that have not been mapped are removed by filter, identified by FLAG.
    # FLAG = 0 or 16: mapped 
    # FLAG = 4: unmapped
    dplyr::filter(FLAG != 4) %>% 
    
    # Set the genomic position of READ to the position of A in TA
    dplyr::mutate(TA_pos = case_when(
      
      # FLAG = 0: mapped to the + strand, so the last character in the read array should be at position A
      FLAG == 0 ~ as.double(POS + str_length(SEQ)-1),
      
      # FLAG = 16: mapped to the - strand, so the first character in the read array should be at position A
      FLAG == 16 ~ as.double(POS + 0)
    ))
}


# Function to create tibble aggregated by TA position
# The first argument is the sam file, the second argument is the TA position list
fx_TApos_mapping <- function(x_sam, y_TA){
  output <- y_TA %>% 
    right_join(x_sam, by = "TA_pos") %>% 
    group_by(TA_id) %>% 
    summarize(TA_reads = sum(read_count)) %>% 
    mutate(TA_RPM = TA_reads * 1000000 / sum(TA_reads))
}

# Function to execute all of each function
fx_all_function <- function(x, y){
  tmp1 <- fx_parseSAMfile(x)
  tmp2 <- fx_SAM4Tnseq(tmp1)
  tmp3 <- fx_TApos_mapping(tmp2,y)
}


# running ---------------------------------------------------------------

sam <- sample_sheet$sam
for (i in seq_along(sam)) {
  print(sam[[i]])
  sam_file <- str_c("3_mapping/", sample_sheet$sam[[i]])
  output_file = str_c(output_directory, 'A2_CountReads_', sample_sheet$sample_name[[i]], '.csv')
  output <- fx_all_function(sam_file, TA_pos_list)
  write_csv(output, output_file)
  print(str_c("finish ", sam[[i]]))
}
```
