## Supplementary Code for "A role for the S4-domain containing protein YlmH in ribosome-associated quality control in *Bacillus subtilis*": A3_calc_Fn.R

```
# input -------------------------------------------------------------------

input_sample_sheet <- 'sample_sheet2.csv'
input_TA_position <- 'reference/TA_position_168.csv'
input_TA_annotation <- 'reference/TA_annotation.csv'


# Package -----------------------------------------------------------------

library(tidyverse)

# parse sample sheet and TA position list------------------------------------------------------

sample_sheet <- read_csv(input_sample_sheet)
TA_position <- read_csv(input_TA_position)
TA_annotation <- read_csv(input_TA_annotation#, col_types = 'ccddddcccccccccccccccccccccccccccccccc'
)

cutoff_low_reads <- 8

# function ----------------------------------------------------------------


fx_Reads_per_TA <- function(x){
  tmp1 <- TA_position %>% 
    left_join(x) %>% 
    group_by(TA_start) %>% 
    summarize(TAb_RPM = sum(TA_RPM, na.rm = TRUE),
              TAb_reads = sum(TA_reads, na.rm = TRUE)) %>% 
    filter(TAb_reads >= cutoff_low_reads)
  tmp2 <- TA_annotation %>%
    left_join(tmp1, by = 'TA_start') %>%
    select(locus_tag, TA_start, TAb_reads, TAb_RPM) %>%
    distinct()
}

fx_AggGene <- function(x){
  output <- TA_annotation %>% 
    select(TA_start, locus_tag) %>% 
    distinct() %>% 
    left_join(x) %>%
    group_by(locus_tag) %>%
    summarize(Agg_fitness = mean(Fitness, na.rm = TRUE),
              TRPM_t1 = sum(TAb_RPM_t1, na.rm = TRUE),
              TRPM_t2 = sum(TAb_RPM_t2, na.rm = TRUE),
              FC_TRPM = TRPM_t2 / TRPM_t1,
              total_TA = n(),
              non_na_TA = sum(!is.na(TAb_RPM_t1)),
    )
}


fx_GeneInfo <- function(x){
  tmp1 <- x %>% 
    left_join(TA_annotation) %>% 
    mutate()
}


# processing ----------------------------------------------------------------

count_reads <- tibble(count_reads = fs::dir_ls('4_analysis/q20')) %>% 
  dplyr::filter(str_detect(count_reads, 'CountReads')) %>% 
  dplyr::mutate(sample_name = str_extract(count_reads, paste0(sample_sheet$sample_name, collapse = '|'))) %>% 
  print()


compare_list <- sample_sheet %>% 
  tidyr::pivot_wider(
    id_cols = c(cell_library, genotype),
    names_from = time_point,
    values_from = c(sample_name, inc_time, fold_change, sam)
  ) %>% 
  dplyr::select(-inc_time_t1, -fold_change_t1) %>% 
  dplyr::left_join(count_reads, by = c('sample_name_t1' = 'sample_name')) %>% 
  dplyr::rename(CountReads_t1 = count_reads) %>% 
  dplyr::left_join(count_reads, by = c('sample_name_t2' = 'sample_name')) %>% 
  dplyr::rename(CountReads_t2 = count_reads) %>% 
  print()


compare_set <- compare_list$cell_library

for (i in seq_along(compare_set)) {
  # Read t1 data frame
  CountReads_t1 <- str_c(compare_list$CountReads_t1[[i]])
  DF_t1 <- read_csv(CountReads_t1) 
  # Sum the reads of t1 for each TA_site
  Reads_per_TA_t1 <- fx_Reads_per_TA(DF_t1) %>% 
    rename(
      TAb_reads_t1 = TAb_reads,
      TAb_RPM_t1 = TAb_RPM
    )
  
  # Read t2 data frame
  CountReads_t2 <- str_c(compare_list$CountReads_t2[[i]])
  DF_t2 <- read_csv(CountReads_t2) 
  # Sum the reads of t2 for each TA_site
  Reads_per_TA_t2 <- fx_Reads_per_TA(DF_t2) %>% 
    rename(
      TAb_reads_t2 = TAb_reads,
      TAb_RPM_t2 = TAb_RPM
    )
  
  # Join t1 and t2 data frames
  EF = compare_list$fold_change_t2[[i]]
  output1 <- Reads_per_TA_t1 %>% 
    left_join(Reads_per_TA_t2, by = c('locus_tag'='locus_tag', 'TA_start'='TA_start')) %>% 
    
    # Calculate Fitness per TA site
    mutate(
      Fitness = case_when(
        !is.na(TAb_reads_t1) & !is.na(TAb_reads_t2) ~ log(EF * (TAb_RPM_t2/1000000) / (TAb_RPM_t1/1000000)) / log ( EF * (1-(TAb_RPM_t2/1000000)) / (1-(TAb_RPM_t1/1000000))),
        !is.na(TAb_reads_t1) & is.na(TAb_reads_t2) ~ 0,
        is.na(TAb_reads_t1) == 0  ~ NA_real_
      )
    )
  # export
  output_file1 = str_c("4_analysis/q20/A2_CalcFitness_", compare_list$cell_library[[i]], '.csv')
  write_csv(output1, output_file1)
  print(str_c("finish ", compare_set[[i]]))
}
```
