## Supplementary Code for "A role for the S4-domain containing protein YlmH in ribosome-associated quality control in *Bacillus subtilis*": A4_plotting.R

```
# input -------------------------------------------------------------------

input_gff <- 'reference/GCF_000009045.1_ASM904v1_genomic.gff'
input_cl1 <- '4_analysis/q20/A2_CalcFitness_cl1_wt.csv'
input_cl2 <- '4_analysis/q20/A2_CalcFitness_cl2_ssrA.csv'

# package -----------------------------------------------------------------

library(tidyverse)
library(rtracklayer)
library(patchwork)

# function ----------------------------------------------------------------


plot_TNseq <- function(x, y){
  target <-  x
  gene_selected <- gff %>% 
    filter(locus_tag == target) %>% 
    filter(type == 'gene')
  gene_selected
  gene_centre <- (gene_selected$start[1] + gene_selected$end[1]) / 2
  gene_left <- gene_selected$start[1]
  gene_right <- gene_selected$end[1]
  reg <- 1900
  
  p_A <- df_cl1 %>% 
    # filter(TA_start > gene_left - gene_expand - 500) %>% 
    # filter(TA_start < gene_right + gene_expand + 500) %>%
    filter(TA_start > gene_centre - reg - 500) %>%
    filter(TA_start < gene_centre + reg + 500) %>%
    ggplot() +
    geom_col(aes(TA_start, Fitness), width = 4, color = 'black')+
    #gene region
    geom_vline(aes(xintercept=gene_left), color="#FF4B00", alpha = 0.4) +
    geom_vline(aes(xintercept=gene_right), color="#FF4B00", alpha = 0.4) +
    ggtitle('wt') +
    coord_cartesian(
      xlim = c(gene_centre - reg, gene_centre + reg),
      ylim = c(0, 1.5)
    )
  
  
  
  p_B <- df_cl2 %>% 
    # filter(TA_start > gene_left - gene_expand - 500) %>% 
    # filter(TA_start < gene_right + gene_expand + 500) %>%
    filter(TA_start > gene_centre - reg - 500) %>%
    filter(TA_start < gene_centre + reg + 500) %>%
    ggplot() +
    geom_col(aes(TA_start, Fitness), width = 4, color = 'black')+
    #gene region
    geom_vline(aes(xintercept=gene_left), color="#DE2725", alpha = 0.4) +
    geom_vline(aes(xintercept=gene_right), color="#DE2725", alpha = 0.4) +
    ggtitle('ssrA') +
    coord_cartesian(
      xlim = c(gene_centre - reg, gene_centre + reg),
      ylim = c(0, 1.5)
    )
  
  
  p_C <- gff %>% 
    mutate(gene_init = if_else(strand == '+', start, end)) %>% 
    filter(type == 'gene') %>% 
    filter(start > gene_centre - reg - 5000) %>%
    filter(end   < gene_centre + reg + 5000) %>%
    ggplot() +
    geom_rect(aes(xmin = start, xmax = end, ymin = 0, ymax = 1), fill = 'grey80', color = 'grey40') +
    geom_vline(aes(xintercept = gene_init), color='black', size = 2) +
    geom_text(aes(x=(start + end)/2, y=0.5, label = gene)) +
    coord_cartesian(xlim = c(gene_centre - reg, gene_centre + reg)) +
    theme_void()
  
  p_A / p_B / p_C + plot_annotation(
    title = as.character(x)
  )
}


# loading -----------------------------------------------------------------

gff <- rtracklayer::readGFF(input_gff) %>% 
  as_tibble()

df_cl1 <- read_csv(input_cl1)
df_cl2 <- read_csv(input_cl2)


# plotting ----------------------------------------------------------------

# ssrA, BSU_misc_RNA_55
pdf('figure_1A/q20_a_ssrA.pdf', width = 9.9, height = 4.5)
print(plot_TNseq('BSU_misc_RNA_55'))
dev.off()

# brfA, BSU_23540
pdf('figure_1A/q20_b_brfA.pdf', width = 9.9, height = 4.5)
print(plot_TNseq('BSU_23540'))  # ssrA, BSU_misc_RNA_55)
dev.off()

# rqcH, BSU_15640
pdf('figure_1A/q20_c_rqcH.pdf', width = 9.9, height = 4.5)
print(plot_TNseq('BSU_15640'))  # ssrA, BSU_misc_RNA_55)
dev.off()

# rqcP, BSU_00590
pdf('figure_1A/q20_d_rqcP.pdf', width = 9.9, height = 4.5)
print(plot_TNseq('BSU_00590'))  
dev.off()

# ylmH, BSU_15410
pdf('figure_1A/q20_d_ylmH.pdf', width = 9.9, height = 4.5)
print(plot_TNseq('BSU_15410'))  
dev.off()
```
