## Supplementary Code for "A role for the S4-domain containing protein YlmH in ribosome-associated quality control in *Bacillus subtilis*": codes.md

# Codes for fastq processing

## update
(2023/09/19) p2, 3'のTn配列でsplitするところで、len19のミスマッチ許容を他と同じにした  
(2023/09/13) fastxのbarcode splitterを使っていたが、seqkitに置き換えた  

## raw reads
- 210924_NB501353_0170_AHLLNGBGXF/Undetermined_S0_L00*_R1_001.fastq.gz  
(renamed: Undetermined_S0_L00*_R1_001_210924.fastq.gz)  
- 211026_NB501353_0173_AHLL7LBGXF/Undetermined_S0_L00*_R1_001.fastq.gz  
(renamed: Undetermined_S0_L00*_R1_001_211026.fastq.gz)  


## prepare files
- set working directory
- make folder named "1_fastq/0_raw" in working directory and put raw files into the folder
- prep 3' barcode sequences derived from transposon sequences
- prep 5' index sequences for demultiplexing samples
- reference genome sequence (.fna)

## Required
- SeqKit (2.5.1)
- R (v4.3.0)
- tidyverse (v2.0.0)
- collapse_reads.R


## [Pre] check fastq quality
fastqcでfastqのクオリティを確認。大丈夫そうなら次へ。  
### required
- fastqc

### code 
```
fastqc /cont_wor_dir/*.fastq.gz -o /cont_wor_dir/output_fastqc
```

## [P1] 1st Trimming
readの5'と3'を大まかにトリミングし、必要な配列部分（10-45残基目）だけを抽出する。
### required
- seqkit

### code
```
seqkit subseq -r 10:45 ${file} > ./data/p1_${filename%%.*}.fastq
```


## [P2] Split by 3'-tail
MmeI認識配列から切断サイト（= アダプターのライゲーション位置）までの距離が不均一で、readによって異なる。それぞれから必要な配列部分（染色体由来部分）だけを抽出するため、ここではまず、前の過程で得た均一な長さのreadのうち、3'側の配列（ゲノム上のTAの位置からトランスポゾンの配列あたりにかけて）で仕分ける。  
- L16やL17 (MAGenTAでいうところのTrim2, 3) が大部分  
- L18やL19 はほとんどないので無視してもいいのかもしれない？　　

(KF調べ)


### required
- seqkit

### code
```
for file in 1_fastq/p1*.fastq.gz; do
for INDX in 1_fastq/barcodes/BC3_ADBC_L*.txt; do
    echo ${file}
    filename=${file##*/p1_Undetermined_S0_}
    index_seq=`cat ${INDX}`
    index_file=${INDX##*/}
    index_name=${index_file%.*}
    echo ${index_name} ${index_seq}

    seqkit grep ${file} \
    -i \
    -s \
    -P \
    -p ${index_seq} \
    -R -6:-1 \
    -m 0 \
    -o 1_fastq/p2_${index_name}_${filename%%.*}.fastq.gz


done
done
```


## [P3] 2st Trimming (3')
3'末端にあるトランスポゾン由来の配列部分を除去し、TAの配列が３’末端に来るようにする。染色体由来配列の長さに応じて変える必要がある。  
処理後、できればこの段階で各lane毎にconcatenateする
### required
- seqkit

### code  
```
# for trimming
for file in 1_fastq/p2_BC3_ADBC_L15*.fastq.gz; do
    echo ${file}
    filename=${file##*/p2_}
    seqkit subseq -r 1:-16 ${file} > 1_fastq/p3_${filename%%.*}.fastq.gz
done

for file in 1_fastq/p2_BC3_ADBC_L16*.fastq.gz; do
    echo ${file}
    filename=${file##*/p2_}
    seqkit subseq -r 1:-15 ${file} > 1_fastq/p3_${filename%%.*}.fastq.gz
done
... etc

# for concatenating
cat 1_fastq/p3*L001* > 1_fastq/p3c_L001.fastq.gz
cat 1_fastq/p3*L002* > 1_fastq/p3c_L002.fastq.gz
... etc

```


## [P4] Split by 5' barcode
DNAライブラリ作成時のアダプターライゲーションの段階で、サンプル毎にバーコードが割り当てられている。そのバーコード毎にreadを仕分ける。複数レーンのファイルがあるのであれば、一旦バラバラでsplitし、後からconcatenateする。

### required
- seqkit
- fastx_barcode_splitter.pl

### code
```
for file in 1_fastq/p3c*.fastq.gz; do
    for INDX in 1_fastq/barcodes/INDX*.txt; do
        echo ${file}
        filename=${file##*/p3c_}
        index_seq=`cat ${INDX}`
        index_file=${INDX##*/}
        index_name=${index_file%.*}
        echo ${index_name} ${index_seq}

        seqkit grep ${file} \
        -i \
        -s \
        -P \
        -p ${index_seq} \
        -R 1:6 \
        -m 0 \
        -o 1_fastq/p4_${index_name}_${filename%%.*}.fastq.gz
    done
done

echo "concatenating by INDX"
for INDX in 1_fastq/barcodes/INDX*.txt; do
    index_file=${INDX##*/}
    index_name=${index_file%.*}
    echo ${index_name}
    cat 1_fastq/p4*${index_name}* > 1_fastq/p4_${index_name}_merged.fastq.gz
done

echo "finished"
```


## [P5] check duplicate
### required
- seqkit  


### code
```
seqkit rmdup -n -i ${file} -o /cwd/data/under_processing/p5_clean_${filename%%.*}.fastq -d /cwd/data/under_processing/p5_duplicated_${filename%%.*}.fastq -D /cwd/data/under_processing/p5_duplicate_detail_${filename%%.*}.txt

#duplicateのチェックはp7の前後（同時？）の方が効率的かもしれない？でもそんなに変わらないかも？
```


## [P6] 3rd trimming (5')
5'側にあるバーコード領域をトリミングする。これで、readは染色体由来配列部分のみになる。
### required
-  seqkit
### code
```
seqkit subseq -r 7:-1 ${file} > /cwd/data/under_processing/p6_${filename%%.*}.fastq
```


## [p7] filtering by quality
シーケンスクオリティの低いreadをここで除去する。  
基準: 8  
ref:  MAGenTA  
クオリティスコアが8でカット。これはかなり緩めの設定らしい。  
（23911時点）もしかしたら、これまでの解析で、間違えてフィルタリングしているケースがあるかもしれない。というよりフィルタリングできていないことがあるかも。ただ、フィルタリングはそもそもほとんどいらないらしいので（MAGenTAマニュアル参照）、結果に大きな影響はないはず。再解析することでもしかしたらもっと綺麗なデータになるかもしれない？  

### required 
- FASTX-toolkit
- or seqkit
### code
```
# FASTX-toolkitの場合
# 平均クオリティスコアが8以上のリードのみを抽出
fastq_quality_filter -q 8 -i ${file} -o /cwd/data/under_processing/p7_${filename%%.*}.fastq

# seqkitの場合
# 平均クオリティスコアが8以上のリードのみを抽出
seqkit seq -Q 8 ${file} > /cwd/data/under_processing/p7_${filename%%.*}.fastq

```


## [p8] collapse reads
マッピング効率を上げるため、複数ある同一readを1 readにする。配列名にread数を入れることで、同一リードのカウント情報を含める。  
~~FASTX-toolkit以外のモダンソフトでこれができたら？さがす~~  
~~seqkitのrmdupでうまいことできない？~~  
そもそもRでできた。
### required
- R (v4.3.0)
- tidyverse (v2.0.0)
- collapse_reads.R
```
Rscript collapse_reads.R
```
