## Supplementary Code for "A role for the S4-domain containing protein YlmH in ribosome-associated quality control in *Bacillus subtilis*": collapse_reads.R

```
# required packages:
## tidyverse
## ShortRead

# input -------------------------------------------------------------------

args <- commandArgs(trailingOnly = T)
input <- args[1]

# package -----------------------------------------------------------------

library(tidyverse)
library(ShortRead)

# prep file list ----------------------------------------------------------


# formatting --------------------------------------------------------------

output = str_remove(input, '.gz') %>% str_replace('p6', 'p7')

tmp1 <- input %>% 
  ShortRead::readFastq() %>% 
  ShortRead::sread() %>% 
  as.character() %>% 
  dplyr::as_tibble() %>% 
  dplyr::count(value) %>% 
  dplyr::arrange(desc(n), value) %>% 
  dplyr::mutate(name = str_c(row_number(), '-', n)) %>% 
  dplyr::select(name, seq = value)

tmp2 <- DNAStringSet(tmp1$seq)

names(tmp2) <- tmp1$name


# output ------------------------------------------------------------------

tmp2 %>% 
  ShortRead::writeFasta(
    file = output
  )

print('finished')
```
