## Supplementary Code for "A role for the S4-domain containing protein YlmH in ribosome-associated quality control in *Bacillus subtilis*": p1_trimming.sh

```
#!/bin/bash

for file in 1_fastq/0_raw/*.fastq.gz; do
    echo ${file}
    filename=${file##*/}
    #echo p1_${filename%%.*}.fastq
    seqkit subseq -r 10:45 ${file} > 1_fastq/p1_${filename%%.*}.fastq.gz
done
```
