## Supplementary Code for "A role for the S4-domain containing protein YlmH in ribosome-associated quality control in *Bacillus subtilis*": p3_trim_tail.sh

```
#!/bin/bash

for file in 1_fastq/p2_BC3_ADBC_L15*.fastq.gz; do
    echo ${file}
    filename=${file##*/p2_}
    seqkit subseq -r 1:-16 ${file} > 1_fastq/p3_${filename%%.*}.fastq.gz
done

for file in 1_fastq/p2_BC3_ADBC_L16*.fastq.gz; do
    echo ${file}
    filename=${file##*/p2_}
    seqkit subseq -r 1:-15 ${file} > 1_fastq/p3_${filename%%.*}.fastq.gz
done

for file in 1_fastq/p2_BC3_ADBC_L17*.fastq.gz; do
    echo ${file}
    filename=${file##*/p2_}
    seqkit subseq -r 1:-14 ${file} > 1_fastq/p3_${filename%%.*}.fastq.gz
done

for file in 1_fastq/p2_BC3_ADBC_L18*.fastq.gz; do
    echo ${file}
    filename=${file##*/p2_}
    seqkit subseq -r 1:-13 ${file} > 1_fastq/p3_${filename%%.*}.fastq.gz
done

for file in 1_fastq/p2_BC3_ADBC_L19*.fastq.gz; do
    echo ${file}
    filename=${file##*/p2_}
    seqkit subseq -r 1:-12 ${file} > 1_fastq/p3_${filename%%.*}.fastq.gz
done


echo "concatenating L001"
cat 1_fastq/p3_*L001* > 1_fastq/p3c_L001.fastq.gz

echo "concatenating L002"
cat 1_fastq/p3_*L002* > 1_fastq/p3c_L002.fastq.gz

echo "concatenating L003"
cat 1_fastq/p3_*L003* > 1_fastq/p3c_L003.fastq.gz

echo "concatenating L004"
cat 1_fastq/p3_*L004* > 1_fastq/p3c_L004.fastq.gz


echo "finished"
```
