## Supplementary Code for "A role for the S4-domain containing protein YlmH in ribosome-associated quality control in *Bacillus subtilis*": p5_trim_5.sh

```
#!/bin/bash

for file in 2_fastq_demultiplexed/s*.fastq.gz; do
    echo ${file}
    filename=${file##*/}
    seqkit subseq -r 7:-1 ${file} > 2_fastq_demultiplexed/${filename%%.*}_p5.fastq.gz
done

echo "finished"
```
