## Supplementary Code for "A role for the S4-domain containing protein YlmH in ribosome-associated quality control in *Bacillus subtilis*": p6_filter8.sh

```
#!/bin/bash
# filtering

for file in 2_fastq_demultiplexed/*p5.fastq.gz; do
    echo ${file}
    filename=${file##*/}
    seqkit seq -Q 8 ${file} > 2_fastq_demultiplexed/${filename%%_p5.*}_p6.fastq.gz
done

echo "finished"
```
