## Supplementary Code for "A role for the S4-domain containing protein YlmH in ribosome-associated quality control in *Bacillus subtilis*": p7_collapse_reads.sh

```
#!/bin/bash
# collapse into a virtual single read 

for file in 2_fastq_demultiplexed/*_p6.fastq.gz; do
    echo ${file}
    filename=${file##*/}
    Rscript collapse_reads.R ${file}
done

echo "finished"
```
