## Supplementary Code for "A role for the S4-domain containing protein YlmH in ribosome-associated quality control in *Bacillus subtilis*": p8_bowtie_mapping.sh

```
#!/bin/bash
# mapping by bowtie to 168 genome sequence

echo "mkdir"
mkdir -p 3_mapping/

for file in 2_fastq_demultiplexed/*_p7.fasta; do
    filename=${file##*/}
    echo "start $filename"
    bowtie -S -n 1 --best -y -m 1 -p 6 \
    -x 3_mapping/Bowtie_Index_168/bt1IDX_Bs168 \
    -f ${file} 3_mapping/${filename%%.*}.sam
done

# -S/--sam           write hits in SAM format
# -n/--seedmms <int> max mismatches in seed (can be 0-3, default: -n 2)
# --best             hits guaranteed best stratum; ties broken by quality
# -y/--tryhard       try hard to find valid alignments, at the expense of speed
# -m <int>           suppress all alignments if > <int> exist (def: no limit)
# -p/--threads <int> number of alignment threads to launch (default: 1)
# -x <ebwt>          <ebwt>  Index filename prefix (minus trailing .X.ebwt).
# -f                 query input files are (multi-)FASTA .fa/.mfa
```
